# Towards a Chimeric Vaccine against Multiple Isolates of *Mycobacteroides* - An Integrative Approach

**DOI:** 10.1101/2019.12.15.869081

**Authors:** Rohit Satyam, Tulika Bhardwaj, Niraj Kumar Jha

**Author notes:** Correspondence, Dr. Niraj Kumar Jha.

## Abstract

Nontuberculous mycobacterial infection (NTM) such as endophthalmitis, dacryocystitis, canaliculitis are pervasive across the globe and are currently managed by antibiotics such as cefoxitin/imipenem and azithromycin/clarithromycin. However, the recent cases of Mycobacteroides developing drug resistance reported along with the improper practice of medicine intrigued us to explore its genomic and proteomic canvas at a global scale. A timely developed vaccine against Mycobacteroides is, therefore, a much requirement. Consequently, we carried out a vivid Genomic study on five recently sequenced strains of Mycobacteroides and explored their Pan-Core genome/ proteome. The promiscuous antigenic proteins were identified via a subtractive proteomics approach that qualified for virulence causation, resistance and essentiality factors for this notorious bacterium. An integrated pipeline was developed for the identification of B Cell, MHC class I, II epitopes. Our final vaccine construct, V6 qualified for all tests such as absence for allergenicity, presence of antigenicity, etc. and contains β defensin adjuvant, linkers, LAMP1 signal peptide, and PADRE (Pan HLA-DR epitopes) amino acid sequence. The vaccine construct, V6 also interacts with a maximum number of MHC molecules, and the TLR4/MD2 complex confirmed by docking and molecular dynamics simulation studies. The knowledge harnessed from the current study can help improve the current treatment regimens and propel further related studies.

## 1. Introduction

Mycobacteroides gen. nov (Taxonomy ID: 670516) is a new genus formed according to the new classification made effective in the year 2018, distinguished by 51 unique molecular markers (Conserved Signature Indels CSIs and Conserved Signature Proteins CSPs) and constitute fast-growing members of *Abscessus-Chelonae clade*. **[1]** The members of this genus form a deep and branching monophyletic clade within the Mycobacteriaceae family. The comprising members of *Abscessus-Chelonaeclade*arenon-sporulating, acid-fast, Gram-positive bacillus with distinguishing phenotypic traits such as optimum growth at 30^0^ C, resistance to polymyxin B, positive arylsulfatase test (require 3 days), and negative iron uptake and nitrate reductase test. **[2]** Members of this genus are referred to as Rapidly Growing Mycobacteroides (RGMs) and are responsible for causing diseases related to Pulmonary, Cutaneous **[3]**, Musculoskeletal diseases and other body parts **[4]**. The bacterium exhibit pathogenesis in human and are developing resistance against antimicrobial therapeutics **[5, 6, 7]**.

*Mycobacterium chelonae* is a Non-TuberculousMycobacteroide (NTM) and a Gram-positive bacillus. The strain belonged to Group IV of Runyon Classification and are non-pigmented, non-motile, acid-fast staining bacterium **[8]** that along with *M. abscessus species* forms the most drug-resistant members **[22]**. The difference at the genomic level also influences the therapy and treatment **[22]**. *Chelonae* strains respond better to the drugs as compared to *Abscessus* strains due to lack of *erm gene* that is responsible for macrolide-resistance in *M. abscessus* **[24]**. Unlike *Abscessus, Chelonae* strains are inhibited at a higher concentration of cefoxitin but are susceptible to tobramycin **[22]**. In a recent review **[9]**, both *Chelonae* and subsp. *bolletii* strains and are known to cause Non-Tuberculosis Infection (NTI) post to renal transplant; disseminated disease being the most prevalent manifestation (40%) and *M. chelonae* being the most common pathogen. Organ transplants, malignancy, diabetes mellitus, tumor necrosis factor-alpha (TNF-α) inhibitors, immunosuppressant therapy, and long-term corticosteroid administration causes the Disseminated disease **[19, 25]**. *M. chelonae* species are found to thrive even in chlorinated water and in industrial-grade detergents such as glutaraldehyde **[10]**, freshwater sources, soil, contaminated solutions, distilled water, and potable water. The optimum condition of growth for the strains is documented at 30-32°C and requires a prolonged incubation period. The global epidemiology demarcates the presence of this *Mycobacteroides* on global canvas ranging from America, Europe, and Asia to Australia, with increased cases in southern and coastal states of Texas. Louisiana, Georgia, Florida **[2, 22]**. Pulmonary infections are however rarely caused by *Chelonae***[19]**.

The pathophysiology of *Chelonae* strains include NTM keratitis, a rare complication of keratomileusis [LASIK], and conjunctivitis, (since eyes are the second most frequent organ affected by *Chelonae)* endophthalmitis, dacryocystitis, canaliculitis, scleritis, **[11, 12]**, cutaneous infections **[13, 15]**, renal abscess **[14]**, catheter and surgical related infections after transplants, scleropathy and implants such as stent **[16]**. Impaired immune system, accidental trauma, tattoo inks **[17]**, mesotherapy and acupuncture **[18]** are well known reason that facilitate infection often characterized by bacteremia (with or without chills, fever and sweat) **[19]**, disseminated cutaneous infestation, osteomyelitis, intra-abdominal abscess, localized cellulitis, bloodstream infection **[20]** and chronic kidney diseases. When grown over high nutrient media, both *Abscessus* and *Chelonae*produce extensive cording colonies **[21]**. The *M. chelonae* strains are currently being treated clinically with drugs such as imipenem (60%), amikacin (50%) and highly susceptible to Tobramycin (100%), clarithromycin (100%), linezolid (90%), clofazimine, doxycycline (25%), and ciprofloxacin (20%) **[22]**. Nonetheless, cases of antimicrobial resistance have surfaced recently due to a single point mutation at position 2058 of 23S rRNA. **[23]**

*M. abscessus*, on the other hand, is the most notorious and common mycobacteroide causing pulmonary disease and the drug therapy of this NTM is cost-intensive, exhaustive and sometimes lead to drug toxicities **[5, 19, 26, 27, 28]**.Both subsp. *Abscessus* and *Bolletii* have a difference in clinical symptomatic presentation and Drug Sensitivity Test (DST) and were found to respond differently to antibiotic treatment **[30, 31]**. The drug susceptibility tests carried out elsewhere **[29]** as per CLSI guidelines indicate that *subsp. Bolletii* exhibited distinguishing characteristics from *subsp Abscessus. Bolletii* strains were found to form rough colonies in major proportion in contrast to *subsp. Abscessus* where rough and smooth colonies were formed in equal proportion and were found to be levofloxacin-resistance in addition to Clarithromycin. However, it is susceptible to azithromycin, amikacin, linezolid, and imipenem significantly **[29]**. ATS/IDSA (American Thoracic Society/Infectious Diseases Society of America) in 2007 released guidelines for treatment *M. abscessus* infection via intravenous administration of amikacin and cefoxitin/imipenem in conjugation with azithromycin/clarithromycin, for 2–4 months **[30]**.

The treatment regimens against Mycobacteroides infection vary globally and so does the pattern of resistance due to drug abuse. The growing antimicrobial resistance recently has raised public health concerns **[32, 33]**. *In-vivo* experiments reveal that bronchiectasis in the Japanese population is frequently due to infection with *subsp. abscessus* than with *subsp. bolletii* and imipenem-resistance was higher in *M. abscessus subsp. bolletii* comparatively. Similar kinds of results were also reported elsewhere **[30]**. However, resistance and drug susceptibility are different in China. Due to the overexposure of fluoroquinolones, the Chinese population exhibits levofloxacin-resistance in NTM patients but still susceptible to other drugs such as imipenem significantly. Similarly, when both levofloxacin and moxifloxacin are fluoroquinolones, resistance to levofloxacin was much higher comparatively **[29]**. Studies also suggest the prolonged incubation of *subsp. Abscessus* with clarithromycin induces clarithromycin-resistance making it inefficient for treatment. However, *subsp. bolletii* do not exhibit an inducible resistance phenomenon **[24]**.

The present study tries to understand the pathogenesis of the above-mentioned Mycobacteroides and address the issue of escalating the antimicrobial resistance in the *Bolletii-Chelonae* strains whose complete genomes were available at the time of writing this paper. We carried out literature mining to construct a novel and robust pipeline amalgamating the Pan-Genome analysis, Subtractive Proteomics, Immunoinformatics, and reverse vaccinology approach to identify putative vaccine targets that could eventually lead to the construction of thermodynamically stable chimeric vaccine. Given to paucity of information of these completely sequenced strains of Mycobacteroides, we decided to carry out extensive genomic and proteomic analysis in multiple Phases (I, II, & III). We translated the core genome to core proteome and then used the Subtractive Proteomics approach to identify Non-Homologues against Human Proteome. Once shortlisted, the putative vaccine target proteins can act as promising candidates to construct a chimeric gene that can be overexpressed in *E. coli* with subsequent extraction and purification by affinity chromatography/ Size Exchange Chromatography (SEC), respectively **[34]**. The multi-epitope subunit vaccine then can be used to activate Cell-mediated and Humoral immunity of the immunocompromised host.

## 2. Materials and Methodology

In the present study, we integrated different approaches to understand both proteomic and the genomic repertoire of Mycobacteroides. In Phase I, we explored the genomic landscape of the organism and performed Pan-genome analysis. The protein dataset so obtained was subjected to a subtractive proteomic approach in Phase II. The shortlisted candidates from Phase II were then used in Phase III to develop a thermodynamically stable multi-epitope vaccine. (Figure 1).

**Figure 1:**
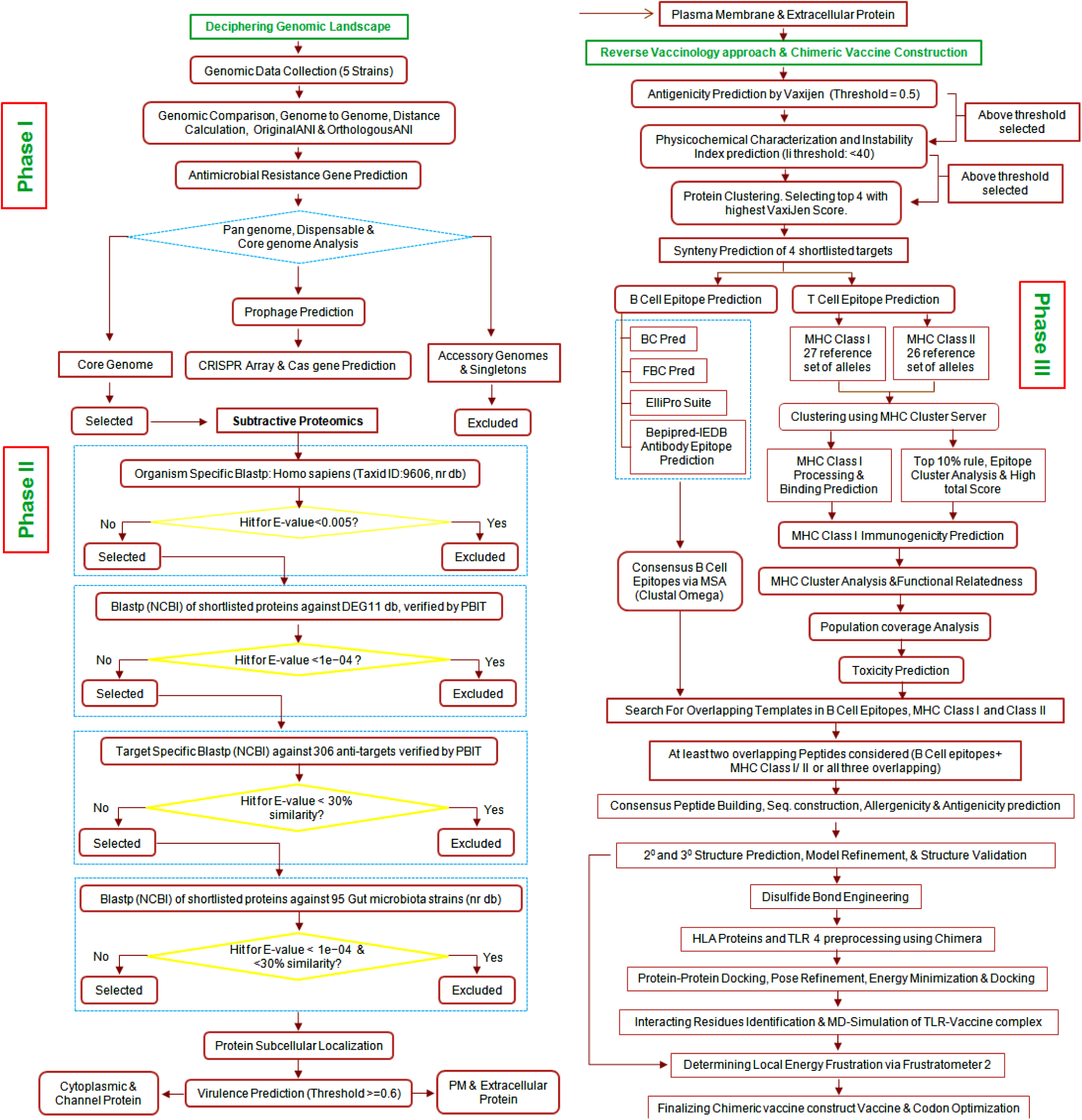
Flowchart of sequential analysis Performed: This is the complete workflow of the present study that guides the Chimeric Vaccine Construction and Drug Target Identification. The study was divided into 4 Phases; Phase I: Deciphering Genomic Landscape, Phase II: Subtractive Proteomics Approach, Phase III: Reverse Vaccinology approach & Chimeric Vaccine Construction, Phase IV: Drug Target Identification.

### Phase I: Deciphering the Genomic Landscape

#### Genomic Data Collection and Extraction

In the present study, we retrieved complete genome sequences of strains of Mycobacteroides from the genome browser of NCBI both in GenBank (gbk) format and FASTA (fas/ fasta) format. At the time of writing, there are 90 complete genomes of Mycobacteroides present at the NCBI database(https://www.ncbi.nlm.nih.gov/genomes/GenomesGroup.cgi?taxid=670516). However, we decided to limit our study to five recently sequenced strains to unravel features of these new members (*Mycobacterium abscessus subsp. bolletii* strain MM1513, *Mycobacterium abscessus subsp. bolletii* strain MC1518, *Mycobacterium abscessus subsp. bolletii* strain MA 1948, *Mycobacteroides chelonae* strain MOTT36W, *Mycobacterium chelonae* CCUG 47445).

#### Genomic Comparison

The whole-genome sequences of five strains of Mycobacteroides were aligned using the progressiveMauve mode of Mauve (release: 2015-02-26) **[35, 36]** and the alignment statistics were generated and visualized. To analyze distinctness at the genomic level, Genome-to-Genome distance and Average nucleotide identity (ANI) percentage were calculated using OrthoANI Tool v 0.93.1 **[37]**(https://www.ezbiocloud.net/tools/orthoani). The OrthoANI is a new approach developed to decipher genomic relatedness of Bacterial and Archaeal species by fragmenting the genomes and subsequently using the orthologous fragments for nucleotide identity calculation **[37]**. The heatmap was generated and visualized to demarcate the overall genomic relatedness. We used Artemis of Sanger Institute to probe into the genomes of 5 Mycobacteroides and analyze GC content, CDS and presence of Pseudogenes **[44]**. Tandem Repeat Finder (version 4.09) Benson (1999) was used to identify the number of tandem repeats present in the respective genome **[45]**.

#### Antimicrobial Resistance Gene Prediction

To identify if the genomes of five strains were populated with antimicrobial genes we used ResFinder v3.1 **[38]**. We queried all the antimicrobial classes mentioned at the server with percent identity set at 90% and the minimum query cover length of 80%. The genes so obtained were mapped against the previously aligned sequence to identify and visualize the genomic stretch and confirm the gene presence across strains.

#### Pan-genome, Dispensable, and Core genome Analysis

The Pangenome analysis would pool out the genes and eventually proteome that is common to all five strains. Therefore, the BPGA pipeline was deployed **[39]** that carry out genome clustering using a suitable algorithm and then carrying out Pan Genome analysis over the data generated. The sequences were input in GenBank format. We uploaded gbk files in the following order: *Mycobacterium abscessus subsp. bolletii* strain MA 1948.gb, *Mycobacterium abscessus subsp. bolletii* strain MC1518.gb, *Mycobacterium abscessus subsp. bolletii* strain MM1513.gb, *Mycobacterium chelonae* CCUG 47445.gb, *Mycobacteroides chelonae* strain MOTT36W.gb. with strain 1918 taken as a reference. We ran the USEARCH algorithm for genomic clustering with an identity cutoff of 50%. The BPGA provides an option to utilize the clustering data to perform a spectrum of analysis in addition to Pan-Genome Analysis. Therefore, atypical GC Content analysis was performed with an extreme GC content cutoff of 5%. The pan and core phylogenetic trees were constructed using the Neighbor Joining Method. KEGG/COG Functional analysis was performed to group the core, accessory, and Unique genomes into various functional categories.

#### ProPhage Prediction

The presence of Prophage sequences in the bacterial genome helps us understand how the bacterial genome evolves. In our study, we deployed PHASTER server (phaster.ca), an improved and relatively fast version of PHAST for identification of prophage sequences in the whole genomes of bacterial strains using genome Accession ID **[40]**. PHASTER queries Virus & Prophage Database and Bacterial database to identify potent prophage sequences and ranks hits based on a score as Intact (score > 90), Questionable (score 70-90) and Incomplete (score < 70). The hits were visualized in PHASTER Genome Viewer **[40]**

#### CRISPR Array and Cas gene prediction

We carried out CRISPR related Cas gene search in five strains of Mycobacteroides using CRISPERCasFinder **[41]** and CRISPRDetect **[43]**. The Evidence Level **[42]** and the Score **[43]**, respectively, were checked after prediction to distinguish the spurious predicted CRISPR array and cas genes (false positives) from the true CRISPR elements.

### Phase II: Subtractive Proteomics Approach

#### Non-Homology Analysis

The core proteome that was obtained after Pan-Core Analysis was subjected to a Non-Homology search against *Homo sapiens* (taxid: 9606). This step is critical to reducing the off-target effect and undesirable cross-reactivity of vaccine/drug in humans **[49]**. We used a Nonredundant database of Human proteome to screen bacterial core proteome using a BLASTp search with an expected threshold value set at 0.005 **[50]**. Proteins that were found non-homologous to human proteome repertoire were retained while the homologoues were segregated separately. The non-homology BLASTp was carried out in triplicate, taking 10,000 proteins each time, to remove false positives.

#### Essentiality Analysis

The sequence similarity search with an E value of 0.0001 was carried out against the DEG database (http://www.essentialgene.org/) **[51]**. The DEG database contains more than 22,343 essential genes, regulatory sequences, origins of replications (Ori) from both Eukaryotes and Prokaryotes which are experimentally identified. We performed a BLAST search of Unannotated Whole genomes of all 5 Mycobacteroides strains against DEG with cut-off parameters of 1e^-04^ E-value, bit score of 100, BLOSUM62 matrix and gapped alignment mode **[50]**. Protein-coding genes and protein alignment hits lying below the threshold were considered significant. The PBIT Pipeline was also used that combinatorically search DEG and VFDB to identify putative essential proteins **[52]**. E-value > 0.0001 and sequence identity < 30% was chosen as the criteria for Non-Homology. The proteins that were in alignment with the PBIT Pipeline were finally considered Essential.

#### Human Anti-target Non-Homology screening

Human anti-target in Pharmacology refers to the receptor proteins, enzymes or any other biological entity inside human which aren’t destined to be deliberately targeted by drug compounds designed, however, when binds cause side-effects. Such essential human (host) proteins are termed as ‘anti-targets’. The most significant one includes the human ether-a-go-go-related gene (hERG), 5-HT_2B_ receptor (Serotonin receptor 2B), constitutive androstane receptor (CAR), the pregnane X receptor (PXR), and P-glycoprotein (P-gp). Some membrane receptors falling in this Off-Target list includes dopaminergic D2, adrenergic a1a, serotonergic 5-HT_2C_, and the muscarinic M1 **[53]**. The GenInfo Identifiers or GI Numbers of the anti-targets were extracted from the literature the list of which is given in the List 3S, Supplementary section. We used 306 shortlisted anti-targets (after removal of obsolete/repeated proteins) to screen the non-homologous proteins by using BLASTp with 0.005 as an e-value threshold.

#### Gut Flora non-homology analysis

The proteins delineated from the previous step were assessed for homology with the proteome of human gut microbiota. Around 300 to 500 bacterial species are reported to populate our GI Tract with approximately 10 times in numbers of all the cells in our body **[54]**. Gut-commensal flora interactions play a vital role in metabolism by fermenting indigestible food particles and promoting homeostatic functions such as upregulation of cytoprotective genes, immunomodulation, maintenance of barrier function, prevention and regulation of apoptosis, etc**[55]**. Indeliberate blocking of the proteins essential for this menagerie of microbes might inadvertently slow their growth rate **[54]**. The proteins screened from the previous step were therefore subjected to a homology search against proteome of 95 essential Gut bacteria to avoid such circumstances using BLASTp with 0.0001 as an e-value threshold. The Gut microbiota used for this analysis has been enlisted in List 5S, Supplementary section.

#### Protein Sub-Cellular Localization

Mycobacteroides are Gram-positive with an outer membrane. Though proteins can be broadly categorized into 5 subcellular location i.e. extracellular proteins, outer membrane, plasma membrane, cytoplasm, and periplasm **[56]**; we chose to categorize them broadly into 3 groups Cytoplasmic, Plasma Membrane and Extracellular to segregate putative vaccine candidates. We used consensual subcellular prediction where we used multiple prediction servers to reach to a consensus subcellular location for a protein. The protein, to belong to the Cytoplasmic region (say), must be predicted cytoplasmic by two or more prediction methods. We used BUSCA (Bologna Unified Subcellular Component Annotator) **[57]**, CELLO v.2.5 **[127]**, PSORTb v 3.0.2 **[128]**, LocateP v2 **[59]** and BLAST2GO for Subcellular Localization (SCL) **[58]**. The subcellular pipeline adopted in BLAST2GO is shown in Fig 3S Supplementary Section.

**Figure 2:**
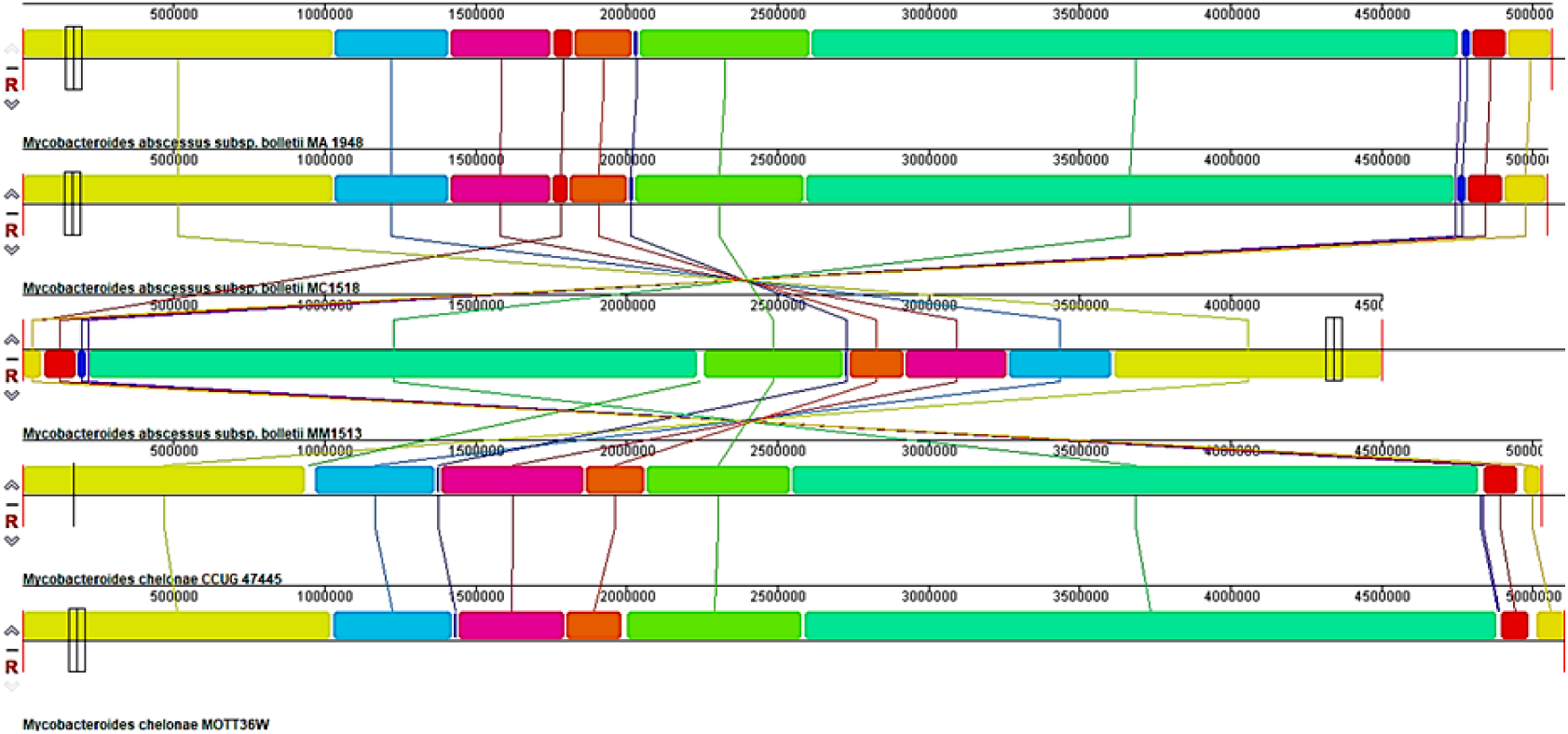
Multiple genome alignment of 5 Mycobacteroides strains using progressive MAUVE (From top to bottom); Strain MA 1948, MC1518, MM1513, CCUG47445, MOTT36W

**Figure 3:**
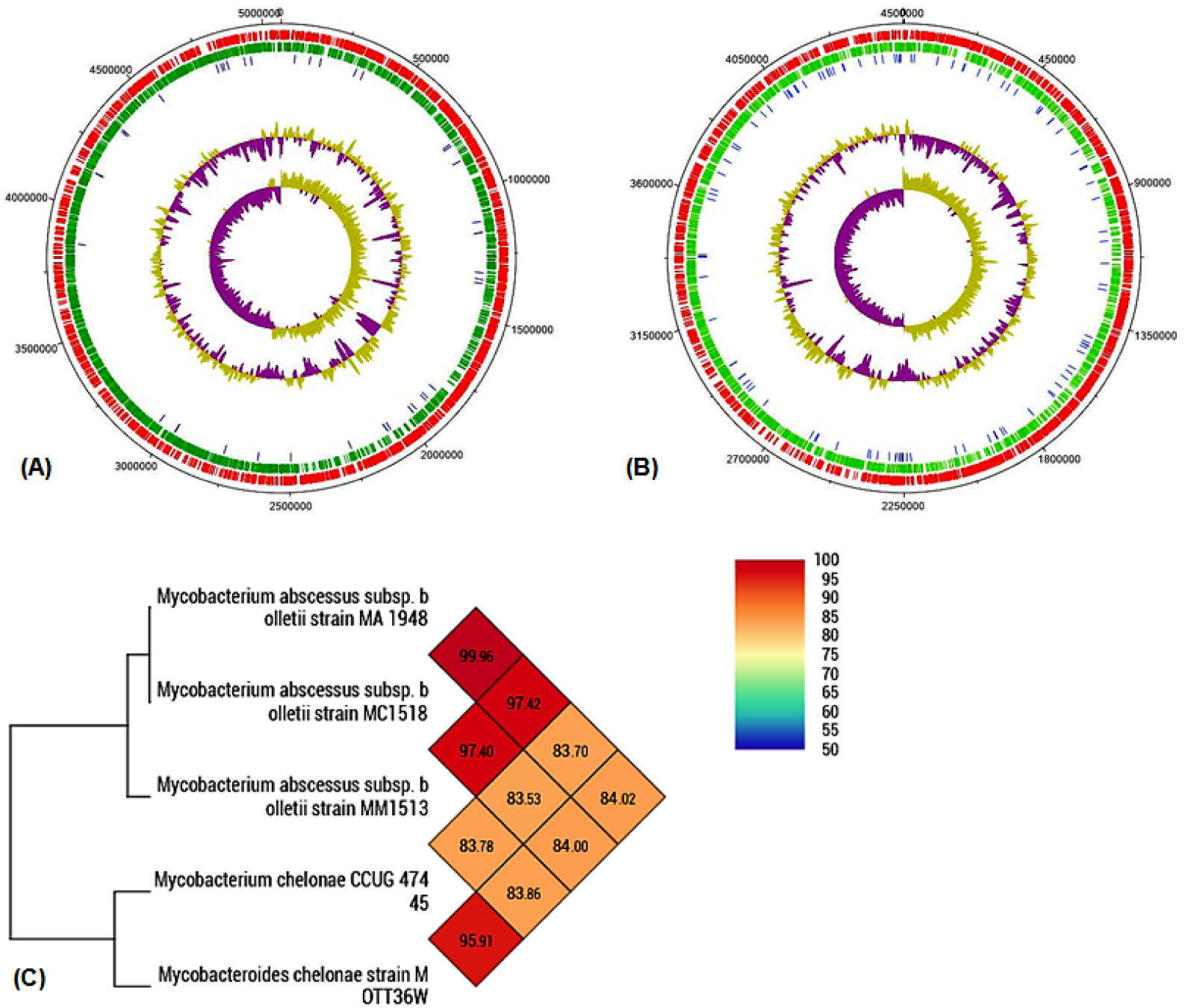
(A) Circular maps of the Strain MA 1948 and (B) MM1513 genome. Marked characteristics are shown from outside to the center: CDS on the forward strand, CDS on the reverse strand, Pseudogenes, GC content, and GC skew. (DNA Plotter was used) (C) OrthoANI value-based Phylogenetic Tree. Values were calculated using OAT software with available complete genomes all the strains. The results between any two strains are given by the junction point of the diagonals departing from each strain.

BUSCA is a new localization server that integrates several published Subcellular Prediction methods such as BaCelLo, MemLoci and SChloro and localization-related protein features such as DeepSig, ENSEMBLE3.0, BetAware, PredGPI, and TPpred3 **[57]**. (http://busca.biocomp.unibo.it/). The FASTA Sequences of proteins from previous steps were retrieved from BATCH Entrez (https://www.ncbi.nlm.nih.gov/sites/batchentrez) using .txt file of Accession Numbers. The fasta file, so generated, was used as input to BUSCA and 3 Subcellular compartment prediction was performed. CELLO v.2.5 uses a multiclass SVM classifier that uses consensual score generated from 4 different scoring criteria: the di-peptide composition, the amino acid composition, the partitioned amino acid composition and the protein sequence composition based on the amino acids’ physicochemical properties **[127]**. PSORTb v3.0.2 **[128]** interface integrated into BLAST2GO was exploited for SCL which employs SVM, S-TMHMM, and SCL-BLAST against 11692 proteins with known SCL from bacteria (Gram-positive and negative) and archaea **[58]**. LocateP v2 server predicts the SCL of a protein by mimicking protein targeting and secretion processes (http://bamics2.cmbi.ru.nl/websoftware/locatep2/locatep2_start.php) **[59]**. In the case of disputable localization, we considered either BUSCA or BLAST2GO Annotation prediction.

#### Virulence Prediction

Virulent Proteins assist bacterium in colonizing the host and pathogenesis and these proteins could be either Cytoplasmic, Membranous or Secretory. They help in adhesion, adaption to the changing environment, and protection against host immune response **[60]**. It is; therefore, prioritization of these proteins is necessary since they act as potential Drug Targets and Immunogenic Vaccine Candidates **[61]**. We used MVirDB and VirulentPred server (http://203.92.44.117/virulent/) to carry out Virulence prediction. In MVirDB, regular BLASTp was run and proteins with identity ≥ 30% and bit score of 100 were considered as virulent and taken for further analysis **[64]**. Proteins predicted as Pathogenic were further checked for virulence by VirulentPred with 0.6 as Threshold Value for the Cascade SVM-based Module **[65]**. The proteins so obtained were carried further for Vaccine development.

### Phase III: Reverse Vaccinology approach & Chimeric Vaccine Construction

#### Antigen Prediction

The virulent plasma membrane and extracellular proteins were sorted out from the Subtractive Proteomics Approach employed in Phase II and the Antigenic property of the proteins was assessed *in-silico* by Vaxijen Server v2.0 using a threshold of 0.50 **[66]**. Proteins with Antigenicity scores below 0.5 were considered as Non-Antigenic and were excluded from the study.

#### Physicochemical Characterization and Instability Index Prediction

The proteins screened from the above step were analyzed for stability in a test tube environment. The unstable proteins would otherwise degenerate in a test tube environment during isolation. Ii >40 was considered as benchmark above which the proteins were considered unstable and were excluded from the study. For Ii prediction and physicochemical characterization, XtalPred Server was used **[68]**.

#### Protein Clustering

Since proteins for epitope prediction were a subset of core-genome, there was a high probability for a homologous protein to occur multiple times as putative vaccine targets. To remove redundancy and use an only unique subset of proteins, we performed protein clustering using the h-CD-Hit Algorithm **[69]** (http://weizhong-lab.ucsd.edu/cdhit-web-server/cgi-bin/index.cgi?cmd=h-cd-hit) with 90% homology criteria.

#### Protein Structure Prediction

Due to a large subset of putative targets, it was not feasible to subject all predicted targets for epitope propensity analysis. Therefore, we considered the top 4 proteins having the highest VaxiJenScore for Antigenicity. Further, we modeled the protein structure using the I-TASSER of Zhang Lab **[70]**. The five models were produced for a single protein sequence and were then evaluated based on the Ramachandran Plot generated using SAVES v5.0 (Fig 4S). The model with a smaller number of residues in the disallowed region was selected for further model refinement**[71]**. We used a high-resolution structure refinement method i.e. ModRefine (https://zhanglab.ccmb.med.umich.edu/ModRefiner/) which improves poor rotamers by simulating both protein backbones and side-chain **[72]**. The Secondary structure predicted using PSIPRED 3 was used as a reference against which simulations were run. With DISOPRED 2, we tried to find our intrinsically disordered regions **[80]**. The model so generated were then subjected to other structure validation methods available on SAVES v5.0 again. We also assessed our model quality by using ProSA-Web and the Z-score indicated overall model quality **[73]**. The final model quality was assessed using Protein Quality Predictor-ProQ that uses two trained Neural Network: one that predicts LGscore and one that predicts the MaxSub score to assess the protein quality into three categories; Correct, Good and Very Good **[74]**. We finally used the GO-FEAT server for the functional annotation of the proteins using *e-value* 1e^-03^ **[75, 81]**.

**Figure 4:**
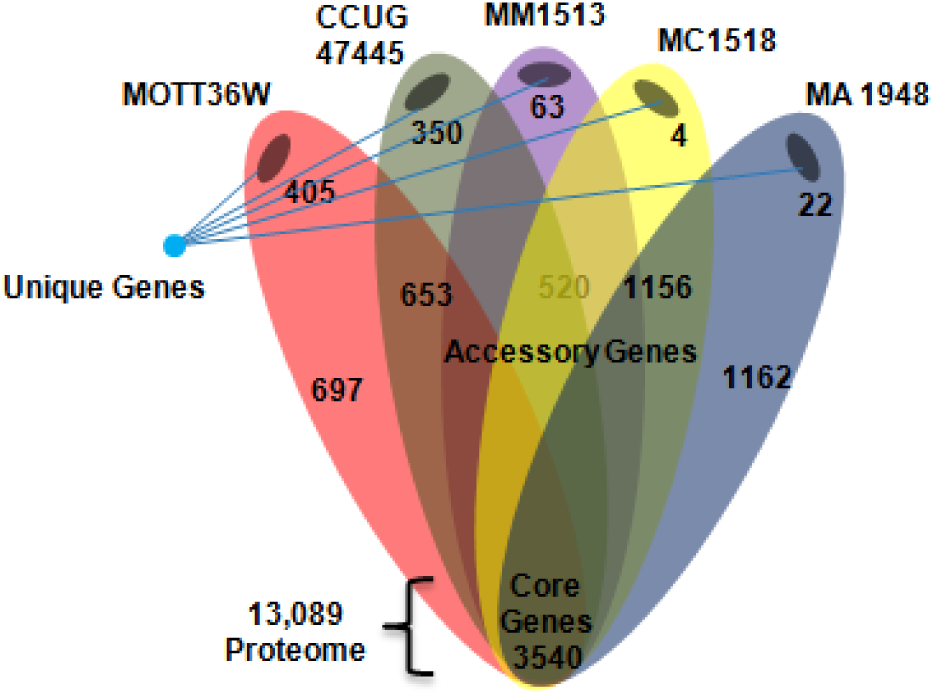
The Pan-core gene repertoire of 5 Mycobacteroides strains. The genes were classified into Core genes, Accessory gens and Unique Genes. 3540 genes comprise the core genome that eventually translates into 13,089 proteome. MOTT36W has the highest number of unique genes whereas MC1518 has the lowest.

#### Conserved Syntenic Block Prediction

The conserved syntenic regions shared among complete genomes of 5 Mycobacteroides strains were predicted using the Absynte server (http://archaea.u-psud.fr/absynte/) for four putative vaccine targets in the present study. Conserved Synteny /Gene order conservation of shared synteny is a feature of common descent and refers to the preservation of gene order across species **[76, 77, 78**].

#### B-Cell Antigenic Determinant Identification

The B-Cell antigenic determinants are segments of antigen that bind to immunoglobulin or antibody and generate humoral immunity **[87]**. Therefore, identification of B-Cell epitopes and their characterization is an important step for vaccine design. The linear B-Cell epitopes refer to continuous stretches of amino acid that forms an epitope-paratope complex with antibodies and were predicted using BCPred**[88]**, FBCPred server, ElliPro Suite of IEDB **[89]** and IEDB Antibody Epitope Prediction suite **[90]**. We used default parameters for prediction of non-overlapping epitopes using BCPred**[88]** and FBCPred with a specificity of 75% and epitope length of 20 and 14 respectively. For ElliPro Suite we used 0.5 as the minimum score and 6 as a minimum distance for the prediction of both continuous and discontinuous epitopes (antigenic patches) using the tertiary structure of modeled proteins. IEDB Antibody Epitope Prediction suite uses biochemical parameters such as flexibility, exposed surface, hydrophilicity, hydrophilicity, solvent accessibility, polarity, and amino acid antigenic propensity to predict continuous epitopes **[91]**. The server employs various prediction methods and expresses antigenic regions in a user-friendly and graphical method for a specified threshold. The methods include Chou and Fasmanbeta-turn prediction **[91]**, Emini surface accessibility scale **[92]**, Karplus and Schulz flexibility scale **[93]**, Kolaskar and Tongaonkar antigenicity scale **[94]**, Parker Hydrophilicity Prediction **[95]**, Bepipred-1.0 Linear Epitope Prediction **[96]**. We then built the epitopes by summing up peptides having overlapping regions. To check for overlapping regions amidst multiple epitopes, we used ClustalOmega (https://www.ebi.ac.uk/Tools/msa/clustalo/) to carry out MSA of peptides **[97]**. Only peptides which were predicted by at least two methods mentioned above were considered for the consensus B cell epitopes construction.

#### MHC Class I Binding Prediction

T-Cell MHC Class I epitope prediction was made by using the standalone tool of IEDB i.e. Tepitool for the screened proteins. We used a panel of 27 reference alleles (16 HLA A subtype and 11 HLA B subtype) for both A and B groups that cover >97% of the population and are enlisted in the List 1S, Supplementary section. The protein sequence was used as an input query and a window size of 9mers to predict short non-overlapping, non-redundant antigenic peptides **[98, 99, 100]**. We used the IEDB Recommended method for prediction and chose a threshold consensus percentile rank to be ≤ 1%. IEDB Recommended prediction method is a consensus ranking method that uses prior knowledge of predictions and the prediction performance and chooses the best available prediction method for any MHC Molecules. The decreasing order of prediction method is as follows: Consensus > ANN (Artificial neural network, also called as NetMHC, version 3.4) > SMM (Stabilized matrix method) >NetMHCpan (version 2.8) > Comblib_Sidney2008 (Scoring Matrices derived from Combinatorial Peptide Libraries).

#### MHC Class I Peptide Clustering

We clustered MHC Class I peptides to remove redundancy due to identical peptides. We used 100% identity as the threshold to cluster peptides based on their identity and then selected a single peptide from every cluster and the HLA type for which it has the highest binding score **[101]**. The clustering was attempted using the Epitope Cluster Analysis tool of IEDB (http://tools.iedb.org/cluster/). We then mapped these peptides to the data generated from the MHC Class I processing analysis.

#### MHC Class I processing analysis

We also performed MHC Class I processing analysis using protein amino acid sequence in FASTA format and IEDB Recommended method with an alpha factor of 0.2 **[100]**. Type Immunoproteasomes was used for analysis which is induced by IFN-γ secretion and is found to increase and enhance antigenic presentation on the surface of Antigen Presenting Cells, the detailed mechanism of which is discussed elsewhere **[102]**. Three scores; proteasome Cleavage score, TAP Transport score and MHC Binding scores are calculated, however, we used the consensus total score that combines all three above-mentioned scores for further analysis.

#### MHC Class I Immunogenicity Prediction

The resulting peptides were checked for their ability to elicit the immune response. Binding prediction generates a large set of peptides, however, not all peptides are able to evoke immunity to the same extent. We, therefore, used IEDB MHC Class I Immunogenicity prediction tool to identify peptides with high antigenicity scores in addition to the high binding score. The peptides sorted from the above step were used as input and we masked the 1^st^, 2^nd^ and C-terminus amino acid from the immunogenicity score. Peptides having positive Immunogenicity Scores were subjected to population coverage analysis **[103]**.

#### MHC Class II Antigenic Determinant Prediction

The MHC Class II prediction was also performed using IEDB standalone workbench i.e. Tepitool. We used the IEDB recommended method with a reference set of 26 alleles (15 DR, 5 DP, and 6 DQ alleles) with a default window of 15mers and not more than 10 overlapping regions were allowed. A threshold of ≤ 10 % consensus percentile rank was considered for the prediction. The resulting peptides were clustered. We then used only the top 10% of the total peptides for population coverage analysis.

#### MHC Cluster Analysis and Functional Relatedness

In addition, were used MHCcluster 2.0 Server to cluster the reference set of 27 alleles used for MHC Class I prediction and 26 alleles used in MHC Class II prediction and visualize their functional relatedness in the form of heatmaps and phylogenetic tree **[104]**. The server clusters the MHC molecules based on their binding specificity. The analysis would further strengthen our prediction and serves as an additional crosscheck for the predictions made through IEDB analysis **[105]**.

#### Population Coverage analysis of MHC Class I & II Epitopes

Given the polymorphic nature of MHC molecules, there are hundreds and thousands of HLA alleles and their subtypes. The epitopes will provoke a significant immune response in the candidates whose MHC molecules have a high affinity toward them **[106]**. Because different HLA types and subtypes are localized differently and occur dramatically at different frequencies and depend on the ethnicities and origin of the individual, we undertook the dataset of the “World Population” with a view to discriminate peptides covering major population portion **[106]** (Figure: 1). We prepared an input .txt file with peptides tab separated by their respective HLA subtypes. The population coverage analysis was carried out separately for Class I and Class II peptides keeping the other parameters constant. This would help surface peptides that would be used eventually for global vaccine production.

#### Antigenicity and Toxicity Analysis

We determined peptidic antigenicity and toxicity to evaluate if the peptides continue to carry immunogenic property and whether the peptides would have a toxic effect on host cells. The peptidic antigenicity was predicted using VaxiJen v2.0 with a threshold of 0.5 **[66]** (Figure: 1). The peptidic toxicity was calculated using the ToxinPred online server (http://crdd.osdd.net/raghava/toxinpred/multi_submit.php) that uses an amino acid sequence of peptides as input **[107]**. The toxicity analysis also deciphers other Physicochemical Parameters such as Hydrophobicity, hydropathicity, Amphipathicity, net charge, pI, Molecular weight, steric hindrance, etc. The prediction helps us predict whether the toxicity generated against bacterial cells inside the host cell would affect the host tissues in the vicinity or not **[107]**.

#### B cell Epitopes Mapping on T cell Epitopes

We used B cell predicted epitopes as templates and mapped them back individually to the data generated in MHC Class I and Class II epitope prediction. The underlying idea was to find out B cell epitopes overlapping with the regions of T cell epitopes so that the number of linkers would eventually get minimized in the vaccine engineering steps. (For results: This is a method we adopted to minimize the number of “Junk” epitopes that are usually formed at the junction of two peptides in a polyepitopevaccine)**[114]**. We then used the respective B cell immunogenic peptides and T cell peptides (both class I and class II) to build a consensus peptide (Figure 1). The final peptide construct, therefore, represents a continuous stretch of amino acid containing B cell epitopes, MHC Class I and Class II epitopes.

#### Multi-epitope vaccine Model Construction

For constructing a multi-epitope novel vaccine model with high immunogenicity and low allergenicity and toxicity, we analyzed multiple combinations of sequence constructs. While constructing vaccine sequence we kept the number of sequences in all constructs equal, however, their length varied. We used 6 consensus peptides that were joined together with the help of linker peptides. To enhance the immunogenicity of the vaccine constructs we added the following four different adjuvants: HBHA protein, HBHA conserved sequence, β-defensin, and L7/L12 ribosomal protein. A universal immunogenic PADRE peptide sequence was also conjugated using the EAAK linker with the adjuvant in each construct. The sequence is reported to induce helper T Cells (CD4+ T-cells) that enhance the potency and efficacy of the vaccine. Immuno-adjuvant HBHA (heparin-binding hemagglutinin) and L7/L12 ribosomal protein binds to TLR4/MD2 complex but not TLR2 and show agonistic effect activating Dendritic Cells (DC’s) **[115]**. HBHA causes polarization of T effector cells via DCs maturation and is studied to have an immune-stimulatory effect **[116]**. β-defensin adjuvant, however, exhibits an agonistic effect to TLR4 as well as TLR2, and TLR1 and leads to co-stimulatory expression of CD40. CD80 and CD86 molecules on myeloid DCs and monocytes **[117]**.

We also used target signal peptides at both N- and C-terminus that is reported to enhance Helper-T cell response when present in conjugation with polypeptide **[118]**. We used both original and modified version of HER2 signal peptide that would ensure vaccine targeting to Endoplasmic Reticulum (ER) and Secretory pathway and an 11 amino acid long C-terminus LAMP-1 peptide which ensures immunogen targeting to the lysosome for disintegration and subsequent presentation of MHC Class II-epitope complex on the cell surface[patent]. We used GGGS and HEYGAEALERAG peptide linkers to join epitopes whereas adjuvants sequences and signal peptides were linked with the help of EAAAK linkers at N-terminus and GGGS liker on C-terminus **[98, 99, 100]** (Fig 5S).

**Figure 5:**
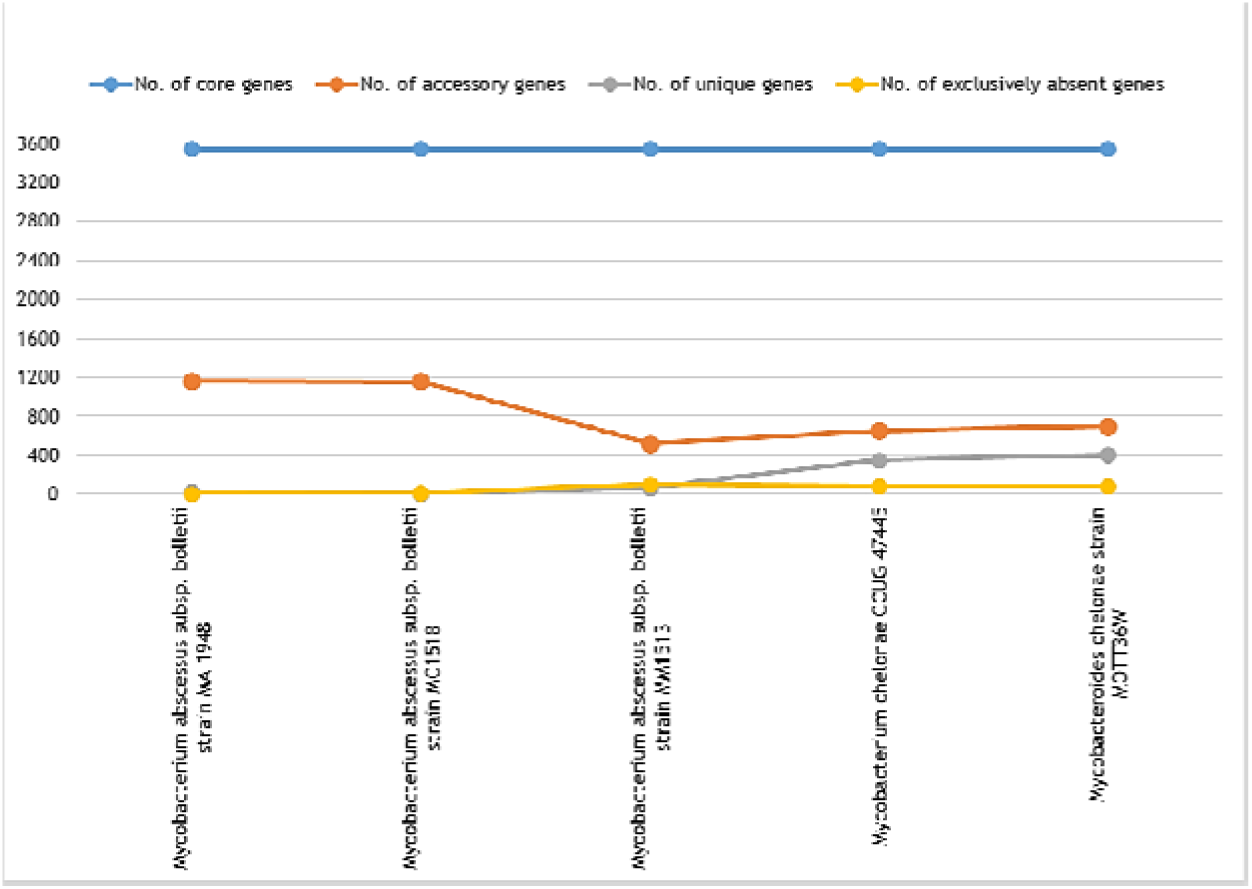
The comparative graphical account of 4 categories under which BPGA cluster genes.

#### Determination of Allergenicity, Antigenicity, Solubility and Solvent Accessibility of Vaccine models constructed

To ensure our vaccine construct were non-allergic, we subjected our 8 constructs for allergenicity analysis. We used Hybrid approach of AlgPred server that employs integrated methods such as SVM amino acid composition or dipeptide-based method, IgE epitope mapping, BLAST against ARPs (allergen representative peptides) or MAST (Motif Alignment & Search Tool)-MEME (Multiple Em for Motif Elicitation) suites to measure putative allergenicity if any **[119]**. The Allergenicity threshold for SVM (amino acid composition) by default was taken to be −0.4 and for SVM (dipeptide) to be −0.2. We did this analysis for vaccine constructs with signal peptides and for vaccine construct without signal peptides. The non-allergic constructs were taken further for antigenicity assessment using ANTIGENPro of Scratch Protein Predictor (http://scratch.proteomics.ics.uci.edu/) **[100, 131]** followed by Vaxijen v2.0. SOLpro of Scratch Protein predictor was used for vaccine Solubility estimation and Solvent accessibility. SOLpropredicts protein’s propensity of being soluble when overexpressed in *E. coli* using SVM classifier **[100, 131, 132]**.

#### Physicochemical Characterization of Vaccine Constructs and Structure Prediction

To predict the secondary structure of the vaccine construct we used PSIPRED v3.3 workbench (http://bioinf.cs.ucl.ac.uk/psipred/) that performs a PSI-BLAST and then employ the output to feed-forward trained Neural Network to predict secondary structure. For protein disorder prediction we used DISOPRED 2 (https://iupred2a.elte.hu/) to identify intrinsically disordered regions **[80]**. The vaccines were modeled using I-TASSER **[70]** and the structure was validated using the strategy mentioned in Figur 4S.

#### Estimation of Local energy frustration of vaccine tertiary structure

To discover local regions of energy frustration in the vaccine model, we used the ProteinFrustratometer 2 server (http://frustratometer.qb.fcen.uba.ar). This helps us localize regions of high energy that arise due to the electrostatic interaction between amino acids, correlates with regions of potential biological importance and might be involved in functions such as binding or allosteric transitions **[120]**. The low energy region, on the other hand, represents stable backbones and tend to remain conserved both in structure and conformations **[121]**. We used refined models of Vaccine constructs and included Long-range electrostatic interactions with a default K value of 4.15 (for the aqueous system) that is a measure of electrostatic strength of the system. We used new sequence separation value = 3 for the calculation local amino acid densities and used all three modes viz the native conformation (Conformational frustration), mutational (mutational frustration) and single residue mode (Single-residue level frustration) to calculate frustration index and the variance of the energy values **[121]**.

#### Disulfide Bond Engineering via mutation and structure stabilization

We cautiously tried to mutate some amino acid residues and create Disulfide bonds for protein structure stabilization. However, we did not create a mutation in the regions that were predicted to be potentially binding regions during disorder and binding region analysis in the above steps since such regions are often flexible and aid in ligand-receptor interaction at the interface. For engineering Disulfide bond, we used refined model and mutated it using Disulfide by Design, version 2.12 with X_3_ (Chi_3_) angle to be +97°/-87° with the tolerance of ± 10° and C_α_ -C_ß_-S_□_ angle to be 114.60° with the tolerance of ±10° **[122]**. We used the bond energy ≤ 2.2 kcal/mol as the primary criteria to sort out favorable and energetically stable S_2_ bonds followed by a torsional angle as secondary criteria.

#### Receptor Preprocessing and Protein-Protein Docking

Molecular docking of all 6 vaccine constructs was carried out with the following 8 receptor protein molecules for Binding affinity was estimated. The MHC Class II receptors that were used includes HLA-DR B1*03:01 (PDB ID: 1A6A), HLA-DR B3*02:02 (PDB ID: 3C5J), HLA-DR B5*01:01 (PDB ID: 1H15) whereas MHC Class I receptors proteins includes HLA-B*35-01 (PDB ID: 2H6P), HLA-A*11:01 (PDB ID: 2HN7), HLA-B*51:01 (PDB ID: 1E28), HLA-A*24:02 (PDB ID: 2BCK). The preprocessing of these peptides was carried out manually using UCSF Chimera. The homodimers were reduced to a single chain to reduce docking time and non-amino acid molecules (ligands, ions, and solvent water) were removed to prevent hindrances during docking. The PatchDock server was used to perform rigid-body molecular docking of the engineered vaccine with the processed receptors based on shape complementarity principles **[123]**. The patch dock algorithm divides the protein’s surface into small patches (Convex, concave and flat) using a segmentation algorithm that is the superposed using shape matching algorithm (Sorted according to binding energy function) **[123]**. The top 10 docking solutions obtained from PatchDock were subjected to docking score refinement using FireDock. FireDockrefines the docked poses by optimizing side-chain conformations and rigid body orientation by performing Monte Carlo simulation **[124]**.

Similarly, the best vaccine candidate obtained from the above docking results was docked further with TLR 4/MD2 complex (PDB ID 4G8A) and results were refined using Firedock. The complex was simulated using Groningen Machine for Chemical Simulations GROMACS v5.1.2 **[136]**described elsewhere **[125]**. The binding affinity of best vaccine candidate with its all 8 receptor molecules was quantified using PRODIGY along with dissociation constant, the number of Interfacial contacts and Interacting residues are tabulated under **[137]**.

#### MD-Simulation of TLR/MD2-Vaccine Complex and Energy Frustration Estimation

Molecular dynamics and simulation of the docked complex of TLR/MD2 and Vaccine was carried out using GROMACS v5.1.5 using OPLS-AA/L all-atom force field (2001 amino acid dihedrals) to study the physical movement of the interfacial atoms and the stability of the complex in explicit water box (dodecahedron) at 309K for 150 ns **[136]**. We used both NVT and NPT ensembles to mimic the real experimental conditions.

#### Codon Optimization of for Vaccine Cloning

OPTIMIZER was used for codon optimization of the Vaccine Construct sequence **[134]**. The protein sequence was back-translated using guided Random method that is based on the Monte Carlo algorithm and used mean Codon usage Table of *Escherichia coli* str. K12 substr. MG1655 for optimization. The sequence so obtained contained various Type II Restriction sites. To further optimize the sequence and make it Industry ready, we used IDT Codon Optimization Tool. In addition to codon optimization, IDT also predicts other complexities hidden in the sequence such as extreme GC, hairpin sequence and a gene burdened with extensive repeats. A complexity score is calculated that establishes how easily a gene can be produced synthetically and a score below 7 demarcates Low Complexity. The final sequence that was adapted for overexpression in E Coli was incorporated with sticky ends AGCT. We then inserted this sequence in *E coli pET28a* vector DNA obtained from http://dnasu.org/DNASU/GetVectorDetail.do?vectorid=319 using SnapGene software for its successful expression **[138]**.

## Results and Discussion

### Phase I: Deciphering Genomic Landscape

#### Genomic Features of Mycobacteroides and Evolutionary Disparity

To study the gene repertoire of 5 unannotated genomes of Mycobacteroides we carried out multiple sequence alignment using MAUVE with default parameters **[36]**. The progressiveMauve constructs “positional homology multiple genome alignments” by employing 3 algorithms: The sum-of-pairs breakpoint score for scoring alignment-anchor across the genomes, greedy heuristic for the score optimization and homology-driven Hidden Markov model (HMM) to eliminate spurious alignment **[36]**. The whole-genome alignment shown in Figure 5 represents the genomic relatedness among 5 strains of Mycobacteroides. The genome of Mycobacterium abscessus subsp. bolletii strain MM1513 is below the bold Blackline and indicates that the whole genome of this strain aligns in the reverse complement (inverse) orientation. Also, it is evident from the alignments that events of deletions have taken place in Strain MM1513, CCUG 47445, and MOTT36W. The gene order can be inferred to be preserved among all 5 strains (Figure 2). We used OAT (Orthologous Average Nucleotide Identity Tool) to measure overall relatedness amidst all 5 genomes. We usedOrthoANI algorithm to decipher the genetic distance and used the OrthoANI values **[37]** to construct a phylogenetic tree as shown in Figure 3. Strain MA 1948 is closely related to strain MC1518 (99.96% homology) followed by MM1513 (97.42%). Strains CCUG4745 and MOTT36W are more related to each other (95.91% homology) than with other strains. However, the overall shared homology was no less than 83%. (For data, refer to the List 6S Supplementary section)

To have a closer look at the genomes of these strains, we used JAVA based Artemis Visualization interface to find out the cause of deletions that have occurred in the genomes **[44]**. We tried to tabulate our findings of the number of coding sequences (CDS), and pseudogenes present across the genomes while keeping MA 1948 as a reference genome. It was found that the number of pseudogenes were least in strain MA 1948 followed by MC1518 and were maximum in MM 1513 strain [Table 1]. Also, as we move down the phylogenetic tree (Figure 3), we observe a continuous increase in pseudogenes and multiple stop codons with maximum concentration in MM1513 strains. Similarly, we observed that the number of coding sequences reduces as we go down the tree from the strain Ma 1948 with the least no of CDS in MM 1518. The GC content plot was viewed @ window size of 262 in Artemis and we found that MM1513 strain comparatively has more regions of atypical GC content (for instance at position 161291-181718, 421297-438521, etc.) than other strains. This is indicative that MM1513 strain has undergone reductive evolution since the presence of a large number of pseudogenes, lower GC content is hallmarks of evolutionary bottleneck that the species must have faced which eventually leads to downsizing of the gene repertoire thereby confirming Reductive evolution **[46]**. During this course, the reductive evolution, the genes of metabolic importance or virulence and their function could be lost due to genome shrinkage and genome decay **[47]**. Since the genomes are newly sequenced and unannotated, it’s difficult to identify which genes lost their functionality.

**Table 1:** Number of Coding Sequences (CDS), Pseudogenes and Tandem Repeats (TRs) present in 5 strains of Mycobacteroides.

| Organism name | CDS | Pseudogenes | Tandem Repeats |
| --- | --- | --- | --- |
| *Mycobacterium abscessus subsp. bolletii* strain MM1513 | 4427 | 111 | 145 |
| *Mycobacterium chelonae* CCUG 47445 | 4867 | 63 | 145 |
| *Mycobacterium abscessus subsp. bolletii* strain MC1518 | 5003 | 55 | 193 |
| *Mycobacterium abscessus subsp. bolletii* strain MA 1948 | 5018 | 41 | 197 |
| *Mycobacteroides chelonae* strain MOTT36W | 4984 | 91 | 165 |

Tandem Repeats Finder was used to identify tandem repeats in each genome **[45]**. The largest number of Tandem Repeats (TRs) were found to be populated in MA 1948 and least in MM1513 and MC1518 (Table 1). Genetic Loci containing TRs are hypermutable and prone to events such as recombination or strand slippage. The abundance of TRs in a bacterial genome makes it unstable enabling bacteria to modulate (Switch on/off) genes/ functions to withstand changing the environment and tolerate stress for short evolutionary time-frame without increasing overall mutation rate **[48]**.

#### Antimicrobial Resistance (AMR)

To find out if the genome of the strains were populated with any Antimicrobial-resistant gene, we used the RESFinder server **[38]**. Only two out of 5 strains; *Mycobacterium abscessus subsp. bolletii* strain MA 1948: (2345402-2345818) with the identity of 99% (413/417 nucleotides) and *Mycobacterium abscessus subsp. bolletii* strain MC1518: (2328523-2328939) with an identity of 99% (413/417 nucleotides), were found to possess Erm (41) methylase gene as shown in Fig 6S, Supplementary section. Erm (41) confers to Macrolide, Lincosamide and Streptogramin B (MLS) resistance (Accession No: EU177504) and is reported to be present in *abscessus* strain but absent in chelone strain **[24]**. However, the absence of Erm (41) from MM1513 indicates that the functional Erm gene is lost during gene decay. (https://www.ncbi.nlm.nih.gov/nuccore/EU177504).

**Figure 6:**
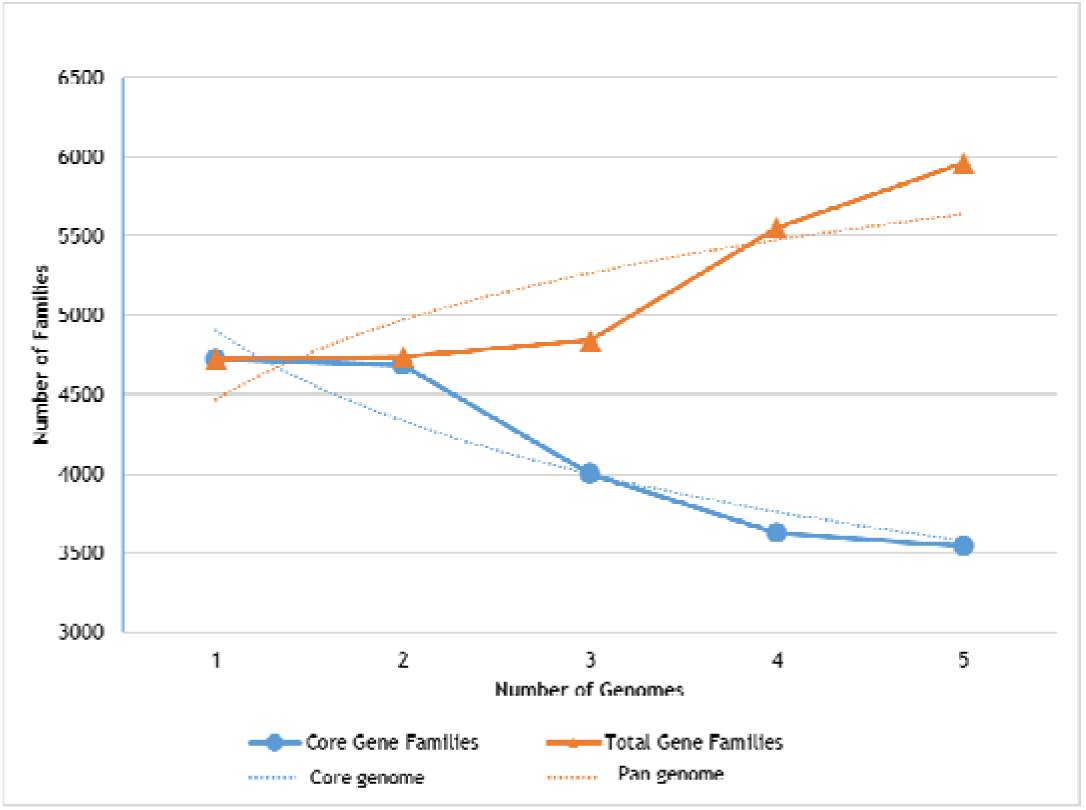
Pan-Core Repertory according to the number of genomes added sequentially. The power-fit and exponential model of pan and core genes are demonstrated by orange and blue lines, respectively, while the corresponded equations are as follows: Power-fit Law equation: f(x)=a.x^b and exponential curve equation: f1(x)=c.e^(d.x) and where a= 4451.45, b= 0.151203, c= 5251.34, d= −0.0834852

#### Pan-Core Analysis and Comparative Genomics

The genomes were subjected to Pan-Core Analysis using BPGA Pipeline **[39]**. Examination of genomes revealed that the gene repertoire of all five Mycobacteroides contained 3540 core genes with a variable number of accessory, unique and exclusively absent genes (Figure 4). As we move from left to right in Figure 5 we observe that the number of accessory genes drops abruptly from MM1513 strain onwards and the number of unique genes increases.

A Clusters of Orthologous Group (COG) classification of Mycobacteroide’s gene repertoire indicates that unique genes were majorly assigned to COG categories R (General function prediction only), followed by K (Transcription) and Q (Category Secondary metabolites biosynthesis, transport, and catabolism), whereas core genes tended to be in the categories R (General function prediction only), K (Transcription), E (Amino acid metabolism and transport and I (lipid transport an metabolism). Distribution of COG categories within Mycobacteroides strain’s gene repertoire is demonstrated graphically in Fig 1S, Supplementary section.

The power-fit and exponential decay model was used to extrapolate pan-core gene changes after the sequential addition of each new Mycobacteroide genome (Figure 6). For pan-genome (Total gene repertoire), the expected size of gene repertoire was 5960 genes and estimated to decrease progressively with addition of each new genome in accordance to the power-law equation “f(x)=a.x^b” where expansion rate b = 0.151203. In contrast, for core genome (common/ shared genes), the expected size was 3540 genes in accordance to exponential equation f1(x)=c.e^(d.x) and is estimated to decrease with d= −0.0834852. The pan-genome was still open but it was predicted that it may be closed soon **[39]** (Figure 6).

The number of functional proteins, GC content and no. of gene families were plotted graphically(data not shown). We found a low number of Functional proteins in the MM1513 strain. The genes were clustered in 4 different functional categories: Information storage and processing, cellular processes and signaling, Metabolism, poorly characterized genes (data not shown). KEGG classification also revealed that most of the Accessory and Core genes belonged to the Amino acid metabolism functional group (see Fig 2S, Supplementary section).

#### Prophages and CRISPR/Cas System

The PHASTER tool was deployed to find the phage genes, if any, are present in Mycobacteroides **[40]**. Two Intact Phage (score > 90) were found to be present in each; *Mycobacterium abscessus subsp. bolletii* strain MC1518 and *Mycobacterium chelonae CCUG 47445* the details of the regions of which are given in Table 2. On the other hand, incomplete phages with the length between 7.5kb to13.8kb were found in multiple strains and came from different species of *Mycobacterium phage: Gaia* NC 026590(10), *Severus NC 021307(1), and Charlie NC 023729(2)*. The ORFs of the Intact Phages were visualized (data not shown).

**Table 2:** Intact Prophage Regions and their Bacteriophage Source.

| Protein Accession Number | Region | Region Length | Score | Total Proteins | Region Position | Most Common Phage | GC % |
| --- | --- | --- | --- | --- | --- | --- | --- |
| NZ_CP009613.1 | 1 | 50.2Kb | 150 | 78 | 1753180 to1803396 | PHAGE Entero Arya NC_031048(12) | 59.39% |
| NZ_CP007220.1 | 1 | 72.7Kb | 150 | 91 | 1521425 to1594189 | PHAGE ErwinivBEamP Frozen NC_31062(18) | 60.37% |

#### Genome Expansion and analysis for CRISPR-Cas gene presence

CRISPRFinder and CRISPRDetect were deployed for identification of CRISPR/Cas system, if any, present in the Mycobacteroides strains. The level of significance of predicted systems was evaluated based on Evidence Level. However, not more than 1 spacer sequence was identified in each genome with Evidence Level 1 (Low) **[40]**. Thus, the CRISPR/ Cas gene predicted were spurious and demarcated the absence of any such system in Mycobacteroides.

### Phase II: Subtractive Proteomics Approach

#### Non-homology analysis

A shortlisted input dataset of 13,089 core Proteins from Pangenome analysis was subjected to Non-Homology Analysis using BLASTp with an E-value threshold of 0.005 **[50]**. Approximately, half proteome of 5 Mycobacteroidal strains i.e. 6,047 Proteins were found to be non-homologous to Human Proteome. Proteins with zero hits for the E-value threshold 0.005 were regarded as non-homologs, whereas those exhibiting hits were considered as close homologs. The analysis was carried out thrice to remove false positives. The non-homology analysis is often the primary step in several *in silico* drug target identification studies to prevent any off-target side-effects **[67]**.

#### Essentiality analysis

The essential genes codes for proteins which are indispensable to cellular life and constitutes a minimal set of genes required by the cell to sustain a functioning cellular life **[62]**. It is, therefore, the second most critical step to identify essential proteins that are part and parcel for cellular life sustenance. The 6,047 Proteins were subjected to essentiality analysis using the DEG database with an expected value of 0.0001 **[50]**. Of 6,047 proteins, only 1,819 proteins were found to be essential to Mycobacteroides. Proteins with No-hits against the DEG database were considered non-essential. A confirmatory analysis was performed for all 6,047 proteins using PBIT Pipeline that queries the DEG database along with VFDB to trace essential proteins **[52]**. PBIT results were found to be in complete alignment with the DEG results.

#### Anti-Target Non-homology analysis

The 1,819 proteins were carried further to check for the presence of anti-targets if any. Anti-targets are an important consideration during the drug development process since they might cause long-term health issues or could be fatal **[53]**. The analysis against 306 anti-targets at the expected value of 0.0001 and < 30% similarity criteria revealed that only 2 out of 1,819 proteins were Anti-Targets and were therefore excluded from the dataset.

#### Gut-Flora Non-Homology

The complex ecosystem of our GI Tract helps in the assimilation and absorption of the poorly digestible dietary component, degradation of xenobiotics, and vitamin synthesis. To prevent the proteome of Gut microbiota from off-target effects we performed BLAST run against 95 gut microbes **[63]**. Out of the dataset of 1,817 proteins obtained from the previous step, we discovered that only 823 proteins were non-homologous to Gut Microbiota proteome at e-value of 0.0001 and < 30% rule and with not more than 10 hits rule got further reduced to 391. Proteins not following the above criteria were excluded from the study.

#### Protein Subcellular Localization

To characterize shortlisted targets suitability as candidates for Chimeric Vaccine construction or Drug Targets their distribution within the call was required. It has been reported in the literature that the proteins localized in the cytoplasmic region can be used as drug targets while the extracellular and membrane-bound proteins can act as putative vaccine candidates **[50]**. Our consensual SCL method as described in the methodology, conclusively, found that of 391 proteins, 215 proteins were found to be localized in Plasma Membrane, 156 Cytoplasmic Proteins, and 23 Extracellular Proteins Fig 7S. In the case of unknown/ disputable location, BLAST2GO **[58]** or results from BUSCA prediction were considered final.

**Figure 7:**
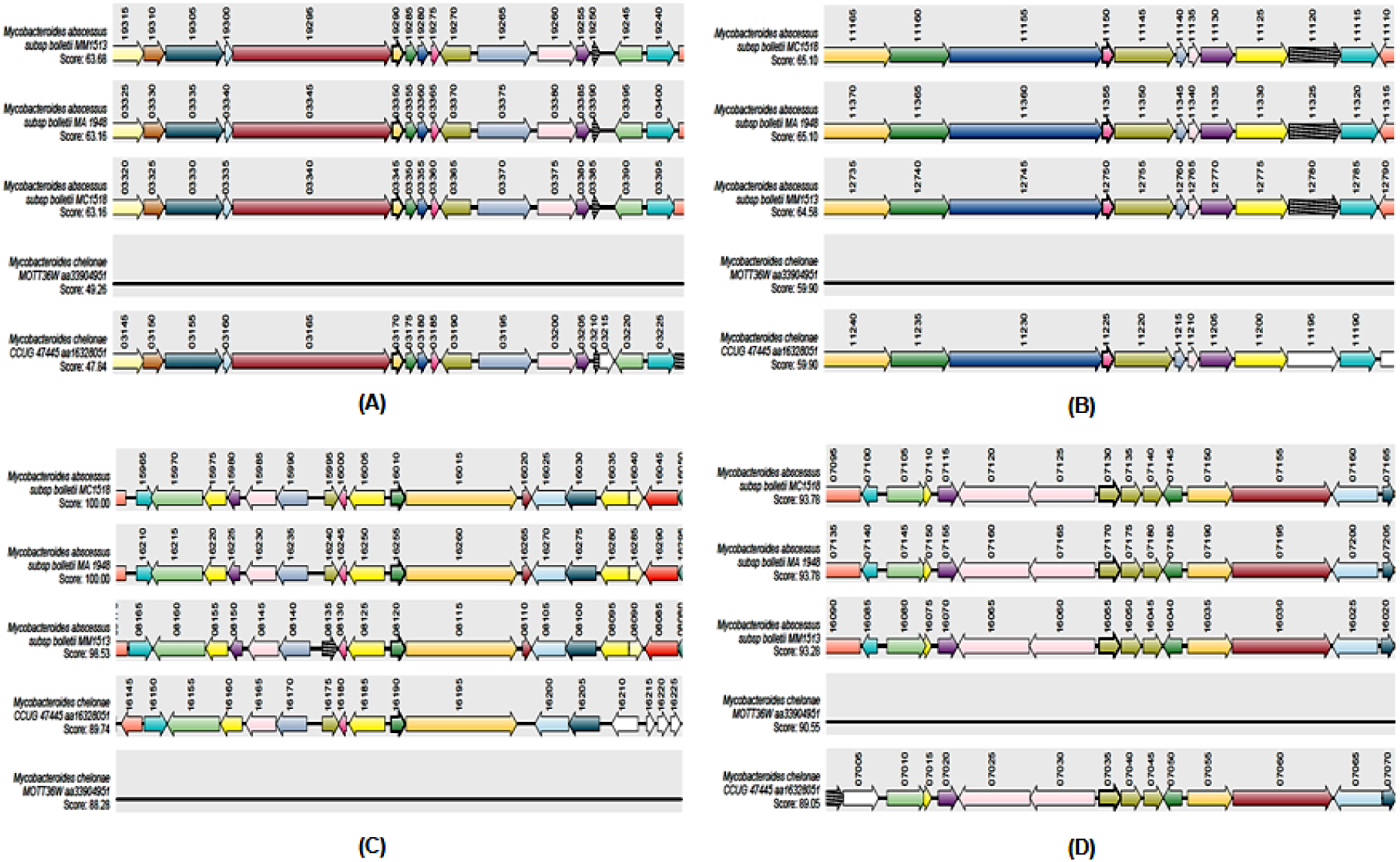
Local synteny prediction and representation in completely sequenced 5 Mycobacteroidal genomes along with synteny score. Synteny score is maximal bit score generated by performing blast against all sequences/reference blast score. Genomes are arranged in order of decreasing Synteny score. Colour codes are automatically generated representing a coding sequence. (For details on the interpretation of color code in this figure legend, please refer to the web version of this article. The proteins (A) WP_005064885.1 (B) WP_005091736.1, (C) WP_005111658.1, (D) WP_012296428.1 are highlighted by an arrow with bold Black Boundary.

#### Virulence Prediction

The final step of subtractive proteomics was to underline those proteins that could cause virulence in the host and provoke the immune response. Such proteins often serve as best Drug targets since such virulence factors are vitally responsible for the establishment and severity of the infection and their inhibition would render the pathogen avirulent **[50]**. Out of 391 proteins, 220 proteins were predicted to be pathogenic by MVirDB using the Hybrid Prediction scoring method (HMM and SVM) **[64]**. These 220 proteins were input into VirulentPred with a threshold of 0.6 to identify the Virulent Proteins. Of 220, 64 proteins were found Virulent. The Plasma Membrane and Excretory proteins from the Virulent protein’s dataset were used for Vaccine Construction and the cytoplasmic for Drug Target Identification.

### Phase III: Reverse Vaccinology approach & Chimeric Vaccine Construction

Of 64 Virulent Proteins obtained from the Subtractive proteomics approach, we chose 52 proteins that span Plasma Membrane or were predicted to be secretory in SCL analysis. Of these 52 candidates, 25 proteins were finally shortlisted using stringent criteria of >1 Threshold score of VirulentPred. The Allergenicity prediction was carried out on these 25 proteins to check if the Pathogenic proteins had the potential to trigger a significant humoral immune response by using Vaxijen Server v2.0 at a threshold of 0.5. VaxiJen classifies proteins to be antigenic/ Non-antigenic based on the physicochemical properties of the proteins **[66]**. All 25 proteins were found to be antigenic.

The 25 antigenic proteins were then checked for the protein stability in the test-tube environment. For this, we used the XtalPred server to estimate the Instability Index using the approach of Guruprasad et al., 1990 **[68]**. The Ii is a calculation of regional instability in protein by estimation of the weighted sum of dipeptides occurring more frequently in unstable proteins when compared to stable proteins. Proteins with instability index >40 were considered unstable and were deducted from the protein dataset. We found that 35 only 14 proteins were predicted to be stable in the Test-tube environment by XtalPred. All 4 proteins were predicted to be “very difficult to crystallize” by Expert pool method that delineates the crystallization probability using consensus score generated from 8 protein features: GI, pI, Ii, predicted secondary structure, length, predicted structural disorder, insertion score, predicted coiled-coiled region, predicted coiled secondary region **[68]**. However, the Random Forest method (RF) predicted high probability of WP_005091736.1 (pI: 4.67) and WP_005064885.1 (pI: 4.75) being easily crystallized, WP_005111658.1 (pI: 7.74) to be most difficulty crystallized while WP_012296428.1 (pI: 6.58) was predicted to exhibit moderate difficulty. Only WP_005111658.1 possessed Transmembrane helices from 5th to 24^th^ amino acids spanning the Plasma Membrane. XtalPred-RF uses TMHMM Server v. 2.0 to predict region with a high propensity of being in the transmembrane region **[68]**. SignalP-5.0 Server was used by XtalPred to check the presence of signal peptides. Proteins, except WP_005064885.1, were found to possess signal peptide; WP_005091736.1 with Sec signal peptide (Cleavage site between 31 and 33 AA), WP_005111658.1 with Sec/SPI (Cleavage site between 27 and 28), WP_012296428.1 with Lipoprotein signal peptide (Sec/SPII) (Cleavage site between 27 and 28) **[79]**.

The 14 stable antigenic proteins were then assessed for Homology to remove redundancy since the constituent proteins were a subset of the core genome. We used the h-CD-Hit algorithm to cluster proteins with similarity greater than or equal to 90% and they were eventually eliminated **[69]**. We found that the homologous proteins were clustered in 9 clusters with homology greater than 93.14%. There were proteins that were 97-98% similar. Thus 9 proteins (one from each cluster) were shortlisted. However, we carried further only 4 proteins with highest VaiJen Score (WP_005064885.1, WP_005091736.1, WP_005111658.1, WP_012296428.1).

We then went on to predict the secondary structure of these 4 proteins using PSIPRED v3.3 workbench. This prediction shall be used as an input to the I-TASSER for the Homology Modelling of these proteins (Fig 8S, Supplementary section). We also predicted natively disordered regions in the protein since these regions are highly flexible with hydrophilic and aromatic residues and are usually involved in molecular recognition/interaction with the receptors and orchestrate its binding **[80]**. Finally, the tertiary structures of the proteins were synthesized using I-TASSER and the secondary structure was given as a template. The structures so obtained were assessed for the quality parameters using SAVESv5.0. Ramachandran plot was compared for the 5 structures so produced by I-TASSER and the model with the least number of residues in the disallowed region was selected for refinement. Post-refinement of the model using ModRefine, we again compared the Ramachandran plot with the old plot as reference. We observed improvement in the spatial arrangement of residues and poor rotamers (data not shown). Further, the structure was also validated using ProSA **[73]** and ProQ **[74]** and the quality score was found to be “Very Good”. We also tried to annotate the proteins using Gene Ontology Functional Enrichment Annotation Tool (GO-FEAT) server that is freely accessible and is used for genomic and transcriptomic data annotation based on sequence homology search **[75]** (http://computationalbiology.ufpa.br/gofeat/). The FASTA sequences of 4 proteins were used for the homology search at *e-value* 1e^-03^**[81]**. It was found that both Protein WP_005064885.1 and WP_005091736.1 protein belongs to proline-glutamic acid (PE) family protein that has conserved PE motif at the N-terminal of aa sequence with variations in C-terminal region and is found to have a role in pathophysiology and virulence of *Mycobacterium*. It has been known to play an important role in immune evasion and generate antigenic variation **[82]**. WP_005111658.1 was predicted to be Putative membrane protein belonging to matrix metalloproteinases (MMPs). MMPs constitute a large family of Ca^2+^ and Zn^2+^ dependent endopeptidase and are capacitated to break the extracellular matrix (ECM) **[83]**. A study carried out elsewhere **[84]** highlights that MMPs production may play a pivotal role in the development of a protective immune response to mycobacteria and could be responsible for the impairment of tissue integrity.

**Figure 8:**
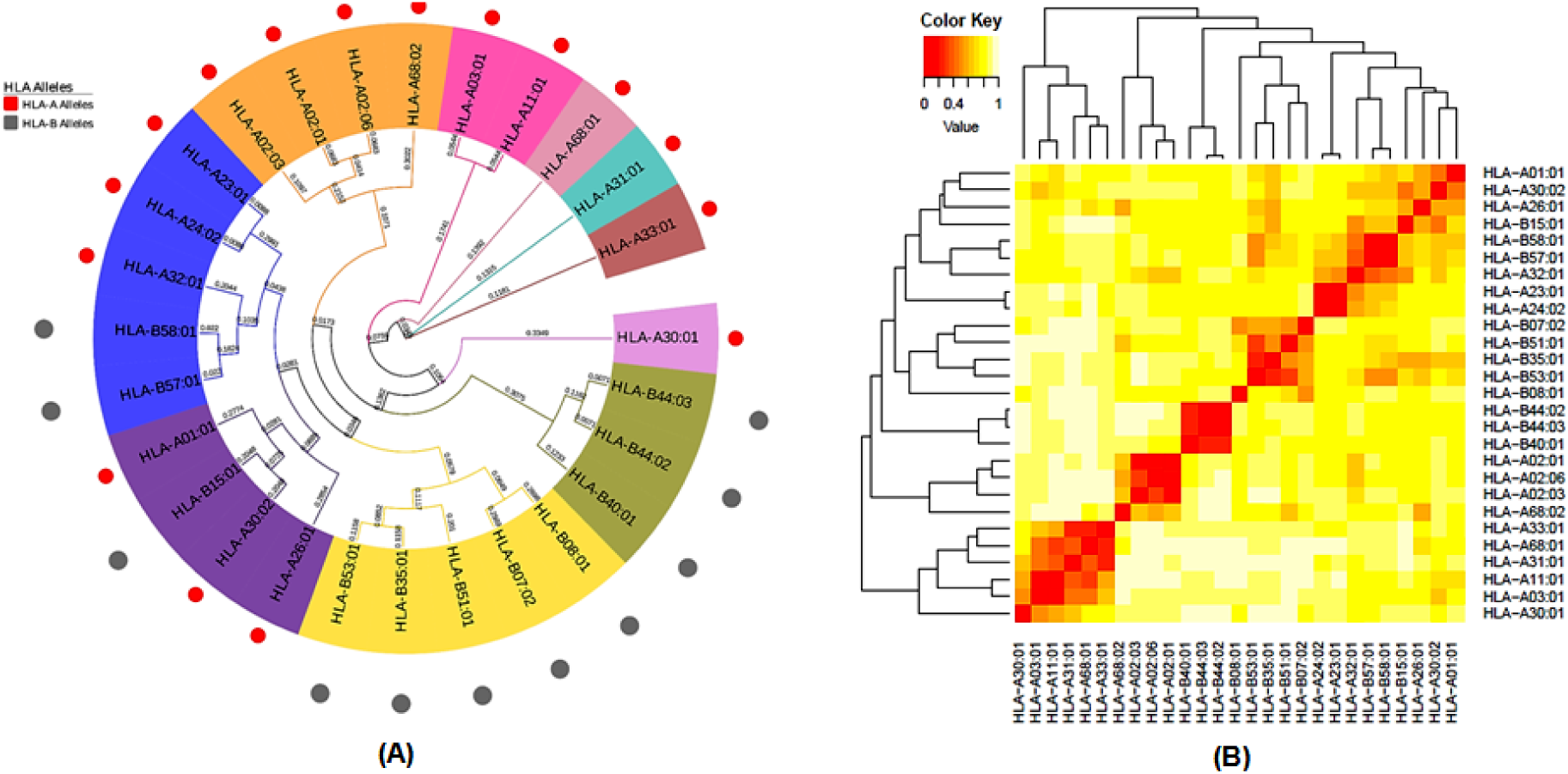
Functional relatedness of the HLA alleles for a reference set of 27 MHC Class I molecules on the basis of binding specificity (A) Represented the cluster analysis using Circular Phylogenetic tree for both HLAA- alleles and HLA-B alleles at a functional level (B) Represented using Heat Map. The Epitopes were clustered based on interaction with HLA molecules. In the heat map, the red color demarcates strong interaction and the Yellow region indicates the weaker interaction.

The amino acid sequences of the 4 proteins were used as input in the Absynte browser **[85]**to search local syntenies in completely sequenced 5 Mycobacterial chromosomes (Figure 7). The Synteny analysis was performed to trace the orthology among the N number of genomes (here 5) and identify the functional relationship among genes. The synteny analysis comprises 4 stages: (i) Absynte generates reference score by performing BLASTp of a query protein sequence against itself (ii) Translation of the selected chromosome into 6 frames using TBLASTN algorithm and similarity search of the query protein sequence (s) against it with ‘bit score at default parameters’ (iii) bit score normalization using reference score (obtained in step i) and additional chromosomal ranking based on decreasing score (iv) Further, proteins derived from high ranking chromosomes were compared to each other using the Smith-Waterman–Gotoh (SWG) algorithm to identify potential paralogs/duplicates **[85]**. Our Synteny analysis revealed that the gene order for the 4 proteins was conserved in at least 4 strains. Then multiple center star gene clustering topologies helped us locate the potential orthologues and could be clearly seen in Figure 7 demarcated by the identical color lying one below the other whilst the white color indicates its absence **[85]**. The individual genes are numbered as per GenBank annotations. No events of gene inversion or recombination were seen for the genes that encode our 4 finalized proteins and their gene order was conserved **[86]**.

#### B Cell Epitope Prediction

The secretive role of B cells plays an important role in maintaining humoral immunity. A B cell epitope, therefore, can trigger the antibody production and can help in vaccine designing. Hence, we used BCPred, FBCPred server and Ellipro suite to predict Continuous B cell epitopes. For 4 proteins, we got 13 epitopes with significantly good score i.e. 0.9. from both BCPred **[88]** and FBCPred. Ellipro suite was used to predict both continuous (Linear) and discontinuous (Conformational) epitopes **[89]**. The common sequences generated by at least two of the three algorithms (BCPred, FBCPred, and Ellipro) were considered to select B-cell epitopes and to build consensus antigenic peptide. Sequences predicted as B-cell epitopes by a single algorithm only were excluded from the study. To generate consensus peptide, we used the resulting epitopes were then further shortlisted based on BepiPred linear epitope prediction **[96]**, Karplus-Schulz flexibility prediction **[93]**, Chou-Fasman beta-turn prediction **[91]**, KolaskarTongaonkar antigenicity **[94]**, Emini surface accessibility prediction **[92]**, and Parker hydrophilicity prediction. Clustal Omega was used to align epitopes generated from three algorithms for each protein. The generated peptide was then considered as the final B cell epitopes **[97]**, see Table 2, Supplementary section. The discontinuous epitopes were tabulated in Table 1S, Supplementary section.

#### MHC Class I and Class II antigenic determinant prediction, Population Coverage Analysis, &Antigenicity-Toxicity Prediction

Cytotoxic T-lymphocytes (CTLs) are MHC Class I restricted Lymphocytes (CD8+T cells) that are responsible for killing the target cells infected with the virus, bacteria or protozoa **[108]**. During infection, whenever CD8+T cell encounters the MHC-I-antigen complex specific to their receptor (also known as T Cell Receptor, TCR), the naïve CTLs proliferate by undergoing several mitotic divisions followed by differentiation into the effector cells **[109]**. Here, we tried to predict the MHC Class I receptor-specific immunogenic epitopes using the Tepitool, a standalone tool of IEDB using a reference set of 27 most frequent A and B alleles and IEDB recommended the method. The promiscuous binders of 9mer length so obtained from Tepitool were then clustered using the Epitope Cluster Analysis tool to remove redundant peptides. The unique peptides were extracted manually and were mapped back to the MHC Class I Processing data. We also carried out MHC Class I processing analysis wherein the amino acid sequence of each protein was used as a query sequence. The idea behind this analysis was to identify the MHC Class I ligands that will be generated post proteasomal digestion of the protein inside the cell, travel via TAP transporter and is finally complexed with MHC receptor and presented to the T_c_. More than 2500 antigenic peptides were predicted for each protein. The list of epitopes was sorted according to the Total score which is a consensus score based on the proteasomal cleavage score, TAP transport score, and MHC binding prediction score. The top 10% peptides were extracted from the results and then common peptides were searched from MHC Class I binding prediction dataset and MHC Class I processing analysis dataset by running an in-house script. The common peptides so obtained will have a high binding score, however, not necessarily high antigenicity. Therefore, we used the IEDB MHC Class I Immunogenicity prediction tool to score peptides based on their amino acid composition and their amino acid position/order in the peptide. The peptides with the positive Immunogenicity score were selected and those with negative scores were excluded. This gave us 53 highly immunogenic peptides in total coming from 4 different proteins.

We furthered these 53 peptides for Population coverage analysis. This analysis calculates what percentage of the population that is likely to respond to a given set of epitope-based on MHC binding and T cell restriction data and HLA genotypic frequencies **[106]**. This is due to antigenic specificity towards a particular MHC molecule and MHC polymorphism across individuals that this step becomes critical for vaccine design. We prepared a .txt file with one epitope-allele combination per line. In the case of peptides exhibiting high affinity for multiple HLA alleles, we used a .txt file with each line containing a tab-separated epitope name and comma-separated allele names.

The *M. abscessus* group is commonly found to affect East Asia, India, Singapore and Okinawa, southern Australia and Africa and *M. chelonae* in China **[110]**. Therefore, for a vaccine to benefit multitudes across the globe we used World allelic frequency dataset that covers 115 countries and 21 different ethnicities grouped into 16 different geographical zones. Additionally, we also report the population coverage of the regions that are popularly reported to be affected by Mycobacteroides under study i.e. East Asia, India, Singapore, Australia, Africa, and China **[109]**, see Table 3S.

The data on individual epitope contribution toward Population Coverage was extracted and the immunogenic peptides now were sorted based on percentage contribution to population coverage in descending order. We then chose the top 4 unique peptides with the highest percent population coverage from each protein-making 16 peptides in total. These 16 peptides were then subjected to Antigenicity and Toxicity Prediction. The antigenic nature of peptides was assessed using VaxiJen with a threshold of 0.5 and Toxicity by ToxinPred**[107]**. We used a hybrid approach i.e. SVM (Swiss-Prot) + Motif based input method that searches for motifs, if any, present in peptides against motifs present in toxic peptides and would increase the SVM score by 5 if a hit was found. All the 16 peptides were found antigenic and Non-toxic to human host tissues **[107]**.

T Helper cells (T_h_) also called CD4^+^T cells to play a key role in both humoral and cell-mediated immune response **[111]**. Therefore, epitopes specific to the T_h_ Cell receptor will be a crucial part of the prophylactic and immunotherapeutic vaccine construction. Like MHC Class I epitope prediction, all 4 protein sequences were subjected to the Tepitool MHC Class II epitope prediction module and epitopes of 15mer length were obtained using the IEDB Recommended method. We used a reference set of 26 alleles (refer to List 2S, Supplementary section) with a threshold set at ≤ 10 %. The epitopes so obtained were arranged in ascending order based on a percentile rank. We used the top 10% rule that was found effective in our previous study as well **[112]** wherein we took only the top 10% epitopes of a total number of predicted peptides and subjected to MHC cluster analysis to reduce peptide redundancy. These unique peptides were then assessed for Population Coverage analysis using the IEDB server, antigenicity using VaxiJen and Peptidic Toxicity using the ToxinPred server.

#### MHC restriction and Cluster analysis of MHC alleles

The HLA molecules have distinct binding specificity. Therefore, to understand the relationships between the binding patterns of the HLA Class I and Class II molecules, we used MHC Cluster 2.0 server that cluster molecules based on their binding specificity **[113]**. The functional relationship amidst HLA Class I and Class II variants was visualized in the form of Heat Maps and Circular phylogenetic trees.

We clustered 27 Reference set of A and B HLA Alleles (HLA-A01:01, HLA-A02:01, HLA-A02:03, HLA-A02:06, HLA-A03:01, HLA-A11:01, HLA-A23:01, HLA-A24:02, HLA-A26:01, HLA-A30:01, HLA-A30:02, HLA-A31:01, HLA-A32:01, HLA-A33:01, HLA-A68:01, HLA-A68:02, HLA-B07:02, HLA-B08:01, HLA-B15:01, HLA-B35:01, HLA-B40:01, HLA-B44:02, HLA-B44:03, HLA-B51:01, HLA-B53:01, HLA-B57:01, HLA-B58:01) (Figure 8 and 15 reference set of DP, DQ and DR alleles out of 26 alleles we used in our analysis (DRB1:0101, DRB1:0301, DRB1:0401, DRB1:0405, DRB1:0701, DRB1:0802, DRB1:0901, DRB1:1101, DRB1:1201, DRB1:1302, DRB1:1501, DRB3:0101, DRB3:0202, DRB4:0101, DRB5:0101) (Figure 9). This is because for rest of the 11 Class II molecules, complete protein sequences and structures are not available.

**Figure 9:**
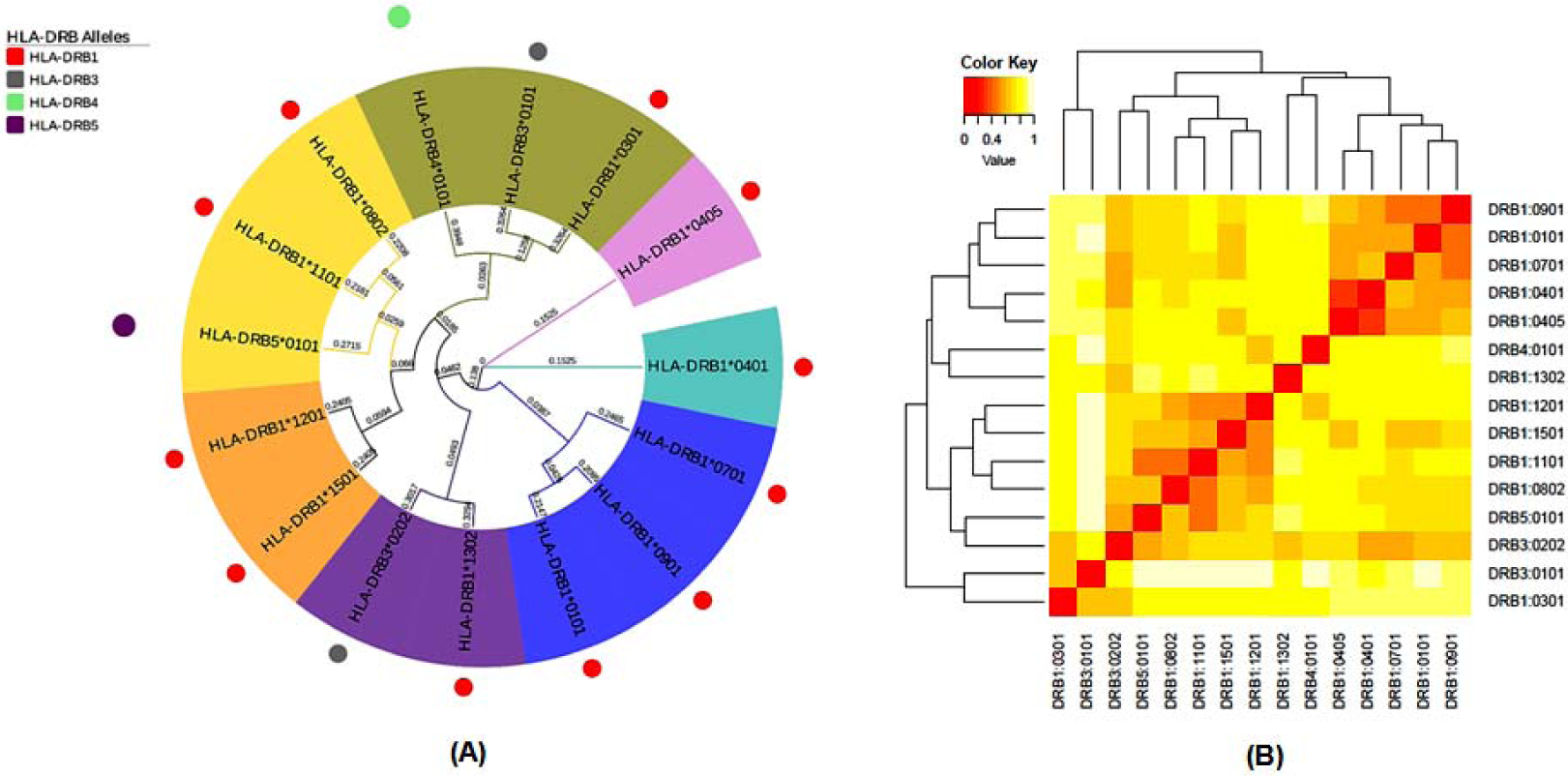
Functional relatedness of the HLA alleles for a reference set of 15 MHC Class II molecules on the basis of binding specificity (A) Represented the cluster analysis using Circular Phylogenetic tree for HLA-DRB alleles at a functional level (B) Represented using Heat Map. The Epitopes were clustered based on interaction with HLA molecules. In the heat map, the red color demarcates strong interaction and the Yellow region indicates the weaker interaction.

#### B cell Epitopes Mapping on T cell Epitopes

Out of 16 peptides finalized from MHC Class, I binding and 18 peptides finalized from MHC Class II binding prediction, we took only those peptides that had regions overlapping with the B cell epitopes. This is to lessen the number of linkers we would use to join peptides during chimeric vaccine construction. In case of overlapping regions detected, we manually summed the B Cell, Class I and Class II epitopes for all 4 proteins to generate a consensus peptide that would be joined end to end in further steps using linkers.

#### Multiepitope Vaccine Construction and Allergenicity Assessment

To construct the sequence of a final chimeric subunit vaccine, we adopted the approach as mentioned in Figure 1. The consensus B-cell epitopes were treated as a template for selecting MHC Class I and Class II epitopes and those epitopes whose sequences were found overlapping with B-cell epitopes were selected for final vaccine constructs. We found 10 MHC Class I and 11 MHC Class II peptides meeting the above criteria. The overlapping peptides were joined, and the resulting peptides were tabulated as Table 6S, Supplementary section. The permutation and combination strategy used to make 8 vaccine constructs are also represented in Table 6S, Supplementary section.

An Immunologic adjuvant is an ingredient of a vaccine that potentiates the immunogenicity of an antigen and helps improve immune response in the patient **[131]**. Different adjuvants were used in the vaccine sequence construction to evaluate the effect they have on the antigenicity and allergenicity of the constructs (For signal peptides and adjuvants used in the present study, refer to List 7S, Supplementary section). PADRE sequence along with signal peptides was also added to subside the problems presented by polymorphism of HLA-DR molecules in the population worldwide and enhance Helper T-cell response respectively **[131]** (Fig 5S).

A total of 8 Vaccine constructs were prepared and were subjected to Antigenicity prediction using AntigenPro and VaxiJen and Allergenicity using the AlgPred Prediction server. The AlgPred server was deployed to assess the allergenicity of the vaccine constructs present if any **[119]**. The hybrid approach of AlgPred uses five approaches (IgEepitope+ARPs BLAST+MAST+SVM) to conclusively predict the allergic nature of peptide with the overall accuracy of 85%. SVM Score with a threshold of −0.4 was considered as a benchmark. The constructs were predicted to be non-allergic if the predicted score was less than −0.4 (consider the -ve sign). Two vaccine constructs out of 8 were found to be allergic and were excluded from the study. The remaining 6 vaccine construct were carried forward for structure prediction (Table 7S, Supplementary section).

AntigenPro is a non-pathogen specific, alignment free-sequence based predictor of Antigenicity that predicts antigenicity with an accuracy of 76% **[100, 131]**. The antigenicity of the shortlisted 6 vaccine constructs was further confirmed using VaxiJen v2.0 that predicts antigenicity with an accuracy of 70-89% (82% for bacterial antigens). The predicted antigenicity values were more than 0.6 for V4, V5, V6 vaccine constructs as predicted by AntigenPro and more than 0.75 in VaxiJen v2.0. V6 had the highest VaxiJen score while V2 the lowest. V4 and V5 constructs had the same score in AntigenPro **[129]** followed by V6 in the line. Thus V4, V5, and V6 show good antigenic characteristics out of the 6 constructs. However, we considered working with all 6 constructs until the end to find the best one out (Table 7S Supplementary section).

We also predicted the propensity of the vaccine to be soluble upon over-expression in E. coli using SOLPro. It uses Two-stage Support Vector Machine (SVM) and based on multiple representations of the primary amino acid sequence predicts the protein/peptide solubility with an accuracy of 76% **[130]**. This analysis was carried out to identify which vaccine constructs would pose a problem from a solubility standpoint and to prioritize vaccine constructs when carrying out production on a large scale. All the vaccine constructs were found to be readily soluble with a score greater than 0.8 (Table 7S Supplementary section).

#### Prediction of Secondary Structure and Physicochemical Properties of Vaccine Constructs

PSIPRED v3.3 server was used to predict the secondary structure of the chimeric vaccine constructs. To find the intrinsically disordered regions present in the sequences of the constructs, we used DISOPRED 2 prediction method **[80]**. The tertiary structures prediction of vaccines was carried out using I-TASSER that employs an integrative modeling approach including protein threading, Iterative structure assembly, and comparative homology modeling **[70]**. The structures so obtained were then subjected to protein structure validation and refinement using the pipeline mentioned in Fig 4S. For structure refinement, we used ModRefine. It uses a two-step refinement algorithm: The first step being low-resolution refinement that involves building the initial backbone atoms using a look-up table for the C_a_ trace followed by energy minimization simulation for backbone quality refinement; The second being the high-resolution refinement that involves the addition of side-chain atoms from a rotamer library followed by a fast energy minimization step for the refinement of both side-chain conformations and backbone conformations **[72]**. The physicochemical characterization was done by the XtalPred-RF server and the characteristics are charted in the Table 4S, Supplementary section.

#### Local Frustration Analysis

Regions of Local frustrations in the refined models were identified using Frustratometer 2 that deploy an Algorithm based on Energy Landscape theory and quantify the degree of local frustration in the input protein model. The highly frustrated regions were more prominent in vaccine constructs V1, V3, and V6 and subtle in V2, V4, and V5 (data not shown). These regions can have a crucial biological function such as interaction with HLA proteins and/or with TLR4-MD2 complex which we tried to probe into after docking in V6 (Fig 9S, Supplementary section).

#### Disulfide bond Engineering and Docking

The knowledge garnered from Intrinsically Disordered region prediction was used for the stabilization of Vaccine constructs via Disulfide bond engineering (data not are shown). We left out those disordered regions that were predicted to be involved in Binding. We created mutations in not more than 3 positions in all vaccine constructs using DbD v2.12. In V1, the mutant SS bond were created at 54-289, 198-312, 389-419 residue, V2 at 270-293, 299-309, 314-319 residue, V3 at 95-102, 190-305, 215-218 residue, V4 at 91-443, 197-202, 227-265 residue, V5 at 134-158, 148-151, 260-261 residue and V6 at 196-199, 212-214, 334-335 residue.

To determine the binding affinity of the resulting vaccine constructs with the HLA molecules, we carried out Protein-Protein docking post protein backbone stabilization. The HLA molecules whose crystal structures had missing strings of residues were excluded from the study and only those with complete crystal structure were considered for docking. We manually preprocessed the HLA proteins and reduced homodimers to monomers to lessen the docking time. Ions, ligands and other non-amino acid molecules were also removed using the Chimera Visualization tool. Rigid docking of each vaccine constructs was then carried out with HLA proteins using PatchDock and the poses so generated were sorted with respect to the Binding Energy function. The top 10 docking outcomes were pulled and furthered for pose refinement using FireDock. For each HLA molecule, we compared the Binding energy of the complexes and those with the lowest Binding energies are highlighted.

Since our Chimeric vaccine’s constructs contained both Linear B-cell epitopes and T-cell epitopes they can bind to HLA molecules and successfully initiate both humoral and cellular immune response. However, immune response evoked against the epitopes might be genetically restricted i.e. epitopes may evoke an immune response in one individual but not be in other. Hence, a vaccine construct against Mycobacteroides should be such that it can potentiate the immune response against the number of epitopes and can be recognized by multiple HLA allele’s proteins. The promiscuous vaccine is successful against a population with polymorphic HLA molecules. We docked all six vaccine constructs with the six different HLA allele’s protein. Vaccine construct 6 (V6) had lowest global binding energy with different HLA alleles HLA-DR B1*03:01 (−17.15), HLA-DR B3*02:02 (−23.86), HLA-DR B5*01:01 (−7.07), HLA-B*35-01 (−12.64), HLA-A*11:01 (−25.31), HLA-B*51:01 (−20.79), HLA-A*24:02 (−20.18) and has been tabulated in Table 5S, Supplementary section. Based on the different parameters, we analyzed all six chimeric constructs and finalized one suitable and best vaccine construct i.e. V6 that can control Mycobacteroides infection. Moreover, the presence of β defensin in V6 can act as chemo-attractants for resting T cells and help in the recruitment of various Antigen Presenting Cells (APC) thereby modulating host’s adaptive immunity **[132]**.

β defensin can act as a ligand for TLR4 and facilitate Dendritic Cells (DCs) Maturation (or Antigen Presenting Cells, APCs) **[133]**. TLR4 is agonist to β defensin. Therefore, to study the interaction between V6 and TLR4/ MD2 complex (PDB ID: 4G8A; processed manually before docking) we carried out protein docking via PatchDock server and Pose refinement with FireDock. The result showed that the global binding energy of the complex was −0.22 kcal/mol that showed good interaction between V6 and TLR4/ MD2 complex. We ran MD-simulation on the complex for 150 ns using GROMACS v5.1.2 **[136]** PLS-AA/L all-atom force field (2001 amino acid dihedrals). The complex was solvated in a dodecahedron water box using a four-point TIP4P rigid water **[135]** model with at least 1 nm of solvation on all sides (Avg. volume: 786.44 nm^3^) and neutralized by adding Na^+^ ions. We used Particle Mesh Ewald (PME) summation method for the treatment of long-range interactions with all bonds constrained using the LINCS algorithm. Further, the energy minimization of the system was carried out using a steepest descent method. We ran simulations at 310 K temperature and 1 bar atmospheric pressure via V-rescale thermostat and Parrinello-Rahman barostat implementation. The conformations were obtained at an interval of 20 ps throughout the 150ns trajectory. Post-simulation, energy minimization, and trajectory analysis, we found that complex showed 2 Å deviation initially but got stabilized later (Figure 10 A). The radius of gyration trajectory obtained for the docked complex showed that the distance in the rotating complex from Centre of Mass (COM) was initially around 3.7 but dropped to 3.3 nm during the due course of the simulation (Figure 10 B). At the end of the simulation, the RMSF of Chain A of V6 was 0.5 to 1Å, that of Chain B (TLR4) was about 0.5 Å and that of Chain D was below 0.25 Å as visible in the RMSF Plots. The scores were satisfactory demarcating strong complex stability. The interacting interfacial residues were determined using PRODIGY Server **[137]**and the Binding energy (ΔG) was estimated to be -15.2 kcal mol^-1^ with dissociation constant (K_d_). Number of Interfacial Contacts (ICs) were as follows: ICs charged-charged: 16, ICs charged-polar: 15, ICs charged-apolar: 30, ICs polar-polar: 7, ICs polar-apolar: 30, ICs apolar-apolar: 11. The charged but Non-Interacting Surface (NIC) was 23.67% and the NIC apolar was 39.68%. The list of interacting residues is attached in the List 8S, Supplementary section. The objective of the *in-silico* cloning was the heterologous expression of the multiepitope vaccine as a protein product using *E. coli* as an expression system.

**Figure 10:**
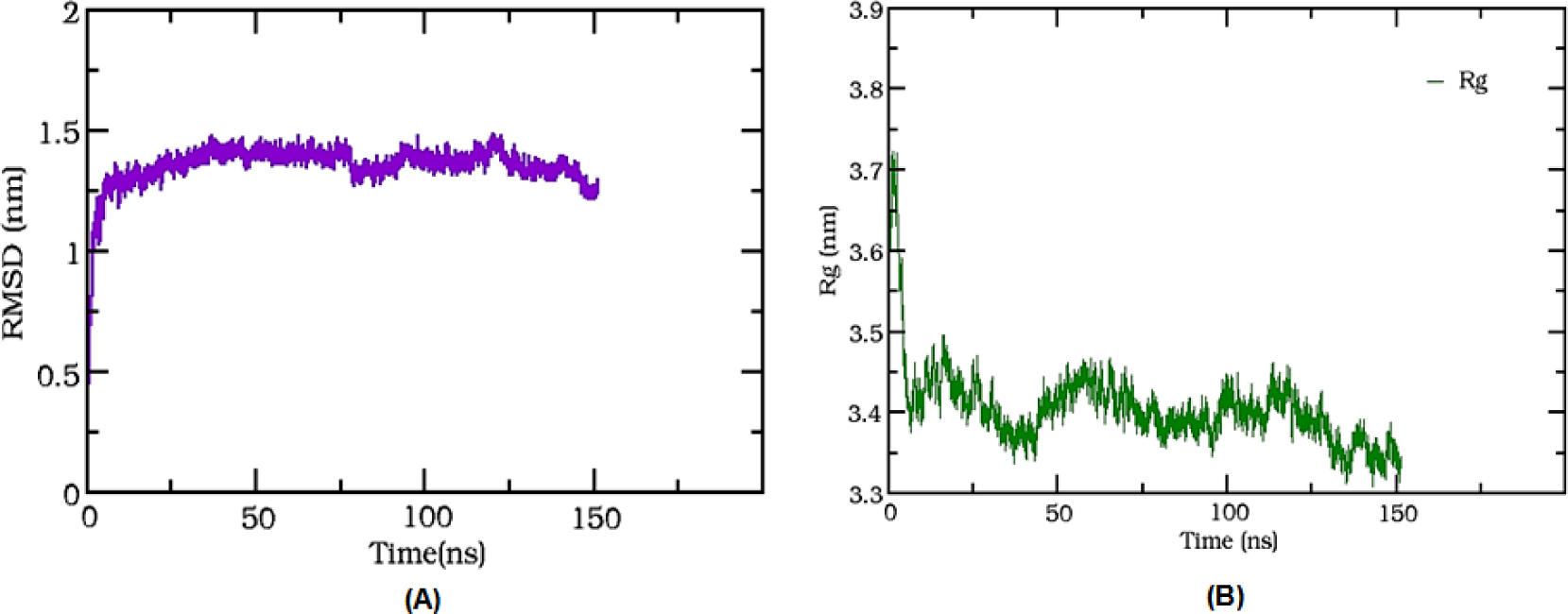
MD-Simulation of V6-TLR4/MD2 complex and RMSD/ RMSF trajectories. (A) Shows the RMSD post complex stabilization at 1.5 nm. (B) Shows the Radius of Gyration stabilizes and drops from 3.7 to 3.3 reflecting complex stability.

#### Codon Optimization for Restriction Cloning

Codon optimization is a technique that improves protein translational efficiency in the expression system of an organism. It is, therefore, necessary to first back-translated the amino acid sequence into nucleotide sequence using the precomputed codon optimization table for *Escherichia coli* str. K12 substr. MG1655. We applied the Guided Random method (Monte Carlo algorithm) in OPTIMIZER (http://genomes.urv.es/OPTIMIZER/) ran the back translation **[134]**. The nucleotide sequence so obtained had the Codon Adaption Index (CAI) of 0.712 with 59.7% GC content and 40.3% AT content. However, the sequence contained multiple Type II Restriction sites. To remove them we used the services of the IDT Codon Optimization Tool that use a codon sampling strategy for CAI improvement. The of Restriction sites were observed to decrease significantly from 24 to 18. The Total Complexity Score was 0.4/7. The GC content in the optimized sequence was 58.4%. Post codon optimization we added AGCT adaptors at both ends of the sequence to create sticky ends for insertion between the Hind3 and Xho1 in the *E coli pET28a*(+) vector DNA. This was achieved using SnapGene software **[138]**. The sequence that code for vaccine V6 along with adapter sequences was 1103 bp long the insertion process of which is shown in Figure 11. The total length of the cloned gene was 6.4kbp. For sequence, refer List 9S, Supplementary section.

**Figure 11:**
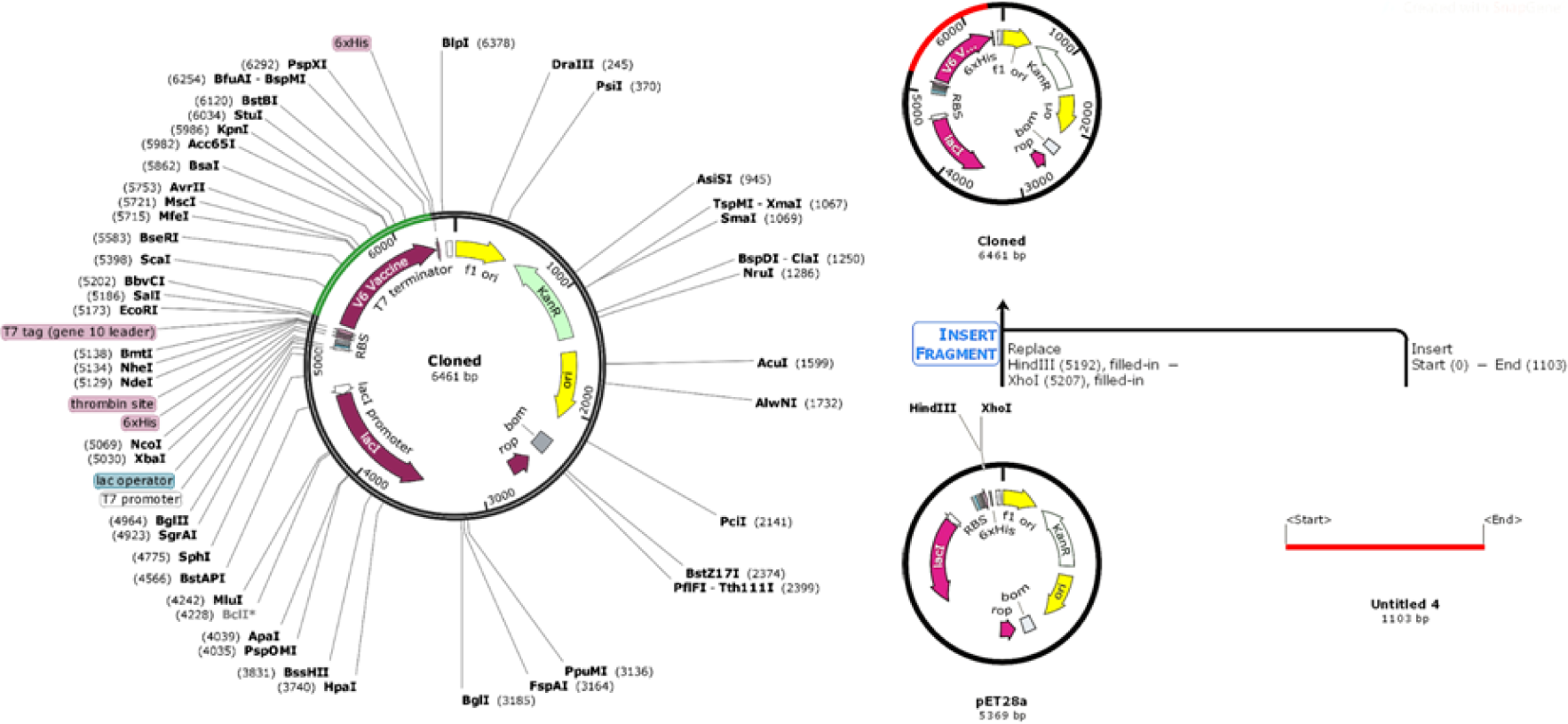
*In-silico* Restriction digestion of pET28a (+) expression vector using SnapGene and cloning of gene sequence of final vaccine construct V6 into the vector between Hind III (5192) and XhoI (5207). The vector plasmid of *E. coli* (left) is inserted with the Vaccine coding gene (Green). The mechanism of insertion is highlighted in the right (Vaccine coding gene in red).

## Conclusion

The treatment regimens to combat bacterial infection are failing at large due to the spread of antimicrobial resistance **[32,33]**. Vaccination on the other side could prove to be an effective and affordable measure to improve public health and prevent individuals from NTM’s **[98, 99, 100]**. The genomic and proteomic study on 5 strains of Mycobacteroides synthesized enormous information that can be finally put for Vaccine Development and/or Drug Development by the scientific community. Our genomic analysis underlines the events of deletion in strains MM1513, CCUG 47445, and MOTT36W. The genome of the Mycobacteroides was found to be populated with pseudogenes with a maximum in MM 1513 strain. The presence of antimicrobial-resistant genes followed by Pan-Core analysis, Prophage Prediction and CRISPR-Cas system detection provided us further details that the Pan-genome of the organism was open but shall close soon. Further, the Subtractive Proteomics Approach helped us identify antigenic human Non-homologues, Reverse Vaccinology approach helped us in chimeric vaccine construction and the best candidates were identified after protein-protein docking studies. The final vaccine construct was back-translated for the purpose of restriction cloning. This vaccine could open new therapeutic avenues to combat this egregious bacterium.

## Supporting information

Supplementary section

## Author Contributions

The present study was conceived by Rohit Satyam. The Methodology designing and the analysis was jointly carried out by Dr Niraj and Dr Tulika the execution of which was carried out by Rohit Satyam. The data validation was taken care by Dr Tulika and Rohit Satyam and Dr Niraj the manuscript was structured and scrutinized by Dr Niraj.

## Acknowledgments

The authors would like to express their sincere gratitude to Noida Institute of Engineering and Technology for providing necessary infrastructure. Additionally, we would like to thanks Dr Surabhi Rathore for sharing her expertise in the MD Simulation and helping us validate the simulation data.

## References

1. Gupta, R.S., Lo, B. and Son, J., 2018. Phylogenomics and comparative genomic studies robustly support division of the genus Mycobacterium into an amended genus Mycobacterium and four novel genera. Frontiers in Microbiology, 9, p.67.

2. Brown-Elliott, B.A. and Wallace, R.J., 2002. Clinical and taxonomic status of pathogenic non-pigmented or late-pigmenting rapidly growing mycobacteria. Clinical microbiology reviews, 15(4), pp.716–746.

3. Gonzalez-Diaz, E., Morfin-Otero, R., Perez-Gomez, H.R., Esparza-Ahumada, S. and Rodriguez-Noriega, E., 2018. Rapidly Growing Mycobacterial Infections of the Skin and Soft Tissues Caused by M. fortuitum and M. chelonae. Current Tropical Medicine Reports, pp.1–8.

4. Runyon, E.H., Wayne, L.G. and Kubica, G.P., 1974. Genus I. Mycobacterium Lehmann and Neumann 1896, 363. Bergey’s Manual of Determinative Bacteriology, Eighth Edition, The Williams and Wilkins Co., Baltimore, pp.682–701.

5. Nessar, R., Cambau, E., Reyrat, J.M., Murray, A. and Gicquel, B., 2012. Mycobacterium abscessus: a new antibiotic nightmare. Journal of antimicrobial chemotherapy, 67(4), pp.810–818.

6. Bryant, J.M., Grogono, D.M., Rodriguez-Rincon, D., Everall, I., Brown, K.P., Moreno, P., Verma, D., Hill, E., Drijkoningen, J., Gilligan, P. and Esther, C.R., 2016. Emergence and spread of a human-transmissible multidrug-resistant nontuberculous mycobacterium. Science, 354(6313), pp.751–757.

7. Churgin, D.S., Tran, K.D., Gregori, N.Z., Young, R.C., Alabiad, C. and Flynn Jr, H.W., 2018. Multi-drug resistant Mycobacterium chelonae scleral buckle infection. American journal of ophthalmology case reports, 10, pp.276–278.

8. Runyon, E.H., 1959. Anonymous mycobacteria in pulmonary disease. Medical Clinics of North America, 43(1), pp.273–290.

9. Song, Y., Zhang, L., Yang, H., Liu, G., Huang, H., Wu, J. and Chen, J., 2018. Nontuberculous mycobacterium infection in renal transplant recipients: a systematic review. Infectious Diseases, 50(6), pp.409–416.

10. Svetlíková, Z., Škovierová, H., Niederweis, M., Gaillard, J.L., McDonnell, G. and Jackson, M., 2009. Role of porins in the susceptibility of Mycobacterium smegmatis and Mycobacterium chelonae to aldehyde-based disinfectants and drugs. Antimicrobial agents and chemotherapy, 53(9), pp.4015–4018.

11. Moorthy, R.S., Valluri, S. and Rao, N.A., 2012. Nontuberculous mycobacterial ocular and adnexal infections. Survey of Ophthalmology, 57(3), pp.202–235.

12. Nascimento, H., Viana-Niero, C., Nogueira, C.L., Bispo, P.J.M., Pinto, F., Uzam, C.D.P.P., Matsumoto, C.K., Machado, A.M.O., Leão, S.C., Höfling-Lima, A.L. and de Freitas, D., 2018. Identification of the infection source of an outbreak of Mycobacterium chelonae keratitis after laser in situ keratomileuses. Cornea, 37(1), pp.116–122.

13. Sander, M.A., Isaac-Renton, J.L. and Tyrrell, G.J., 2018. Cutaneous Nontuberculous Mycobacterial Infections in Alberta, Canada: An Epidemiologic Study and Review. Journal of cutaneous medicine and surgery, p.1203475418776945.

14. Verghese, E., Jabr, A. and Watkins, R.R., 2018, August. Successful treatment of a renal abscess caused by Mycobacterium chelonae: A case report. In Open forum infectious diseases (Vol. 5, No. 9, p. ofy196). US: Oxford University Press.

15. Hashimoto, Y., Ikeda, A., Tokuyasu, Y., Omura, H. and Tanaka, T., 2018. Cutaneous Mycobacterium chelonae infection following autologous peripheral blood stem cell transplantation for POEMS syndrome. Journal of Infection and Chemotherapy.

16. Tejura, N., Bontempo, G. and Chew, D., 2018, August. Disseminated Mycobacterium abscessus infection secondary to an infected vascular stent: Case Report and Review of the Literature. In Open forum infectious diseases (Vol. 5, No. 9, p. ofy207). US: Oxford University Press.

17. Kennedy, B.S., Bedard, B., Younge, M., Tuttle, D., Ammerman, E., Ricci, J., Doniger, A.S., Escuyer, V.E., Mitchell, K., Noble-Wang, J.A. and O’Connell, H.A., 2012. The outbreak of Mycobacterium chelonae infection associated with tattoo ink. New England Journal of Medicine, 367(11), pp.1020–1024.

18. Correa, N.E., Cataño, J.C., Mejía, G.I., Realpe, T., Orozco, B., Estrada, S., Vélez, A., Vélez, L., Barón, P., Guzmán, A. and Robledo, J., 2010. The outbreak of mesotherapy-associated cutaneous infections caused by Mycobacterium chelonae in Colombia. Jpn J Infect Dis, 63(2), pp.143–145.

19. Griffith, D.E., Aksamit, T., Brown-Elliott, B.A., Catanzaro, A., Daley, C., Gordin, F., Holland, S.M., Horsburgh, R., Huitt, G., Iademarco, M.F. and Iseman, M., 2007. An official ATS/IDSA statement: diagnosis, treatment, and prevention of nontuberculous mycobacterial diseases. American journal of respiratory and critical care medicine, 175(4), pp.367–416.

20. Takemori□Sakai, Y., Iwata, Y., Oe, H., Sakai, Y. and Wada, T., 2018. Bloodstream infection caused by Mycobacterium chelonae. Pediatrics International.

21. Williams, M.M., Yakrus, M.A., Arduino, M.J., Cooksey, R.C., Crane, C.B., Banerjee, S.N., Hilborn, E.D. and Donlan, R.M., 2009. Structural analysis of biofilm formation by rapidly and slowly growing nontuberculous mycobacteria. Applied and environmental microbiology, 75(7), pp.2091–2098.

22. Updated 2017 May 16]. In: StatPearls [Internet]. Treasure Island (FL): StatPearls Publishing; 2018 Jan-. Available from: https://www.ncbi.nlm.nih.gov/books/NBK430806/

23. Mannelli, V.K., Rai, M.P., Nemakayala, D.R. and Kadiri, N.P., 2018. Mycobacterium chelonae Developing Multidrug Resistance. BMJ case reports, 2018

24. Nash, K.A., Brown-Elliott, B.A. and Wallace, R.J., 2009. A novel gene, erm (41), confers inducible macrolide resistance to clinical isolates of Mycobacterium abscessus but is absent from Mycobacterium chelonae. Antimicrobial agents and chemotherapy, 53(4), pp.1367–1376.

25. Browne, S.K., Burbelo, P.D., Chetchotisakd, P., Suputtamongkol, Y., Kiertiburanakul, S., Shaw, P.A., Kirk, J.L., Jutivorakool, K., Zaman, R., Ding, L. and Hsu, A.P., 2012. Adult-onset immunodeficiency in Thailand and Taiwan. New England Journal of Medicine, 367(8), pp.725–734.

26. van Ingen, J., 2019. Drug Susceptibility Testing of Nontuberculous Mycobacteria. In Nontuberculous Mycobacterial Disease (pp. 61–88). Humana Press, Cham.

27. Daley, C.L. and Griffith, D.E., 2010. Pulmonary non-tuberculous mycobacterial infections. International Journal of Tuberculosis and Lung Disease, 14(6), pp.665–671.

28. Griffith, D.E. and Aksamit, T.R., 2012. Therapy of refractory nontuberculous mycobacterial lung disease. Current opinion in infectious diseases, 25(2), pp.218–227.

29. Nie, W., Duan, H., Huang, H., Lu, Y., Bi, D. and Chu, N., 2014. Species identification of Mycobacterium abscessus subsp. abscessus and Mycobacterium abscessus subsp. bolletii using rpoB and hsp65, and susceptibility testing to eight antibiotics. International Journal of Infectious Diseases, 25, pp.170–174.

30. Koh, W.J., Jeon, K., Lee, N.Y., Kim, B.J., Kook, Y.H., Lee, S.H., Park, Y.K., Kim, C.K., Shin, S.J., Huitt, G.A. and Daley, C.L., 2011. Clinical significance of differentiation of Mycobacterium massiliense from Mycobacterium abscessus. American journal of respiratory and critical care medicine, 183(3), pp.405–410.

31. Harada, T., Akiyama, Y., Kurashima, A., Nagai, H., Tsuyuguchi, K., Fujii, T., Yano, S., Shigeto, E., Kuraoka, T., Kajiki, A. and Kobashi, Y., 2013. Clinical and Microbiological Differences between Mycobacterium abscessus and Mycobacterium massiliense Lung Diseases. Journal of clinical microbiology, 51(3), pp.1061–1061.

32. Story-Roller, E., Maggioncalda, E.C., Cohen, K.A. and Lamichhane, G., 2018. Mycobacterium abscessus and β-lactams: emerging insights and potential opportunities. Frontiers in Microbiology, 9.

33. Philley, J.V. and Griffith, D.E., 2019. Disease Caused by Mycobacterium Abscessus and Other Rapidly Growing Mycobacteria (RGM). In Nontuberculous Mycobacterial Disease (pp. 369–399). Humana Press, Cham.

34. Zheng, J., Lin, X., Wang, X., Zheng, L., Lan, S., Jin, S., Ou, Z. and Wu, J., 2017. In silico analysis of epitope-based vaccine candidates against the hepatitis B virus polymerase protein. Viruses, 9(5), p.112.

35. Hubbard, A.T., Davies, S.E., Baxter, L., Thompson, S., Collery, M.M., Hand, D.C., Thomas, D.J.I. and Fink, C.G., 2018. Comparison of the first whole genome sequence of ‘Haemophilusquentini’ with two new strains of ‘Haemophilusquentini’ and other species of Haemophilus. Genome, 61(5), pp.379–385.

36. Darling, A.C., Mau, B., Blattner, F.R. and Perna, N.T., 2004. Mauve: multiple alignments of conserved genomic sequence with rearrangements. Genome research, 14(7), pp.1394–1403.

37. Lee, I., Kim, Y.O., Park, S.C. and Chun, J., 2016. OrthoANI: an improved algorithm and software for calculating average nucleotide identity. International journal of systematic and evolutionary microbiology, 66(2), pp.1100–1103.

38. Zankari, E., Hasman, H., Cosentino, S., Vestergaard, M., Rasmussen, S., Lund, O., Aarestrup, F.M. and Larsen, M.V., 2012. Identification of acquired antimicrobial resistance genes. Journal of antimicrobial chemotherapy, 67(11), pp.2640–2644.

39. Chaudhari, N.M., Gupta, V.K. and Dutta, C., 2016. BPGA-an ultra-fast pan-genome analysis pipeline. Scientific reports, 6, p.24373.

40. Arndt, D., Grant, J.R., Marcu, A., Sajed, T., Pon, A., Liang, Y. and Wishart, D.S., 2016. PHASTER: a better, faster version of the PHAST phage search tool. Nucleic acids research, 44(W1), pp.W16–W21.

41. Couvin, D., Bernheim, A., Toffano-Nioche, C., Touchon, M., Michalik, J., Néron, B., Rocha, C., Eduardo, P., Vergnaud, G., Gautheret, D. and Pourcel, C., 2018. CRISPRCasFinder, an update of CRISRFinder, includes a portable version, enhanced performance and integrates search for Cas proteins. Nucleic acids research.

42. Negahdaripour, M., Nezafat, N., Hajighahramani, N., Rahmatabadi, S.S. and Ghasemi, Y., 2017. Investigating CRISPR-Cas systems in Clostridium botulinum via bioinformatics tools. Infection, Genetics, and Evolution, 54, pp.355–373.

43. Biswas, A., Staals, R.H., Morales, S.E., Fineran, P.C. and Brown, C.M., 2016. CRISPRDetect: a flexible algorithm to define CRISPR arrays. BMC Genomics, 17(1), p.356.

44. Carver, T., Harris, S.R., Berriman, M., Parkhill, J. and McQuillan, J.A., 2011. Artemis: an integrated platform for visualization and analysis of high-throughput sequence-based experimental data. Bioinformatics, 28(4), pp.464–469.

45. Benson, G., 1999. Tandem repeats finder: a program to analyze DNA sequences. Nucleic acids research, 27(2), pp.573–580.

46. Singh, P. and Cole, S.T., 2011. Mycobacterium leprae: genes, pseudogenes and genetic diversity. Future Microbiology, 6(1), pp.57–71

47. Gómez-Valero, L., Rocha, E.P., Latorre, A. and Silva, F.J., 2007. Reconstructing the ancestor of Mycobacterium leprae: the dynamics of gene loss and genome reduction. Genome research, 17(8), pp.1178–1185.

48. Zhou, K., Aertsen, A., and Michiels, C.W., 2014. The role of variable DNA tandem repeats in bacterial adaptation. FEMS microbiology reviews, 38(1), pp.119–141.

49. Sarkar, M., Maganti, L., Ghoshal, N. and Dutta, C., 2012. In the silico quest for putative drug targets in Helicobacter pylori HPAG1: molecular modeling of candidate enzymes from lipopolysaccharide biosynthesis pathway. Journal of molecular modeling, 18(5), pp.1855–1866.

50. Shanmugham, B. and Pan, A., 2013. Identification and characterization of potential therapeutic candidates in emerging human pathogen Mycobacterium abscessus: a novel hierarchical in silico approach. PloS one, 8(3), p.e59126.

51. Luo, H., Lin, Y., Gao, F., Zhang, C.T. and Zhang, R., 2013. DEG 10, an update of the database of essential genes that includes both protein-coding genes and noncoding genomic elements. Nucleic acids research, 42(D1), pp.D574–D580.

52. Shende, G., Haldankar, H., Barai, R.S., Bharmal, M.H., Shetty, V. and Idicula-Thomas, S., 2016. PBIT: Pipeline Builder for Identification of Drug Targets for infectious diseases. Bioinformatics, 33(6), pp.929–931.

53. Hughes, J.D., Blagg, J., Price, D.A., Bailey, S., DeCrescenzo, G.A., Devraj, R.V., Ellsworth, E., Fobian, Y.M., Gibbs, M.E., Gilles, R.W. and Greene, N., 2008. Physiochemical drug properties associated with in vivo toxicological outcomes. Bioorganic & medicinal chemistry letters, 18(17), pp.4872–4875

54. Quigley, E.M., 2013. Gut bacteria in health and disease. Gastroenterology & Hepatology, 9(9), p.560.

55. Patel, R.M. and Lin, P.W., 2010. Developmental biology of gut-probiotic interaction. Gut microbes, 1(3), pp.186–195.

56. Barh, D., Tiwari, S., Jain, N., Ali, A., Santos, A.R., Misra, A.N., Azevedo, V. and Kumar, A., 2011. In silico subtractive genomics for target identification in human bacterial pathogens. Drug Development Research, 72(2), pp.162–177.

57. Savojardo, C., Martelli, P.L., Fariselli, P., Profiti, G. and Casadio, R., 2018. BUSCA: an integrative web server to predict subcellular localization of proteins. Nucleic acids research, 46(W1), pp.W459–W466.

58. Conesa, A., Götz, S., García-Gómez, J.M., Terol, J., Talón, M. and Robles, M., 2005. Blast2GO: a universal tool for annotation, visualization, and analysis in functional genomics research. Bioinformatics, 21(18), pp.3674–3676.

59. Zhou, M., Boekhorst, J., Francke, C. and Siezen, R.J., 2008. LocateP: a genome-scale subcellular-location predictor for bacterial proteins. BMC Bioinformatics, 9(1), p.173.

60. Sharma, A.K., Dhasmana, N., Dubey, N., Kumar, N., Gangwal, A., Gupta, M. and Singh, Y., 2017. Bacterial virulence factors: secreted for survival. Indian journal of microbiology, 57(1), pp.1–10.

61. Mora, M., Donati, C., Medini, D., Covacci, A. and Rappuoli, R., 2006. Microbial genomes and vaccine design: refinements to the classical reverse vaccinology approach. Current opinion in microbiology, 9(5), pp.532–536.

62. Mushegian, A.R. and Koonin, E.V., 1996. A minimal gene set for cellular life derived by comparison of complete bacterial genomes. Proceedings of the National Academy of Sciences, 93(19), pp.10268–10273.

63. Hooper, L.V., Bry, L., Falk, P.G. and Gordon, J.I., 1998. Host-microbial symbiosis in the mammalian intestine: exploring an internal ecosystem. Bioessays, 20(4), pp.336–343.

64. Zhou, C.E., Smith, J., Lam, M., Zemla, A., Dyer, M.D. and Slezak, T., 2006. MvirDB—a microbial database of protein toxins, virulence factors and antibiotic resistance genes for bio-defense applications. Nucleic acids research, 35(Suppl_1), pp.D391–D394.

65. Garg, A. and Gupta, D., 2008. VirulentPred: an SVM based prediction method for virulent proteins in bacterial pathogens. BMC Bioinformatics, 9(1), p.62.

66. Doytchinova, I.A., and Flower, D.R., 2007. VaxiJen: a server for prediction of protective antigens, tumor antigens and subunit vaccines. BMC Bioinformatics, 8(1), p.4.

67. Sarkar, M., Maganti, L., Ghoshal, N. and Dutta, C., 2012. In silico quest for putative drug targets in Helicobacter pylori HPAG1: molecular modeling of candidate enzymes from lipopolysaccharide biosynthesis pathway. Journal of molecular modeling, 18(5), pp.1855–1866.

68. Slabinski, L., Jaroszewski, L., Rychlewski, L., Wilson, I.A., Lesley, S.A. and Godzik, A., 2007. XtalPred: a web server for prediction of protein crystallizability. Bioinformatics, 23(24), pp.3403–3405.

69. Huang, Y., Niu, B., Gao, Y., Fu, L. and Li, W., 2010. CD-HIT Suite: a web server for clustering and comparing biological sequences. Bioinformatics, 26(5), pp.680–682.

70. Zhang, Y., 2008. I-TASSER server for protein 3D structure prediction. BMC Bioinformatics, 9(1), p.40.

71. Gunasekaran, K., Ramakrishnan, C. and Balaram, P., 1996. Disallowed Ramachandran conformations of amino acid residues in protein structures. Journal of molecular biology, 264(1), pp.191–198.

72. Xu, D. and Zhang, Y., 2011. Improving the physical realism and structural accuracy of protein models by a two-step atomic-level energy minimization. Biophysical Journal, 101(10), pp.2525–2534.

73. Wiederstein, M. and Sippl, M.J., 2007. ProSA-web: interactive web service for the recognition of errors in three-dimensional structures of proteins. Nucleic acids research, 35(Suppl_2), pp.W407–W410.

74. Wallner, B. and Elofsson, A., 2003. Can correct protein models be identified?. Protein Science, 12(5), pp.1073–1086.

75. Araujo, F.A., Barh, D., Silva, A., Guimarães, L. and Ramos, R.T.J., 2018. GO FEAT: a rapid web-based functional annotation tool for genomic and transcriptomic data. Scientific reports, 8(1), p.1794.

76. Moreno-Hagelsieb, G., Treviño, V., Pérez-Rueda, E., Smith, T.F. and Collado-Vides, J., 2001. Transcription unit conservation in the three domains of life: a perspective from Escherichia coli. Trends in Genetics, 17(4), pp.175–177.

77. Bhardwaj, T. and Somvanshi, P., 2017. Pan-genome analysis of Clostridium botulinum reveals unique targets for drug development. Gene, 623, pp.48–62.

78. Rödelsperger, C. and Dieterich, C., 2010. CYNTENATOR: progressive gene order alignment of 17 vertebrate genomes. PloS one, 5(1), p.e8861.

79. Petersen, T.N., Brunak, S., Von Heijne, G. and Nielsen, H., 2011. SignalP 4.0: discriminating signal peptides from transmembrane regions. Nature methods, 8(10), p.785.

80. Ward, J.J., Mcguffin, L.J., Bryson, K., Buxton, B.F. and Jones, D.T., 2004. The DISOPRED server for the prediction of protein disorder. Bioinformatics, 20(13), pp.2138–2139.

81. da Costa, W.L.O., de Aragao Araujo, C.L., Dias, L.M., de Sousa Pereira, L.C., Alves, J.T.C., Araújo, F.A., Folador, E.L., Henriques, I., Silva, A. and Folador, A.R.C., 2018. Functional annotation of hypothetical proteins from the Exiguobacteriumantarcticum strain B7 reveals proteins involved in adaptation to extreme environments, including high arsenic resistance. PloS one, 13(6), p.e0198965.

82. Van Pittius, N.C.G., Sampson, S.L., Lee, H., Kim, Y., Van Helden, P.D. and Warren, R.M., 2006. Evolution and expansion of the Mycobacterium tuberculosis PE and PPE multigene families and their association with the duplication of the ESAT-6 (esx) gene cluster regions. BMC evolutionary biology, 6(1), p.95.

83. Ehlers, S., 1999. Immunity to tuberculosis: a delicate balance between protection and pathology. FEMS Immunology & Medical Microbiology, 23(2), pp.149–158.

84. Quiding-Järbrink, M., Smith, D.A. and Bancroft, G.J., 2001. Production of matrix metalloproteinases in response to mycobacterial infection. Infection and immunity, 69(9), pp.5661–5670.

85. Despalins, A., Marsit, S. and Oberto, J., 2011. Absynte: a web tool to analyze the evolution of orthologous archaeal and bacterial gene clusters. Bioinformatics, 27(20), pp.2905–2906.

86. Cossu, M., Badel, C., Catchpole, R., Gadelle, D., Marguet, E., Barbe, V., Forterre, P. and Oberto, J., 2017. Flipping chromosomes in deep-sea archaea. PLoS Genetics, 13(6), p.e1006847.

87. Sanchez-Trincado, J.L., Gomez-Perosanz, M. and Reche, P.A., 2017. Fundamentals and Methods for T-and B-Cell Epitope Prediction. Journal of immunology research, 2017.

88. El-Manzalawy, Y., Dobbs, D. and Honovar, V., 2012. BCPREDS: B-cell epitope prediction server. Artificial Intelligence Research Laboratory, Department of Computer Science, Iowa State University of Science and Technology. Available: http://ailab.cs.iastate.edu/bcpreds/. Accessed August.

89. Ponomarenko, J., Bui, H.H., Li, W., Fusseder, N., Bourne, P.E., Sette, A. and Peters, B., 2008. ElliPro: a new structure-based tool for the prediction of antibody epitopes. BMC Bioinformatics, 9(1), p.514.

90. Beaver, J.E., Bourne, P.E. and Ponomarenko, J.V., 2007. EpitopeViewer: a Java application for the visualization and analysis of immune epitopes in the Immune Epitope Database and Analysis Resource (IEDB). Immunome Research, 3(1), p.3.

91. Vita, R., Overton, J.A., Greenbaum, J.A., Ponomarenko, J., Clark, J.D., Cantrell, J.R., Wheeler, D.K., Gabbard, J.L., Hix, D., Sette, A. and Peters, B., 2014. The immune epitope database (IEDB) 3.0. Nucleic acids research, 43(D1), pp.D405–D412.

92. Emini, E.A., Hughes, J.V., Perlow, D., and Boger, J., 1985. Induction of hepatitis A virus-neutralizing antibody by a virus-specific synthetic peptide. Journal of Virology, 55(3), pp.836–839.

93. Karplus, P.A, and Schulz, G.E., 1985. Prediction of chain flexibility in proteins. Naturwissenschaften, 72(4), pp.212–213.

94. Kolaskar, A.S. and Tongaonkar, P.C., 1990. A semi-empirical method for prediction of antigenic determinants on protein antigens. FEBS letters, 276(1-2), pp.172–174.

95. Parker, J.M.R., Guo, D. and Hodges, R.S., 1986. New hydrophilicity scale derived from high-performance liquid chromatography peptide retention data: correlation of predicted surface residues with antigenicity and X-ray-derived accessible sites. Biochemistry, 25(19), pp.5425–5432.

96. Larsen, J.E.P., Lund, O. and Nielsen, M., 2006. An improved method for predicting linear B-cell epitopes. Immunome Research, 2(1), p.2.

97. Sievers, F. and Higgins, D.G., 2014. Clustal Omega. Current protocols in bioinformatics, 48(1), pp.3–13.

98. Khatoon, N., Pandey, R.K. and Prajapati, V.K., 2017. Exploring Leishmania secretory proteins to design B and T cell multi-epitope subunit vaccine using immunoinformatics approach. Scientific reports, 7(1), p.8285.

99. Pandey, R.K., Bhatt, T.K. and Prajapati, V.K., 2018. Novel immunoinformatics approaches to design multi-epitope subunit vaccine for malaria by investigating anopheles salivary protein. Scientific reports, 8(1), p.1125.

100. Solanki, V. and Tiwari, V., 2018. Subtractive proteomics to identify novel drug targets and reverse vaccinology for the development of a chimeric vaccine against Acinetobacter baumannii. Scientific reports, 8(1), p.9044.

101. Kim, Y., Ponomarenko, J., Zhu, Z., Tamang, D., Wang, P., Greenbaum, J., Lundegaard, C., Sette, A., Lund, O., Bourne, P.E. and Nielsen, M., 2012. Immune epitope database analysis resource. Nucleic acids research, 40(W1), pp.W525–W530.

102. Tenzer, S., Peters, B., Bulik, S., Schoor, O., Lemmel, C., Schatz, M.M., Kloetzel, P.M., Rammensee, H.G., Schild, H. and Holzhütter, H.G., 2005. Modeling the MHC class I pathway by combining predictions of proteasomal cleavage, TAP transport and MHC class I binding. Cellular and Molecular Life Sciences CMLS, 62(9), pp.1025–1037.

103. Calis, J.J., Maybeno, M., Greenbaum, J.A., Weiskopf, D., De Silva, A.D., Sette, A., Keşmir, C. and Peters, B., 2013. Properties of MHC class I presented peptides that enhance immunogenicity. PLoS computational biology, 9(10), p.e1003266.

104. Thomsen, M., Lundegaard, C., Buus, S., Lund, O. and Nielsen, M., 2013. MHCcluster, a method for functional clustering of MHC molecules. Immunogenetics, 65(9), pp.655–665. Many, A.R., Pervin, T., Mia, M., Hossain, M., Shahnaij, M., Mahmud, S. and Kibria, K.M., 2017. Vaccinomics approach for designing potential peptide vaccine by targeting Shigella spp. serine protease autotransporter subfamily protein SigA. Journal of immunology research, 2017.

105. Hasan, M., Ghosh, P.P., Azim, K.F., Mukta, S., Abir, R.A., Nahar, J. and Khan, M.M.H., 2019. Reverse vaccinology approach to design a novel multi-epitope subunit vaccine against avian influenza A (H7N9) virus. Microbial pathogenesis, 130, pp.19–37.

106. Bui, H.H., Sidney, J., Dinh, K., Southwood, S., Newman, M.J. and Sette, A., 2006. Predicting population coverage of T-cell epitope-based diagnostics and vaccines. BMC Bioinformatics, 7(1), p.153.

107. Gupta, S., Kapoor, P., Chaudhary, K., Gautam, A., Kumar, R., Raghava, G.P. and Open Source Drug Discovery Consortium, 2013. In silico approach for predicting the toxicity of peptides and proteins. PloS one, 8(9), p.e73957.

108. Jordan, K.A. and Hunter, C.A., 2010. Regulation of CD8+ T cell responses to infection with parasitic protozoa. Experimental Parasitology, 126(3), pp.318–325.

109. Moseman, E.A. and McGavern, D.B., 2013. The great balancing act: regulation and fate of antiviral T□cell interactions. Immunological reviews, 255(1), pp.110–124.

110. Honda, J., Virdi, R., and Chan, E.D., 2018. Global environmental nontuberculous mycobacteria and their contemporaneous man-made and natural niches. Frontiers in Microbiology, 9.

111. Lederman, S., Yellin, M.J., Krichevsky, A., Belko, J., Lee, J.J. and Chess, L., 1992. Identification of a novel surface protein on activated CD4+ T cells that induces contact-dependent B cell differentiation (help). Journal of Experimental Medicine, 175(4), pp.1091–1101.

112. Satyam, R., Janahi, E.M., Bhardwaj, T., Somvanshi, P., Haque, S. and Najm, M.Z., 2018. In silico identification of immunodominant B-cell and T-cell epitopes of non-structural proteins of Usutu Virus. Microbial pathogenesis, 125, pp.129–143.

113. Chauhan, V., Goyal, K. and Singh, M.P., 2018. Identification of broadly reactive epitopes targeting major glycoproteins of Herpes simplex virus (HSV) 1 and 2-An immunoinformatics analysis. Infection, Genetics, and Evolution, 61, pp.24–35.

114. Maksyutov, A., Antonets, D., Bakulina, A. and Maksyutov, R., AVAXIS BIOTHERAPEUTICS, ARTEMEV Timur and MAKSYUTOV Amir, 2013. Polyepitope constructs and methods for their preparation and use. U.S. Patent Application 13/583, 439.

115. Jung, I.D., Jeong, S.K., Lee, C.M., Noh, K.T., Heo, D.R., Shin, Y.K., Yun, C.H., Koh, W.J., Akira, S., Whang, J. and Kim, H.J., 2011. Enhanced efficacy of therapeutic cancer vaccines produced by co-treatment with Mycobacterium tuberculosis heparin-binding hemagglutinin, a novel TLR4 agonist. Cancer Research, pp.canres-3487.

116. Faridgohar, M. and Nikoueinejad, H., 2017. New findings of Toll-like receptors involved in Mycobacterium tuberculosis infection. Pathogens and global health, 111(5), pp.256–264.

117. Funderburg, N., Lederman, M.M., Feng, Z., Drage, M.G., Jadlowsky, J., Harding, C.V., Weinberg, A. and Sieg, S.F., 2007. Human β-defensin-3 activates professional antigen-presenting cells via Toll-like receptors 1 and 2. Proceedings of the National Academy of Sciences, 104(47), pp.18631–18635.

118. Bonini, C., Lee, S.P., Riddell, S.R. and Greenberg, P.D., 2001. Targeting antigen in mature dendritic cells for simultaneous stimulation of CD4+ and CD8+ T cells. The Journal of Immunology, 166(8), pp.5250–5257.

119. Saha, S. and Raghava, G.P.S., 2006. AlgPred: prediction of allergenic proteins and mapping of IgE epitopes. Nucleic acids research, 34(Suppl_2), pp.W202–W209.

120. Rana, A., and Akhter, Y., 2016. A multi-subunit based, thermodynamically stable model vaccine using combined immunoinformatics and protein structure-based approach. Immunobiology, 221(4), pp.544–557.

121. Parra, R.G., Schafer, N.P., Radusky, L.G., Tsai, M.Y., Guzovsky, A.B., Wolynes, P.G. and Ferreiro, D.U., 2016. Protein Frustratometer 2: a tool to localize energetic frustration in protein molecules, now with electrostatics. Nucleic acids research, 44(W1), pp.W356–W360.

122. Craig, D.B. and Dombkowski, A.A., 2013. Disulfide by Design 2.0: a web-based tool for disulfide engineering in proteins. BMC Bioinformatics, 14(1), p.346.

123. Schneidman-Duhovny, D., Inbar, Y., Nussinov, R. and Wolfson, H.J., 2005. PatchDock and SymmDock: servers for rigid and symmetric docking. Nucleic acids research, 33(Suppl_2), pp.W363–W367.

124. Mashiach, E., Schneidman-Duhovny, D., Andrusier, N., Nussinov, R. and Wolfson, H.J., 2008. FireDock: a web server for fast interaction refinement in molecular docking. Nucleic acids research, 36(Suppl_2), pp.W229–W232.

125. Tiwari, V., Tiwari, M. and Biswas, D., 2018. Rationale and design of an inhibitor of RecA protein as an inhibitor of Acinetobacter baumannii. The Journal of antibiotics, p.1.

126. Barh, D., Barve, N., Gupta, K., Chandra, S., Jain, N., Tiwari, S., Leon-Sicairos, N., Canizalez-Roman, A., dos Santos, A.R., Hassan, S.S. and Almeida, S., 2013. Exoproteome and secretome derived broad spectrum novel drug and vaccine candidates in Vibrio cholerae targeted by Piper betel derived compounds. PloS one, 8(1), p.e52773.

127. Yu, C.S., Lin, C.J. and Hwang, J.K., 2004. Predicting subcellular localization of proteins for Gram□negative bacteria by support vector machines based on n□peptide compositions. Protein Science, 13(5), pp.1402–1406.

128. Yu, N.Y., Wagner, J.R., Laird, M.R., Melli, G., Rey, S., Lo, R., Dao, P., Sahinalp, S.C., Ester, M., Foster, L.J. and Brinkman, F.S., 2010. PSORTb 3.0: improved protein subcellular localization prediction with refined localization subcategories and predictive capabilities for all prokaryotes. Bioinformatics, 26(13), pp.1608–1615.

129. Cheng, J., Randall, A.Z., Sweredoski, M.J. and Baldi, P., 2005. SCRATCH: a protein structure and structural feature prediction server. Nucleic acids research, 33(Suppl_2), pp.W72–W76.

130. Magnan, C.N., Randall, A. and Baldi, P., 2009. SOLpro: accurate sequence-based prediction of protein solubility. Bioinformatics, 25(17), pp.2200–2207.

131. Montomoli, E., Piccirella, S., Khadang, B., Mennitto, E., Camerini, R. and De Rosa, A., 2011. Current adjuvants and new perspectives in the vaccine formulation. Expert review of vaccines, 10(7), pp.1053–1061.

132. Kim, J., Yang, Y.L., Jang, S.H. and Jang, Y.S., 2018. Human β-defensin 2 plays a regulatory role in innate antiviral immunity and is capable of potentiating the induction of antigen-specific immunity. Virology Journal, 15(1), p.124.

133. Biragyn, A., Ruffini, P.A., Leifer, C.A., Klyushnenkova, E., Shakhov, A., Chertov, O., Shirakawa, A.K., Farber, J.M., Segal, D.M., Oppenheim, J.J. and Kwak, L.W., 2002. Toll-like receptor 4-dependent activation of dendritic cells by β-defensin 2. Science, 298(5595), pp.1025–1029.

134. Puigbo, P., Guzman, E., Romeu, A. and Garcia-Vallve, S., 2007. OPTIMIZER: a web server for optimizing the codon usage of DNA sequences. Nucleic acids research, 35(Suppl_2), pp.W126–W131.

135. Jorgensen, W.L. and Madura, J.D., 1985. Temperature and size dependence for Monte Carlo simulations of TIP4P water. Molecular Physics, 56(6), pp.1381–1392.

136. Abraham, M.J., Murtola, T., Schulz, R., Páll, S., Smith, J.C., Hess, B. and Lindahl, E., 2015. GROMACS: High performance molecular simulations through multi-level parallelism from laptops to supercomputers. SoftwareX, 1, pp.19–25.

137. Xue, L.C., Rodrigues, J.P., Kastritis, P.L., Bonvin, A.M. and Vangone, A., 2016. PRODIGY: a web server for predicting the binding affinity of protein–protein complexes. Bioinformatics, 32(23), pp.3676–3678.

138. Biotech, G., SnapGene Viewer. Glick B, editor, 3(3).

