## Supplementary section for "Towards a Chimeric Vaccine against Multiple Isolates of *Mycobacteroides* - An Integrative Approach"

**Supplementary Files**

**Figures**

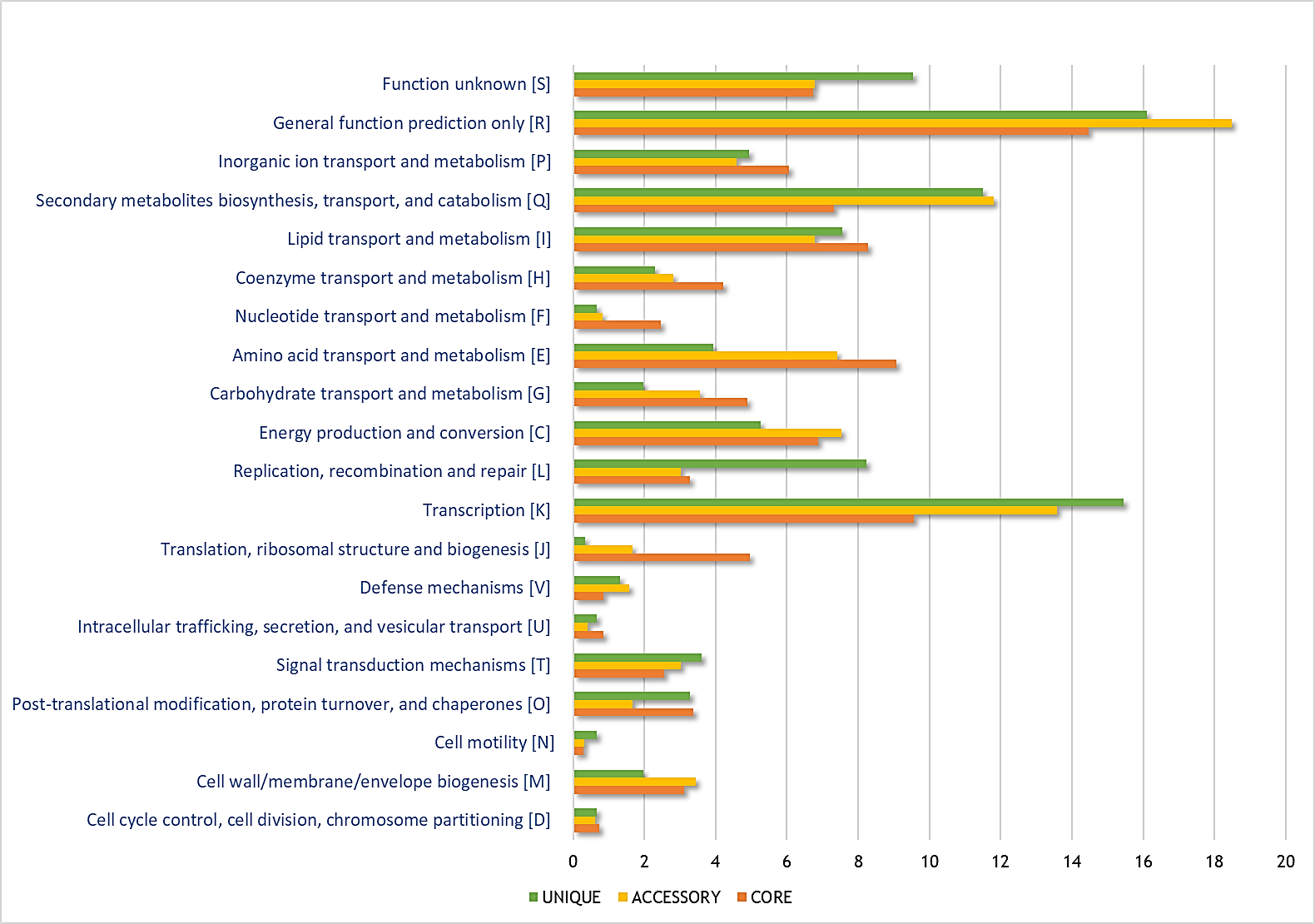
**Fig 1S:** Graphical representation of gene clustering based on COG.

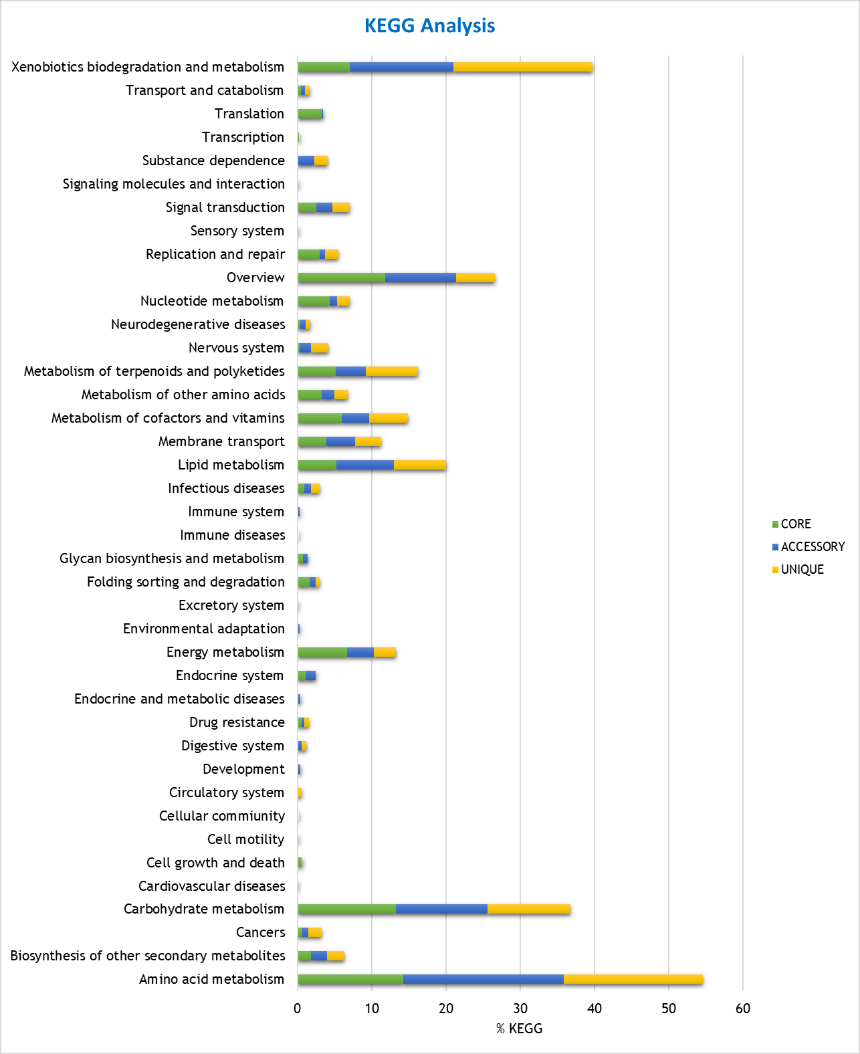
**Fig 2S:** Graphical repertory of number of genes and their functional KEGG category

**Fig 3S:** BLAST2GO pipeline used for SCL and Functional Annotation.

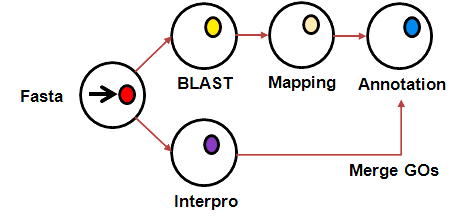

**Fig 4S:** Tertiary structure prediction, Model Refinement and Validation Pipeline.

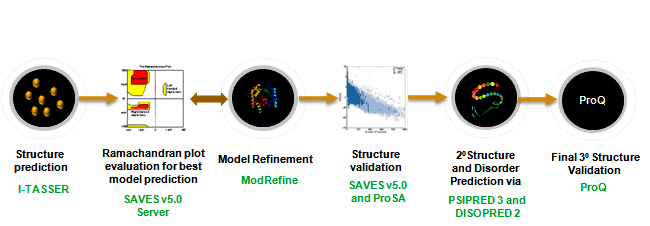

**Fig 5S:** The vaccine constructs scaffold: The net chimeric peptides were joined end to end by using EAAK (blue), GGGS (sand) and HEYGAEALERAG (Grey) linkers. We used PADRE sequence and LAMP 1 peptide in all the vaccine constructs (olive green).

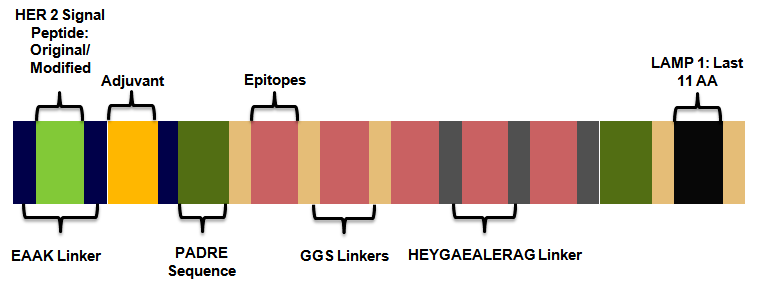

**Fig 6S:** The Macrolide resistance inducing ERM (41) gene visualized in Artemis.

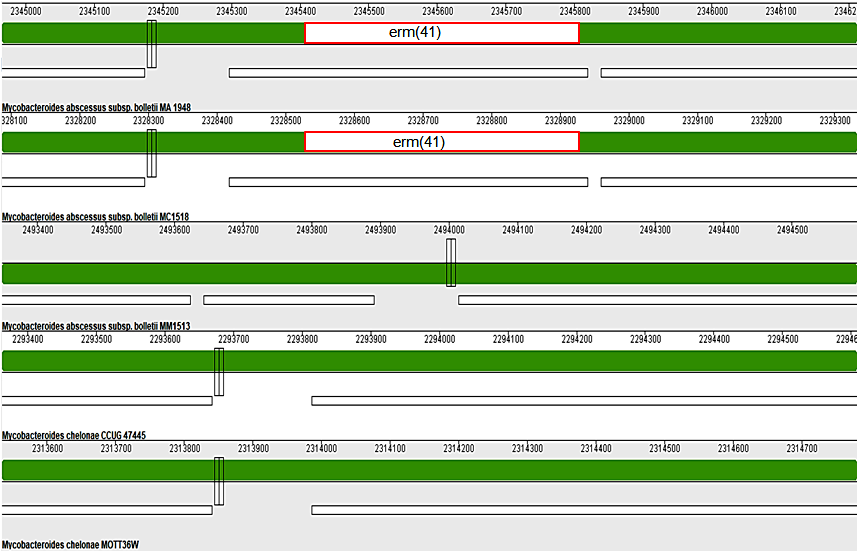

**Fig 7S:** Subcellular Localization of 391 Proteins and Virulence Prediction.

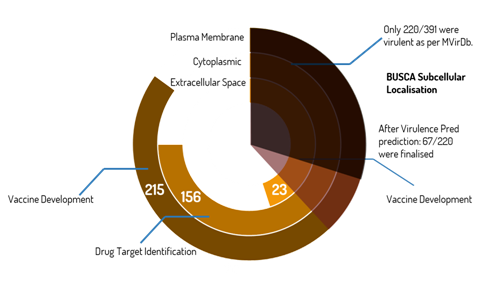

**Fig 8S:** The secondary structure prediction of (A) WP_005064885.1, (B) WP_005091736.1, (C) WP_005111658, (D) WP_012296428.1, their respective Tertiary Structure after model refinement and validation using ProSaA (Z-score plot for tertiary structures). Clearly, all the Black dots lie within the range of scores typically found for native proteins of similar size.

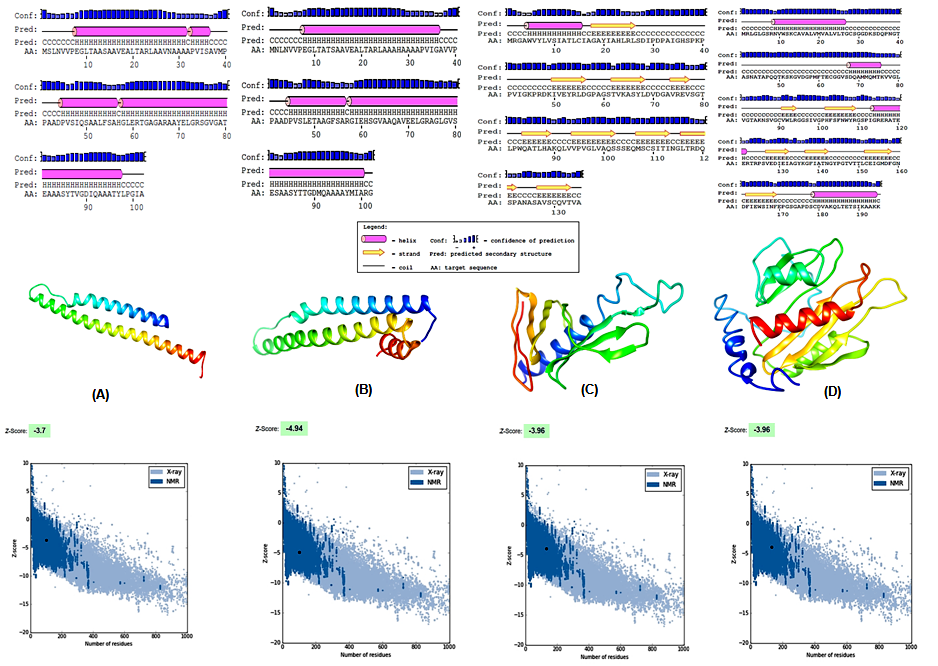

**Fig 9S:** Local Configurational Frustration region identified in (A) V6 chimeric vaccine construct and (B) V6-TLR4/MD2 complex using Frustratometer 2. The protein backbone is represented by gray ribbons and the highly frustrated regions are displayed in red, neutral contacts and minimally frustrated regions are not shown. Encircled region 1 corresponds to highly frustrated regions. Region 1 interacts with the TLR4 receptor adapter that interfaces with ligands (See B), and the region 2 highlights interaction between V6 and the cytoplasmic domain of TLR4 receptor.

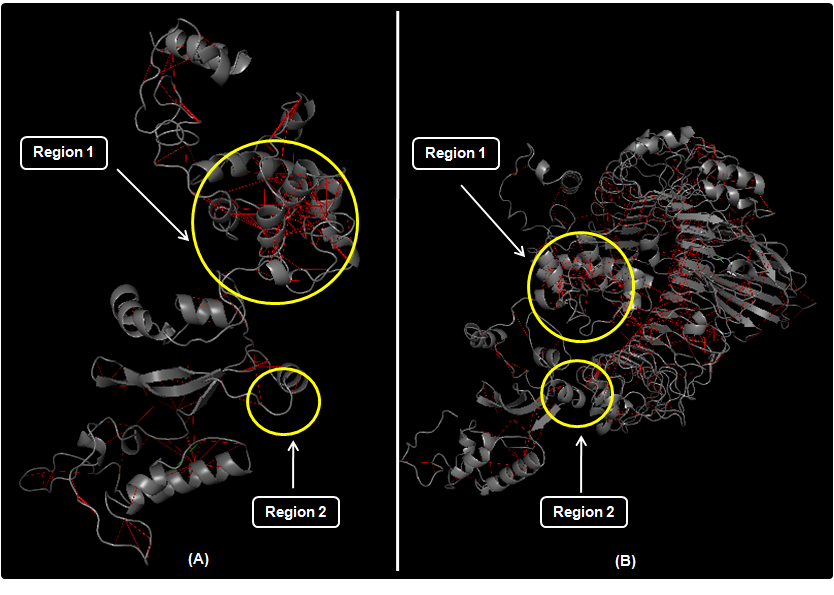

**Lists**

**1S. The panel of 27 reference set of alleles used in MHC Class I epitope prediction.**

1. A*01:01
2. A*02:01
3. A*02:03
4. A*02:06
5. A*03:01
6. A*11:01
7. A*23:01
8. A*24:02
9. A*26:01
10. A*30:01
11. A*30:02
12. A*31:01
13. A*32:01
14. A*33:01
15. A*68:01
16. A*68:02
17. B*07:02
18. B*08:01
19. B*15:01
20. B*35:01
21. B*40:01
22. B*44:02
23. B*44:03
24. B*51:01
25. B*53:01
26. B*57:01
27. B*58:01

**2S. The panel of 27 reference set of alleles used in MHC Class I epitope prediction**

2. DRB1*01:01
3. DRB1*03:01
4. DRB1*04:01
5. DRB1*04:05
6. DRB1*07:01
7. DRB1*08:02
8. DRB1*09:01
9. DRB1*11:01
10. DRB1*12:01
11. DRB1*13:02
12. DRB1*15:01
13. DRB3*01:01
14. DRB3*02:02
15. DRB4*01:01
16. DRB5*01:01
17. DPA1*01/DPB1*04:01
18. DPA1*01:03/DPB1*02:01
19. DPA1*02:01/DPB1*01:01
20. DPA1*02:01/DPB1*05:01
21. DPA1*03:01/DPB1*04:02
22. DQA1*01:01/DQB1*05:01
23. DQA1*01:02/DQB1*06:02
24. DQA1*03:01/DQB1*03:02
25. DQA1*04:01/DQB1*04:02
26. DQA1*05:01/DQB1*02:01
27. DQA1*05:01/DQB1*03:01

**3S. List of Anti Targets Used**

<https://github.com/Rohit-Satyam/Mycobacterium_Repository/blob/master/3S_List%20of%20Anti%20Targets%20Used.txt>

**4S. Broad Spectrum List**

<https://github.com/Rohit-Satyam/Mycobacterium_Repository/blob/master/4S_Broad%20Spectrum%20List.txt>

**5S. Gut Flora**

<https://github.com/Rohit-Satyam/Mycobacterium_Repository/blob/master/5S_Gut%20Flora.txt>

**6S. OrthoANI values calculated from the OAT software to decipher Genome relatedness.**

| **Genome 1** | **Genome 2** | **OrthoANI value (%)** | **Orginal ANI value (%)** | **GGDC distance** |
| --- | --- | --- | --- | --- |
| Mycobacterium abscessus subsp.  bolletii strain MA 1948 | Mycobacterium abscessus subsp.  bolletii strain MC1518 | 99.956 | 99.9438 | 5.22E-04 |
| Mycobacterium abscessus subsp.  bolletii strain MA 1948 | Mycobacterium abscessus subsp.  bolletii strain MM1513 | 97.4183 | 97.1921 | 0.027478002 |
| Mycobacterium abscessus subsp.  bolletii strain MA 1948 | Mycobacterium chelonae  CCUG 47445 | 83.695 | 83.1598 | 0.162870411 |
| Mycobacterium abscessus subsp.  bolletii strain MA 1948 | Mycobacteroides chelonae strain MOTT36W | 84.0153 | 83.5153 | 0.158279016 |
| Mycobacterium abscessus subsp.  bolletii strain MC1518 | Mycobacterium abscessus subsp.  bolletii strain MM1513 | 97.3953 | 97.218 | 0.027448845 |
| Mycobacterium abscessus subsp.  bolletii strain MC1518 | Mycobacterium chelonae CCUG 47445 | 83.5252 | 83.1768 | 0.162910161 |
| Mycobacterium abscessus subsp.  bolletii strain MC1518 | Mycobacteroides chelonae strain MOTT36W | 84.0044 | 83.5243 | 0.158464447 |
| Mycobacterium abscessus subsp.  bolletii strain MM1513 | Mycobacterium chelonae CCUG 47445 | 83.7832 | 83.2754 | 0.163576067 |
| Mycobacterium abscessus subsp. bolletii strain MM1513 | Mycobacteroides chelonae strain MOTT36W | 83.8628 | 83.3025 | 0.161695666 |
| Mycobacterium chelonae CCUG 47445 | Mycobacteroides chelonae strain MOTT36W | 95.9053 | 95.517 | 0.04171606 |

**7S. Signal peptides and Adjuvants used for Vaccine Construction**

1. Modified version of the HER2 signal peptide**: MFXAALCRWGLLLALLPPGAP:**
2. Beta Definsin**: GIINTLQKYYCRVRGGRCAVLSCLPKEEQIGKCSTRGRKCCRRKK:**
3. PADRE sequence*: AKFVAAWTLKAAA*
4. L7/L12 Ribosomal protein**: MAKLSTDELLDAFKEMTLLELSDFVKKFEETFEVTAAAPVAVAAAGAAPAGAAVEAAEEQSEFDVILEAAGDKKIGVIKVVREIVSGLGLKEAKDLVDGAPKPLLEKVAKEAADEAKAKLEAAGATVTVK**
5. HBHA adjuvant**:**

**MAENPNIDDLPAPLLAALGAADLALATVNDLIANLRERAEETRAETRTRVEERRARLTKFQEDLPEQFIELRDKFTTEELRKAAEGYLEAATNRYNELVERGEAALQRLRSQTAFEDASARAEGYVDQAVELTQEALGTVASQTRAVGERAAKLVGIEL**

6. 11 amino acids of the end of LAMP-1 protein: **RKRSHAGYQTI**
7. HBHA Conserved**: MAENSNIDDIKAPLLAALGAADLALATVNELITNLRERAEETRRSRVEESRARLTKLQEDLPEQLTELREKFTAEELRKAAEGYLEAATSELVERGEAALERLRSQQSFEEVSARAEGYVDQAVELTQEALGTVASQVEGRAAKLVGIEL**
8. Original HER2 signal peptide: **MELAALCRWGLLLALLPPGAAS**

**8S. The list of interfacial interacting residues in V6-TLR4/MD2 complex**

| Residue | Position | Chain ID (TLR4/MD2) | Residue Name | Position | Chain ID (V6) |
| --- | --- | --- | --- | --- | --- |
| TYR | 403 | B | ALA | 94 | A |
| HIS | 426 | B | ALA | 94 | A |
| HIS | 426 | B | GLY | 100 | A |
| LYS | 402 | B | PRO | 170 | A |
| GLU | 376 | B | HIS | 174 | A |
| SER | 520 | B | ARG | 277 | A |
| ASN | 544 | B | LEU | 220 | A |
| THR | 106 | B | MET | 140 | A |
| LYS | 153 | B | MET | 140 | A |
| GLU | 425 | B | PRO | 168 | A |
| ARG | 496 | B | ARG | 277 | A |
| TYR | 451 | B | GLY | 100 | A |
| ARG | 496 | B | GLU | 274 | A |
| ASN | 544 | B | SER | 279 | A |
| GLU | 425 | B | GLY | 96 | A |
| LYS | 57 | B | GLY | 47 | A |
| ARG | 355 | B | LYS | 91 | A |
| VAL | 33 | B | LEU | 126 | A |
| LYS | 130 | B | MET | 140 | A |
| SER | 518 | B | PRO | 278 | A |
| LYS | 150 | B | TRP | 14 | A |
| PRO | 142 | D | LEU | 126 | A |
| ASN | 517 | B | ARG | 277 | A |
| GLU | 79 | B | GLY | 46 | A |
| LYS | 402 | B | HIS | 174 | A |
| GLU | 474 | B | SER | 98 | A |
| CYS | 542 | B | PRO | 278 | A |
| HIS | 103 | B | GLY | 46 | A |
| LYS | 130 | B | GLY | 138 | A |
| GLU | 425 | B | PRO | 170 | A |
| SER | 141 | D | SER | 131 | A |
| GLU | 425 | B | ASP | 169 | A |
| GLU | 154 | B | ASP | 139 | A |
| LYS | 150 | B | PHE | 7 | A |
| SER | 141 | D | ALA | 133 | A |
| SER | 518 | B | ARG | 277 | A |
| SER | 102 | B | ALA | 4 | A |
| SER | 545 | B | ASN | 228 | A |
| GLU | 376 | B | LYS | 91 | A |
| ASN | 448 | B | ASP | 169 | A |
| GLU | 178 | B | GLN | 141 | A |
| GLU | 154 | B | GLY | 138 | A |
| CYS | 542 | B | ARG | 277 | A |
| GLN | 129 | B | MET | 140 | A |
| LYS | 354 | B | LYS | 91 | A |
| SER | 141 | D | ALA | 132 | A |
| ARG | 496 | B | THR | 276 | A |
| HIS | 103 | B | GLY | 47 | A |
| GLU | 425 | B | GLY | 95 | A |
| ASN | 497 | B | ASP | 166 | A |
| LYS | 72 | D | ALA | 132 | A |
| ILE | 450 | B | SER | 98 | A |
| HIS | 103 | B | ALA | 3 | A |
| HIS | 103 | B | ALA | 2 | A |
| LYS | 402 | B | ASP | 169 | A |
| SER | 126 | B | TRP | 14 | A |
| ARG | 355 | B | TRP | 88 | A |
| ASN | 544 | B | ASN | 228 | A |
| GLN | 73 | D | ALA | 133 | A |
| GLY | 97 | D | THR | 137 | A |
| GLU | 425 | B | ALA | 171 | A |
| LYS | 72 | D | THR | 136 | A |
| SER | 126 | B | PHE | 7 | A |
| SER | 100 | B | ALA | 3 | A |
| GLU | 178 | B | ASP | 139 | A |
| SER | 520 | B | ASN | 228 | A |
| GLU | 79 | B | GLY | 47 | A |
| PHE | 377 | B | LYS | 91 | A |
| GLN | 423 | B | ASP | 169 | A |
| SER | 141 | D | SER | 134 | A |
| VAL | 32 | B | LEU | 126 | A |
| ASN | 544 | B | PRO | 278 | A |
| HIS | 426 | B | GLY | 95 | A |
| SER | 102 | B | MET | 6 | A |
| SER | 126 | B | ALA | 10 | A |
| SER | 569 | B | LEU | 220 | A |
| LYS | 153 | B | ASP | 139 | A |
| HIS | 426 | B | LEU | 99 | A |
| HIS | 179 | B | ASP | 139 | A |
| HIS | 103 | B | ARG | 48 | A |
| GLY | 140 | D | ALA | 133 | A |
| GLN | 423 | B | PRO | 170 | A |
| SER | 569 | B | GLY | 219 | A |
| TYR | 451 | B | SER | 98 | A |
| LYS | 72 | D | SER | 134 | A |
| SER | 545 | B | ALA | 229 | A |
| HIS | 103 | B | MET | 6 | A |
| ILE | 450 | B | GLY | 95 | A |
| ASN | 176 | B | GLN | 141 | A |
| GLN | 73 | D | ALA | 132 | A |
| SER | 102 | B | ALA | 3 | A |
| ILE | 450 | B | GLY | 96 | A |
| ILE | 450 | B | PRO | 168 | A |
| THR | 151 | B | TRP | 14 | A |
| GLU | 154 | B | THR | 137 | A |
| TYR | 403 | B | GLY | 95 | A |
| GLU | 154 | B | MET | 140 | A |
| SER | 520 | B | ALA | 229 | A |
| LYS | 72 | D | ALA | 133 | A |
| LYS | 72 | D | THR | 137 | A |
| LYS | 153 | B | GLN | 141 | A |
| SER | 102 | B | ALA | 2 | A |
| VAL | 475 | B | SER | 98 | A |
| LYS | 130 | B | THR | 137 | A |
| LYS | 57 | B | ARG | 48 | A |
| PRO | 142 | D | SER | 131 | A |
| SER | 570 | B | LEU | 220 | A |
| SER | 520 | B | PRO | 278 | A |
| GLU | 79 | B | ARG | 48 | A |

**9S. Vaccine Sequence with adapter sequence in Bold**

**AGCT** GAGGCTGCCGCTAAAATGTTCTGTGCAGCTCTGTGCCGGTGGGGACTGCTCCTGGCCCTCCTGCCACCTGGAGCTCCGGAAGCGGCAGCAAAAGGCATAATCAATACGCTGCAAAAATACTATTGTCGTGTCCGTGGAGGACGTTGTGCAGTTCTCAGCTGTCTGCCGAAAGAGGAACAGATAGGAAAGTGTAGTACTCGTGGACGAAAGTGCTGCCGTAGGAAAAAGGAGGCTGCTGCCAAAGCTAAATTTGTCGCGGCCTGGACCTTGAAAGCGGCGGCTGGCGGAGGATCCTTGGGAAGGAGCGGTGTTGGTGCTACAGAGGCAGCGGCCAGCTATACTGTAGGAGACATCCAAGCTGCAGGAGGAGGCTCGTTGGGAGTATCGGAAAGTGCAGCAAGTTACACGACAGGCGATATGCAGGCGGCTGCTGCCTATATGATAGCGCGGGGAGGCGGTGGCTCGGGAGCATATATCGCCCATTTGAGGCTGAGTGATATCCCCGACCCGGCAATTGGCCACAGCCCCAAGCCCCCTGTAATAGGTAAGCCTAGGGACAAAATAGTCGAGTACAGGCTGGACGGATGCGCAGGCTGTACTGTCAAAGCAAGCTATCTGGACCACGAGTATGGATGTGAATGCCTGGAACGTGCCGGCCTGACGCGAGACCAGAGTCCCGCCAACGCCTCGGCTGTATCCTGCCAGGTCACCGTGGCCCATGAATATGGAGCAGAAGCGCTGGAGCGAGCGGGCGTAGGTCCACATTTCAGTTTCAACTGGTACCGAGGCTCCCCCATTGGACGAGAAAGGGCGACCGAAGAGCGCACAAGGCCTTCCGTCGAAGACATAGAGATTGCGGGCCATGAATACGGAGCGGAAGCTCTCGAGCGAGCTGGAATTGAGTGGTCGATTAACTTCGAACCGGGAAGTGGCGCACCTGACAGTTGCGACGTAGCACATGAGTATGGTGCCGAAGCATTGGAACGGGCGGGCGCGAAGTGCTGCGCAGCTTGGACGTTGAAAGCGGCTGCAGGCGGAGGTAGTCGTAAAAGGTCGCACGCAGGTTATCAAACGATCGGCGGTGGCAGC

**AGCT**

**Tables**

**Table 1S: List of residues predicted to be Conformational B-Cell antigenic determinants. The score represents the confidence of the prediction made by the ElliPro server.**

| Protein Accession Number | Discontinuous epitopes residues | Number of residues | Score |
| --- | --- | --- | --- |
| WP_005064885 | A:V88, A:I91, A:Q92, A:A93, A:A94, A:A95, A:T96, A:L98, A:P99, A:G100, A:I101, A:A102 | 12 | 0.808 |
|  | A:M1, A:S2, A:L3, A:N4, A:V5, A:V6, A:E8, A:G9, A:A12 | 9 | 0.782 |
|  | A:V38, A:M39, A:P40, A:P41, A:A42, A:A43, A:D44, A:P45, A:V46, A:S47, A:I48, A:Q49, A:S50, A:A52, A:L53, A:A56 | 16 | 0.735 |
|  | A:A30, A:P33, A:V34 | 3 | 0.588 |
| WP_005091736.1 | A:A30, A:P33, A:G36, A:A37, A:V38, A:V39, A:P40, A:P41, A:A42, A:A43, A:D44, A:P45, A:V46, A:S47, A:L48, A:E49, A:T50 | 17 | 0.777 |
|  | A:M1, A:N2, A:L3, A:N4, A:V5, A:V6, A:P7, A:E8, A:E81, A:A83, A:A84, A:S85, A:Y86, A:T87, A:T88, A:G89, A:D90, A:Q92, A:A93, A:A96, A:Y97, A:I99, A:A100, A:R101, A:G102 | 25 | 0.722 |
|  | A:T11, A:A12, A:A15 | 3 | 0.588 |
| WP_005111658.1 | _:T86, _:L87, _:H88, _:A89, _:K90, _:Q91, _:L92, _:V93, _:V94, _:P95, _:S110, _:N114, _:G115, _:L116, _:T117, _:V132, _:T133, _:V134, _:A135 | 19 | 0.657 |
|  | _:H23, _:L26, _:S27, _:D28, _:I29, _:P30, _:D31, _:P32, _:A33, _:I34, _:G35, _:H36, _:S37, _:P38, _:K39, _:P40, _:P41, _:V42, _:I43, _:G44, _:K45, _:P46, _:R47, _:D48, _:K49, _:Y67, _:L68, _:D69, _:V70, _:D71, _:G72, _:A73, _:V74, _:R75 | 34 | 0.647 |
|  | _:M1, _:R2, _:G3, _:A4, _:W5, _:V6, _:Y7, _:D56, _:G57, _:P58, _:A59, _:G60, _:S61, _:T62, _:A100, _:Q101, _:S102, _:S103, _:S104, _:E105, _:Q106, _:M107, _:S108, _:S121, _:P122, _:A123, _:N124, _:A125, _:S126, _:A127, _:V128, _:S129 | 32 | 0.623 |
| WP_012296428.1 | A:T141, A:N142, A:G143, A:Y144, A:P145, A:G146, A:T147, A:V148, A:T149, A:T150, A:L151 | 11 | 0.772 |
|  | A:F169, A:E170, A:P171, A:G172, A:S173, A:G174, A:A175, A:P176, A:D177, A:S178 | 10 | 0.705 |
|  | A: R94, A: G95, A: G96, A: S97, A: I98, A: V99 | 6 | 0.703 |
|  | A:P47, A:Q49, A:T50, A:K51, A:S52, A:K53, A:G54, A:V55, A:D56, A:G57, A:P58, A:M59, A:Q69, A:M72, A:Q73, A:T75, A:K76, A:V77, A:V78, A:G79, A:L93 | 21 | 0.642 |
|  | A:F60, A:T61, A:E62 | 3 | 0.632 |
|  | A:M1, A:R2, A:L3, A:G4, A:L5, A:G6, A:S7, A:R8, A:N9, A:V10, A:W11, A:S12, A:K13, A:C14, A:A15, A:V16, A:A17, A:L18, A:V19, A:M20, A:V21, A:A22, A:L23, A:V24, A:L25, A:T26, A:G27, A:C28, A:S29, A:G30, A:G31, A:D32, A:K33, A:S34, A:D35, A:Q36, A:P37, A:N38, A:G39, A:T40, A:A41, A:S42, A:N43, A:A44, A:T45, A:A46, A:N86, A:Y108, A:R109, A:G110, A:S111, A:P112, A:I113, A:G114, A:R115, A:A118, A:E121, A:R122, A:R124, A:P125, A:S126, A:V127, A:E128, A:D129, A:I130, A:E131, A:I132, A:A133, A:G134, A:Y135, A:K136, A:A140, A:G159, A:K194 | 74 | 0.604 |

**Table 2S: The final B-Cell, MHC Class I and II epitopes used for Net chimeric peptide construction and their respective HLA alleles.**

| Protein Accession No. | Final B Cell  Epitope | MHC I | Peptide start | Peptide end | HLA alleles | MHC II | Peptide start | Peptide end | HLA Alleles |
| --- | --- | --- | --- | --- | --- | --- | --- | --- | --- |
| WP_005064885 | **NAAAAPVISAVMPPAADPV**SIQSAAL | LAAV**NAAAA** | 24 | 32 | HLA-B*35:01 | **APVISAVMPPAADPV** | 32 | 46 | HLA-DQA1*05:01/  DQB1*03:01 |
|  |  |  |  |  |  | **PPAADPVSIQSAAL**F | 40 | 54 | HLA-DQA1*05:01/  DQB1*02:01 |
|  | LGRSGVGATEAAAS**YTVGDIQA** | **YTVGDIQA**A | 86 | 94 | HLA-A*26:01 |  |  |  |  |
|  | **FSAHGLERTGAGARAA** | **TGAGARAA**Y | 62 | 70 | HLA-B*35:01 | PVSIQSAAL**FSAHGL** | 45 | 59 | HLA-DQA1*05:01/  DQB1*03:01 |
| WP_005091736.1 | A**AAAPVIG**AVVPPAADPVSLETAA | H**AAAAPVIG** | 28 | 36 | HLA-B*35:01 | VEALTARLAAA**HAAA** | 17 | 31 | HLA-DQA1*05:01,  HLA-DQB1*03:01 |
|  |  |  |  |  |  | ARLAAAHA**AAAPVIG** | 22 | 36 | HLA-DQA1*05:01,  HLA-DQB1*03:01 |
|  | LGVSESAASYTTGDMQ**AAAA** | **AAAA**YMIAR | 93 | 101 | HLA-A*11:01 | **TGDMQAAAA**YMIARG | 88 | 102 | HLA-DQA1*05:01,  HLA-DQB1*03:01 |
|  |  | **MQAAAAYMI** | 91 | 99 | HLA-B*51:01 |  |  |  |  |
| WP_005111658.1 | **LSDIPDPA**IGHSPKPPVIGKPRDK | R**LSDIPDPA** | 25 | 33 | HLA-A*02:03 | GAYIAHLR**LSDIPDP** | 18 | 32 | HLA-DQA1*05:01/  DQB1*02:01 |
|  | KIVEYRLDGPAGSTVKASYLD |  |  |  |  |  |  |  |  |
|  | HAKQLVVPPVGLVAQSSSEQMSC |  |  |  |  | **HAKQLVVPPVGLVA** | 88 | 101 | HLA-DPA1*01:03/  DPB1*02:01 |
|  | LTRDQSPANASAVSCQVTVA |  |  |  |  |  |  |  |  |
| WP_012296428.1 | MRLGLGSRNVWSKC**AVAL**  **VMVALVLTGCS**GGDKSDQ  PNGTASNATAPQQTKSKG  VDGPMFTECGGVSD |  |  |  |  | **AVALVMVALVLTGCS** | 15 | 29 | HLA-DPA1*01:03,  HLA-DPB1*02:01 |
|  |  |  |  |  |  | VWSKCAVALVMVALV | 10 | 24 | HLA-DQA1*05:01,HLA-DQB1*03:01 |
|  | **YRGSPIGRERAT**EERTRPSVEDIEIAG | **SPIGRERAT** | 111 | 119 | HLA-B*07:02 | VGPHFSFNW**YRGSPI** | 99 | 113 | HLA-DPA1*01:03,HLA-DPB1*02:01 |
|  |  | FNW**YRGSPI** | 105 | 113 | HLA-B*08:01 |  |  |  |  |
|  | IEWSINFEPGSGAPDSCDVA |  |  |  |  |  |  |  |  |
|  | KGFIATNGYPGTVTTLCEIG | **GYPGTVTTL** | 143 | 151 | HLA-A*24:02 |  |  |  |  |

**Table 3S: The Population Coverage analysis of Class I and II antigenic determinants used in vaccine construction, Where: ^a^ average number of epitope hits/ HLA combinations recognized by the population, ^b^ minimum number of epitope hits/ HLA combinations recognized by 90% of the population**

| **Population/area** | **Class I Antigenic Determinants (enlisted in Table 2)** | | | **Class II Antigenic Determinants (enlisted in Table 2)** | | |
| --- | --- | --- | --- | --- | --- | --- |
|  | **Projected Population Coverage** | **Average Hit^a^** | **pc90^b^** | **Projected Population Coverage** | **Average Hit^a^** | **pc90^b^** |
| **Australia** | **70.39%** | **1.0** | **0.34** | **60.1%** | **4.09** | **0.5** |
| **Central Africa** | **43.02%** | **0.73** | **0.18** | **89.48%** | **6.72** | **1.9** |
| **China** | **80.16%** | **1.28** | **0.5** | **91.67%** | **7.86** | **2.69** |
| **East Africa** | **42.0%** | **0.59** | **0.17** | **89.13%** | **7.52** | **1.84** |
| **East Asia** | **89.99%** | **1.76** | **1.0** | **67.74%** | **4.12** | **0.62** |
| **India** | **68.61%** | **1.09** | **0.32** | **95.92%** | **7.63** | **3.3** |
| **Japan** | **93.15%** | **1.92** | **1.08** | **83.71%** | **5.64** | **1.23** |
| **North Africa** | **56.56%** | **0.96** | **0.23** | **84.03%** | **6.4** | **1.25** |
| **Singapore** | **79.45%** | **1.31** | **0.49** | **72.9%** | **4.09** | **0.74** |
| **West Africa** | **62.14%** | **1.21** | **0.26** | **92.43%** | **10.31** | **4.15** |
| **World** | **71.42%** | **1.21** | **0.35** | **92.47%** | **7.95** | **2.85** |
| **Average** | **68.19** | **1.17** | **0.43** | **83.6** | **6.58** | **1.92** |

**Table 4S: Physicochemical characterization of 6 non-allergic vaccine constructs.**

| **Vaccine Construct** | **EP-class** | **RF-class** | **Length** | **Gravy index** | **Instability index (II)** | **Isoelectric point (pI)** | **Coiled coils** | **Longest disorder region** | **Percentage of coil structure** | **Transmembrane helices (TM)** | **Signal peptides (SP)** | **Insertions score** |
| --- | --- | --- | --- | --- | --- | --- | --- | --- | --- | --- | --- | --- |
| [V1](http://ffas.burnham.org/XtalPred-cgi/result.pl?dir=webdb/1541081585.21646/0) | 5 | 11 | 542 | -0.18 | 37.67 | 5.79 | 85 | 60 | 37 | No | 25 | 0.09 |
| [V2](http://ffas.burnham.org/XtalPred-cgi/result.pl?dir=webdb/1541081585.21646/1) | 5 | 11 | 445 | -0.05 | 44.49 | 4.99 | 43 | 19 | 31 | No | 26 | 0.14 |
| [V3](http://ffas.burnham.org/XtalPred-cgi/result.pl?dir=webdb/1541081585.21646/2) | 5 | 5 | 384 | -0.13 | 35.24 | 8.65 | 0 | 20 | 50 | No | 25 | 0.04 |
| [V4](http://ffas.burnham.org/XtalPred-cgi/result.pl?dir=webdb/1541081585.21646/4) | 5 | 11 | 512 | -0.27 | 36.92 | 5.67 | 85 | 49 | 35 | No | 25 | 0.08 |
| [V5](http://ffas.burnham.org/XtalPred-cgi/result.pl?dir=webdb/1541081585.21646/5) | 5 | 11 | 415 | -0.15 | 44.05 | 4.91 | 43 | 53 | 27 | No | 26 | 0.14 |
| [V6](http://ffas.burnham.org/XtalPred-cgi/result.pl?dir=webdb/1541081585.21646/6) | 5 | 5 | 354 | -0.26 | 33.95 | 8.64 | 0 | 20 | 50 | No | 25 | 0.05 |

**Table 5S: Docking of 6 vaccine constructs with HLA molecules (Both Class I and II) using Patchdock and Pose refined by Firedock. Above tabulated are Binding energies of refined poses from Firedock.**

| **Vaccine Constructs** | **HLA alleles PDB ID’s** | **Receptor** | **Global Energy/  Binding energy** | **Attractive  VdW** | **Repulsive  VdW** | **ACE** | **HB** |
| --- | --- | --- | --- | --- | --- | --- | --- |
| **V1** | HLA-DR B1*03:01 | 1A6A | -4.17 | -6.15 | 4.55 | -1.30 | 0.00 |
|  | HLA-DR B3*02:02 | 3C5J | -37.95 | -36.73 | 23.41 | 4.38 | -2.87 |
|  | HLA-DR B5*01:01 | 1H15 | -11.41 | -28.76 | 11.01 | -0.32 | -2.24 |
|  | HLA-B*35-01 | 2H6P | 3.38 | -8.01 | 1.71 | 8.45 | -2.54 |
|  | HLA-A*11:01 | 2HN7 | 1.01 | -4.27 | 0.09 | 2.20 | 0.00 |
|  | HLA-B*51:01 | 1E28 | -2.86 | -25.94 | 18.44 | 9.62 | -5.45 |
|  | HLA-A*24:02 | 2BCK | -10.58 | -19.92 | 5.60 | 1.79 | -1.92 |
| **V2** | HLA-DR B1*03:01 | 1A6A | -10.95 | -33.89 | 18.57 | 16.19 | -7.79 |
|  | HLA-DR B3*02:02 | 3C5J | -1.86 | -10.09 | 10.78 | 1.05 | -0.43 |
|  | HLA-DR B5*01:01 | 1H15 | -45.94 | -40.78 | 17.76 | 4.74 | -4.27 |
|  | HLA-B*35-01 | 2H6P | -7.45 | -15.43 | 8.77 | 8.27 | -1.29 |
|  | HLA-A*11:01 | 2HN7 | 0.36 | -35.57 | 19.62 | 14.02 | -5.09 |
|  | HLA-B*51:01 | 1E28 | -8.58 | -26.91 | 8.01 | 6.31 | -1.9 |
|  | HLA-A*24:02 | 2BCK | -14.95 | -10.54 | 1.29 | -2.06 | -0.84 |
| **V3** | HLA-DR B1*03:01 | 1A6A | -11.00 | -35.21 | 47.95 | 10.67 | -4.50 |
|  | HLA-DR B3*02:02 | 3C5J | -16.81 | -37.60 | 12.13 | 4.85 | -3.61 |
|  | HLA-DR B5*01:01 | 1H15 | -1.60 | -8.22 | 0.90 | -1.17 | 0 |
|  | HLA-B*35-01 | 2H6P | 0.22 | -29.96 | 30.39 | 8.66 | -1.48 |
|  | HLA-A*11:01 | 2HN7 | -1.36 | -23.39 | 10.67 | 5.81 | -2.96 |
|  | HLA-B*51:01 | 1E28 | -19.28 | -26.75 | 11.90 | 3.93 | -3.41 |
|  | HLA-A*24:02 | 2BCK | -6.19 | -41.10 | 38.59 | 5.03 | -3.99 |
| **V4** | HLA-DR B1*03:01 | 1A6A | -14.48 | -36.81 | 21.32 | 8.31 | -3.78 |
|  | HLA-DR B3*02:02 | 3C5J | -2.33 | -36.63 | 27.49 | 12.99 | -5.68 |
|  | HLA-DR B5*01:01 | 1H15 | 2.15 | -1.36 | 0.00 | -0.09 | 0.00 |
|  | HLA-B*35-01 | 2H6P | -18.51 | -27.27 | 7.91 | 11.08 | -4.17 |
|  | HLA-A*11:01 | 2HN7 | -23.65 | -37.63 | 28.89 | -0.34 | -6.12 |
|  | HLA-B*51:01 | 1E28 | -0.35 | -17.78 | 10.11 | 6.97 | -2.63 |
|  | HLA-A*24:02 | 2BCK | 6.56 | -2.65 | 1.61 | 2.27 | -0.80 |
| **V5** | HLA-DR B1*03:01 | 1A6A | -32.67 | -34.38 | 35.96 | 0.69 | -2.37 |
|  | HLA-DR B3*02:02 | 3C5J | -5.05 | -30.38 | 28.75 | 13.36 | -3.19 |
|  | HLA-DR B5*01:01 | 1H15 | -5.80 | -21.53 | 17.00 | 2.22 | -2.87 |
|  | HLA-B*35-01 | 2H6P | -15.80 | -37.78 | 23.23 | 4.38 | -2.4 |
|  | HLA-A*11:01 | 2HN7 | -10.73 | -28.92 | 17.92 | 9.00 | -0.30 |
|  | HLA-B*51:01 | 1E28 | 3.58 | -25.09 | 15.58 | 14.18 | -3.92 |
|  | HLA-A*24:02 | 2BCK | -18.01 | -22.46 | 9.73 | 2.16 | -3.82 |
| **V6** | HLA-DR B1*03:01 | 1A6A | -17.15 | -37.83 | 26.97 | 14.94 | -7.87 |
|  | HLA-DR B3*02:02 | 3C5J | -23.86 | -50.52 | 19.11 | 11.15 | -5.23 |
|  | HLA-DR B5*01:01 | 1H15 | -7.07 | -27.72 | 27.79 | -1.81 | -1.28 |
|  | HLA-B*35-01 | 2H6P | -12.64 | -22.97 | 15.28 | 1.53 | -4.49 |
|  | HLA-A*11:01 | 2HN7 | -25.31 | -19.78 | 15.24 | -3.90 | -1.32 |
|  | HLA-B*51:01 | 1E28 | -20.79 | -31.89 | 17.09 | 13.02 | -3.10 |
|  | HLA-A*24:02 | 2BCK | -20.18 | -21.09 | 7.18 | -0.04 | -0.92 |

**Table 6S: Net chimeric peptide constructs generated in Phase C for vaccine construction. The eight vaccine constructs V1, V2, V3, V4, V5, V6, V7, V8 were made using the peptides the neighboring cells of which are colored black**

| **Accession No.**  **of Proteins** | **Net Peptides for vaccine construct** | **Position** | **V1** | **V2** | **V3** | **V4** | **V5** | **V6** | **V7** | **V8** |
| --- | --- | --- | --- | --- | --- | --- | --- | --- | --- | --- |
| WP_005064885 | NAAAAPVISAVMPPAADPVSIQSAALFSAHGLERTGAGARAAY | 28-70 |  |  |  |  |  | | | |
|  | LGRSGVGATEAAASYTVGDIQAA | 72-94 |  | | | |  |  |  |  |
| WP_005091736.1 | VEALTARLAAAHAAAAPVIGAVVPPAADPVSLETAA | 17-52 |  |  |  |  |  | | | |
|  | LGVSESAASYTTGDMQAAAAYMIARG | 77-102 |  | | | |  |  |  |  |
| WP_005111658.1 | GAYIAHLRLSDIPDPAIGHSPKPPVIGKPRDKIVEYRLDGPAGSTVKASYLD | 18-69 |  |  |  |  |  |  |  |  |
|  | HAKQLVVPPVGLVAQSSSEQMSC | 88-109 |  |  |  | |  |  |  | |
|  | LTRDQSPANASAVSCQVTVA | 116-135 |  |  |  |  |  |  |  |  |
| WP_012296428.1 | MRLGLGSRNVWSKCAVALVMVALVLTGCSGGDKSDQPNGTASNATAP | 0-68 |  |  |  |  |  |  |  |  |
|  | QQTKSKGVDGPMFT |  |  |  |  |  |  |  |  |  |
|  | VGPHFSFNWYRGSPIGRERATEERTRPSVEDIEIAG | 99-134 |  |  |  |  |  |  |  |  |
|  | IEWSINFEPGSGAPDSCDVA | 163-182 |  |  |  |  |  |  |  |  |
|  | KGFIATNGYPGTVTTLCEIG | 136-155 |  |  |  |  |  |  |  |  |

**Table 7S: Multi-epitope vaccine constructs and their prediction of antigenicity and allergenicity using VaxiJen and Algpred Servers, respectively.**

| **Final 6 Vaccine construct** | **Vaccine Construct Sequences** | **Antigenicity Score  (AntigenPRO)** | **Solubility upon  Over-expression >=0.5** | **Allergenicity (Algpred)  SVM Score (Treshold-0.4)** | **SVM dipeptide  score Threshold= -0.2** | **Vaxijen** |
| --- | --- | --- | --- | --- | --- | --- |
| **V1** | EAAAKMFCAALCRWGLLLALLPPGAPEAAAKMAENPNIDDLPAPLLAALGAADLALATVNDLIANLRERAEETRAETRTRVEERRARLTKFQEDLPEQFIELRDKFTTEELRKAAEGYLEAATNRYNELVERGEAALQRLRSQTAFEDASARAEGYVDQAVELTQEALGTVASQTRAVGERAAKLVGIELEAAAK*AKFVAAWTLKAAA*GGGSNAAAAPVISAVMPPAADPVSIQSAALFSAHGLERTGAGARAAYGGGSVEALTARLAAAHAAAAPVIGAVVPPAADPVSLETAAGGGSGAYIAHLRLSDIPDPAIGHSPKPPVIGKPRDKIVEYRLDGPAGSTVKASYLDHEYGAEALERAGHAKQLVVPPVGLVAQSSSEQMSCHEYGAEALERAGMRLGLGSRNVWSKCAVALVMVALVLTGCSGGDKSDQPNGTASNATAPQQTKSKGVDGPMFTHEYGAEALERAGVGPHFSFNWYRGSPIGRERATEERTRPSVEDIEIAGHEYGAEALERAG*AKFVAAWTLKAAA*GGGSRKRSHAGYQTIGGGS | 0.422791 | 0.862684 | -1.4061132 | -0.64664231 | 0.8028 |
| **V2** | EAAAKMELAALCRWGLLLALLPPGAASEAAAKMAENSNIDDIKAPLLAALGAADLALATVNELITNLRERAEETRRSRVEESRARLTKLQEDLPEQLTELREKFTAEELRKAAEGYLEAATSELVERGEAALERLRSQQSFEEVSARAEGYVDQAVELTQEALGTVASQVEGRAAKLVGIELEAAAK*AKFVAAWTLKAAA*GGGSNAAAAPVISAVMPPAADPVSIQSAALFSAHGLERTGAGARAAYGGGSVEALTARLAAAHAAAAPVIGAVVPPAADPVSLETAAGGGSHAKQLVVPPVGLVAQSSSEQMSCHEYGAEALERAGLTRDQSPANASAVSCQVTVAHEYGAEALERAGIEWSINFEPGSGAPDSCDVAHEYGAEALERAGKGFIATNGYPGTVTTLCEIGHEYGAEALERAG*AKFVAAWTLKAAA*GGGSRKRSHAGYQTIGGGS | 0.552431 | 0.913939 | -0.75967072 | -0.42483292 | 0.7523 |
| **V3** | EAAAKMFCAALCRWGLLLALLPPGAPEAAAKGIINTLQKYYCRVRGGRCAVLSCLPKEEQIGKCSTRGRKCCRRKKEAAAK*AKFVAAWTLKAAA*GGGSNAAAAPVISAVMPPAADPVSIQSAALFSAHGLERTGAGARAAYGGGSVEALTARLAAAHAAAAPVIGAVVPPAADPVSLETAAGGGSGAYIAHLRLSDIPDPAIGHSPKPPVIGKPRDKIVEYRLDGPAGSTVKASYLDHEYGAEALERAGLTRDQSPANASAVSCQVTVAHEYGAEALERAGVGPHFSFNWYRGSPIGRERATEERTRPSVEDIEIAGHEYGAEALERAGIEWSINFEPGSGAPDSCDVAHEYGAEALERAG*AKFVAAWTLKAAA*GGGSRKRSHAGYQTIGGGS | 0.471663 | 0.856609 | -1.2677549 | -0.54703438 | 0.8903 |
| **V4** | EAAAKMFCAALCRWGLLLALLPPGAPEAAAKMAENPNIDDLPAPLLAALGAADLALATVNDLIANLRERAEETRAETRTRVEERRARLTKFQEDLPEQFIELRDKFTTEELRKAAEGYLEAATNRYNELVERGEAALQRLRSQTAFEDASARAEGYVDQAVELTQEALGTVASQTRAVGERAAKLVGIELEAAAK*AKFVAAWTLKAAA*GGGSLGRSGVGATEAAASYTVGDIQAAGGGSLGVSESAASYTTGDMQAAAAYMIARGGGGSGAYIAHLRLSDIPDPAIGHSPKPPVIGKPRDKIVEYRLDGPAGSTVKASYLDHEYGAEALERAGHAKQLVVPPVGLVAQSSSEQMSCHEYGAEALERAGMRLGLGSRNVWSKCAVALVMVALVLTGCSGGDKSDQPNGTASNATAPQQTKSKGVDGPMFTHEYGAEALERAGVGPHFSFNWYRGSPIGRERATEERTRPSVEDIEIAGHEYGAEALERAG*AKFVAAWTLKAAA*GGGSRKRSHAGYQTIGGGS | 0.706497 | 0.915646 | -1.3858001 | -0.6247704 | 0.842 |
| **V5** | EAAAKMELAALCRWGLLLALLPPGAASEAAAKMAENSNIDDIKAPLLAALGAADLALATVNELITNLRERAEETRRSRVEESRARLTKLQEDLPEQLTELREKFTAEELRKAAEGYLEAATSELVERGEAALERLRSQQSFEEVSARAEGYVDQAVELTQEALGTVASQVEGRAAKLVGIELEAAAK*AKFVAAWTLKAAA*GGGSLGRSGVGATEAAASYTVGDIQAAGGGSLGVSESAASYTTGDMQAAAAYMIARGGGGSHAKQLVVPPVGLVAQSSSEQMSCHEYGAEALERAGLTRDQSPANASAVSCQVTVAHEYGAEALERAGIEWSINFEPGSGAPDSCDVAHEYGAEALERAGKGFIATNGYPGTVTTLCEIGHEYGAEALERAG*AKFVAAWTLKAAA*GGGSRKRSHAGYQTIGGGS | 0.706497 | 0.915646 | -0.68331163 | -0.39519952 | 0.7946 |
| **V6** | EAAAKMFCAALCRWGLLLALLPPGAPEAAAKGIINTLQKYYCRVRGGRCAVLSCLPKEEQIGKCSTRGRKCCRRKKEAAAK*AKFVAAWTLKAAA*GGGSLGRSGVGATEAAASYTVGDIQAAGGGSLGVSESAASYTTGDMQAAAAYMIARGGGGSGAYIAHLRLSDIPDPAIGHSPKPPVIGKPRDKIVEYRLDGPAGSTVKASYLDHEYGAEALERAGLTRDQSPANASAVSCQVTVAHEYGAEALERAGVGPHFSFNWYRGSPIGRERATEERTRPSVEDIEIAGHEYGAEALERAGIEWSINFEPGSGAPDSCDVAHEYGAEALERAG*AKFVAAWTLKAAA*GGGSRKRSHAGYQTIGGGS | 0.645761 | 0.869523 | -1.2466774 | -0.53615302 | 0.957 |
